# A vaccine antigen central in influenza A(H5) virus antigenic space confers subtype-wide immunity

**DOI:** 10.1101/2024.08.06.606696

**Authors:** Adinda Kok, Samuel H. Wilks, Sina Tureli, Sarah L. James, Theo M. Bestebroer, David F. Burke, Mathis Funk, Stefan van der Vliet, Monique I. Spronken, Willemijn F. Rijnink, David Pattinson, Dennis de Meulder, Miruna E. Rosu, Pascal Lexmond, Judith M.A. van den Brand, Sander Herfst, Derek J. Smith, Ron A.M. Fouchier, Mathilde Richard

## Abstract

Highly pathogenic avian influenza A(H5) viruses globally impact wild and domestic birds, and mammals, including humans, underscoring their pandemic potential. The antigenic evolution of the A(H5) hemagglutinin (HA) poses challenges for pandemic preparedness and vaccine design. Here, the global antigenic evolution of the A(H5) HA was captured in a high-resolution antigenic map. The map was used to engineer immunogenic and antigenically central vaccine HA antigens, eliciting antibody responses that broadly cover the A(H5) antigenic space. In ferrets, a central antigen protected as well as homologous vaccines against heterologous infection with two antigenically distinct viruses. This work showcases the rational design of subtype-wide influenza A(H5) pre-pandemic vaccines and demonstrates the value of antigenic maps for the evaluation of vaccine-induced immune responses through antibody profiles.

## Main Text

Influenza A viruses are enzootic in wild migratory birds of aquatic habitats around the world (*1*). In wild waterfowl, 17 subtypes of hemagglutinin (HA, H1-H16, H19) and 9 subtypes of neuraminidase (NA, N1-N9), surface glycoproteins of influenza A viruses, have been identified (*2, 3*). The HA is the receptor-binding and fusion protein of influenza A viruses, and its head domain (HA1) contains dominant epitopes targeted by (neutralizing) antibodies. The host cell receptors of avian and human influenza viruses are respectively α2,3-linked and α2,6-linked sialic acids, and receptor binding specificity is one of the major host range determinants (*2*). Viruses of the A(H5) and A(H7) subtypes are of particular importance, as they can evolve into highly pathogenic avian influenza viruses (HPAIVs) in terrestrial poultry, causing severe hemorrhagic disease and outbreaks with high mortality. A(H5) HPAIVs from the A/goose/Guangdong/1/1996 (GsGd) lineage were first detected in Hong-Kong in 1997 (*4, 5*) and have been detected on all continents since (*6–8*). This virus lineage has become enzootic in poultry and wild birds in many countries of the world (*6*). A(H5) GsGd HPAIVs have infected more than 60 mammalian species (*9*), caused mass mortality events in marine mammals (*10, 11*) and widespread outbreaks in dairy cows in the US (*12*), leading to substantial animal health consequences and economic losses. Moreover, the spillover of A(H5) HPAIV to humans has resulted in severe cases of disease, with 520 fatalities out of 993 confirmed human cases (*13–15*), raising global concerns for a new pandemic. Continuous virus circulation in birds has led to the diversification of the HA of GsGd HPAIVs, both at the genetic and antigenic level. Currently, GsGd HPAIVs belonging to genetic clades 2.3.2.1 and 2.3.4.4 are dominating in various parts of the world (*6*). This antigenic diversity has prompted the World Health Organization (WHO) to biannually select multiple candidate vaccine viruses (CVVs) (*16, 17*). Since 2006, 44 candidates have been selected by the WHO, highlighting the challenges posed by the antigenically diverse GsGd HPAIVs to vaccine design and pandemic preparedness. Although this approach is necessary at this stage, the continuous updating of CVVs is a reactive and unsustainable approach to prepare against A(H5) HPAIVs that might emerge and cause outbreaks in humans in the future. At present, manufacturing an inactivated vaccine that antigenically matches the pandemic virus remains the gold standard in influenza pandemic vaccine strategies, which usually takes up to six months, during which the population is vulnerable to infection. Such vaccine formulations against avian influenza viruses generally suffer from low immunogenicity, owing to the need to induce a primary immune response in an immunologically naive population, and to the intrinsic low immunogenicity of avian influenza viruses (*18–20*). While alternative vaccine platforms that may offer higher immunogenicity are under investigation, such as mRNA and vector-based approaches, the choice of vaccine antigen will be a crucial component of vaccine effectiveness.

An integrated analysis of the global antigenic evolution and historical diversification of the A(H5) GsGd HPAIVs is missing, yet will be crucial to design vaccine antigens that induce immunity against antigenically distinct viruses. We set out to characterize the global antigenic evolution of A(H5) HAs by creating a high-resolution antigenic map, using a large collection of historical, recent and current A(H5) influenza viruses. This map was used to design immunogenic and antigenically central vaccine antigens conferring broad reactivity to antigenically distinct viruses, and to visualize antibody-mediated immune responses. Proof-of-concept of high immunogenicity, broad reactivity, and protection is provided using ferrets as a preclinical model.

### High resolution antigenic map of A(H5) avian influenza viruses

We selected representative HA genes of (sub)clades along the phylogenetic tree of A(H5) influenza viruses (Fig. 1A, table S1, Data S1; all supplementary data files are also available at https://epiv-lab.github.io/H5-antigenically-central-vaccine/). HAs from non-GsGd viruses from the American and Eurasian lineages, and HAs of WHO CVVs or closely related viruses were also included in the selection (Fig. 1A, table S2). For viruses that were not already present in-house, synthetic HA genes were ordered and cloned into plasmids to produce recombinant viruses in the genetic background of the attenuated strain A/Puerto Rico/8/1934 (PR/8), using reverse genetics. Ferret sera were generated against a subset of these viruses based on divergent genetic and antigenic properties, the latter assessed in preliminary assays (Fig. 1A, table S3). The resulting dataset comprised 127 antigens and 33 post-infection sera, which were cross-titrated in hemagglutination inhibition (HI) assays (table S4, supplementary text S1). An antigenic map was then constructed to visualize and quantify the antigenic evolution, and diversification of A(H5) influenza A viruses from 1959 to 2022 (Fig. 1B, Data S2), using multidimensional scaling algorithms described previously (*21*), available via the Racmacs R package (*22*). Distances in antigenic maps are inversely correlated to HI titers, i.e., antigens are close in space with sera against which they react with a high HI titer. Antigenic maps also allow the visualization of antigenic relatedness between antigens, which is not directly measured in the HI assay.

**Fig. 1.**
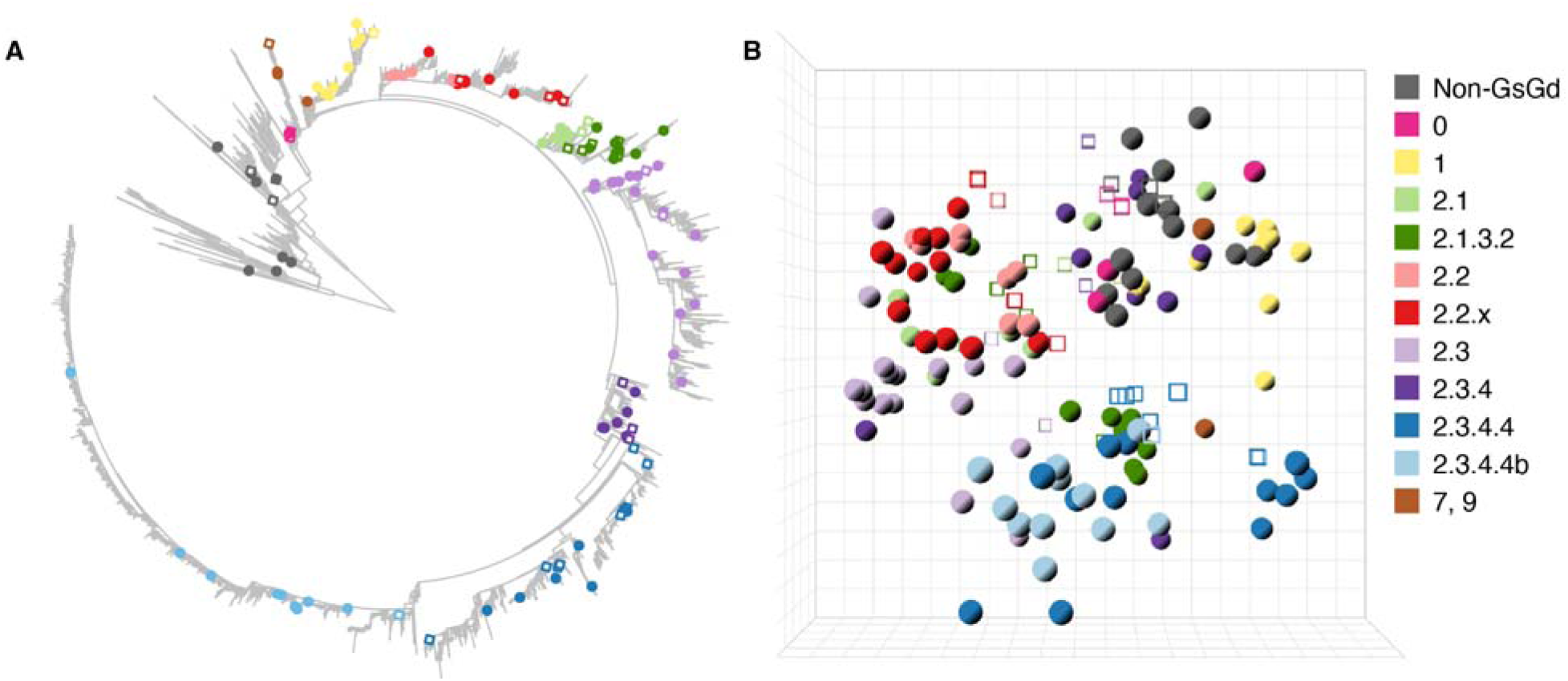
High resolution three-dimensional A(H5) antigenic map comprised of genetically diverse HAs. (**A**) Maximum likelihood phylogenetic tree based on 14896 A(H5) nucleotide sequences, rooted on the midpoint, corresponding to the divergence of the Eurasian and American non-GsGd lineages. The HAs selected for antigenic characterization are highlighted with closed circles or open squares, color-coded based on their respective genetic lineage, as indicated in (B). Open squares represent HAs of viruses against which homologous ferret sera were raised. Zoomable pdf file with isolate names displayed is available in Data S1 and an interactive version of the tree is available at https://itol.embl.de/tree/156831160222541718279907. **(B)** The three-dimensional antigenic map constructed from HI data of 117 antigens titrated against 29 post-infection sera. Antigens are represented as closed spheres and sera as open cubes. Antigens and sera are color-coded based on their respective genetic lineage as shown on the right-hand side of the figure. Each direction (x, y, z) represents antigenic distance, and one square of the grid corresponds to one antigenic unit, defined as a two-fold change in HI titer. The corresponding interactive display (Data S2) provides additional information, e.g., visualization of the map from different angles, and display of antigen and serum names. GsGd: A/goose/Guangdong/1/1996.

The generation and validation of the antigenic map are described in supplementary texts S2 and S3 (see also Data S3, S4, fig. S1-S4). Minimally three dimensions were required to represent the A(H5) HI titer data in an antigenic map. Ten antigens, reacting with fewer than four detectable titers to ferret sera, were removed from the dataset along with four corresponding homologous sera (Table S2), as less than four detectable titers are insufficient to confidently place points in a map of three or more dimensions. Additional information about these antigenic outliers is available in the supplementary text S3. The resulting antigenic map contained 117 antigens and 29 post-infection sera. The three-dimensional antigenic map represented the underlying HI data well, as shown by the correlation between the antigen/serum pairwise distances dictated by the HI titer table and Euclidean distances in the map (R^2^ = 0.64, fig. S3A). Additional analyses were performed to evaluate the accuracy and stability of the antigenic map (supplementary text S3, Data S3, S4, fig. S3, S4). Moreover, a genetically and antigenically diverse representative subset of the antigenic map dataset was used to perform virus neutralization (VN) assays, and VN titers were shown to correlate with the HI titers (R^2^= 0.61, fig. S3D).

The antigenic evolution of A(H5) influenza viruses showed a non-directional pattern over time (Movie 1). Analyses of the genetic and antigenic distances revealed a relative discordance between genotype and antigenic phenotype (fig. S5A). This was exemplified by the fact that antigens from different genetic (sub)clades were in some occasions located in close proximity to one another in the antigenic map (Fig. 1B, Data S2), suggesting that they shared similar antigenic properties. To assess whether this phenomenon might have partially resulted from a distortion of distances in the map due to dimension constraints, we compared raw HI titers of selected antigen pairs with a relatively large pairwise genetic distance (more than 25 amino acid differences in HA1), and a low pairwise antigenic distance (below 1.5 antigenic units (AU), where one antigenic unit corresponds to a two-fold change in HI titer). For 19 out of 24 pairs, similarity in HI reactivity was indeed observed (fig. S6, R^2^>0.5), providing evidence that this observation was not due to dimension constraints, and further supporting the discordance between genetic and antigenic properties of certain A(H5) antigens.

The maximum pairwise distances between the non-GsGd antigens, covering a span of 43 years, was only 5.58 AU. In contrast, the GsGd antigens, spanning 25 years, were positioned at a maximum pairwise distance of 14.71 AU. This highlights the increased level of antigenic evolution relative to the timespan in the GsGd lineage as compared to the non-GsGd antigens (fig. S5B-D, Movie 1). Within the GsGd lineage, relatively little antigenic evolution was observed for clade 1 and 2.2 (table S5). In contrast, antigens from clade 2.1 and 2.3 exhibited greater antigenic evolution with maximum distances of 10.21 and 14.71 AU between antigens, respectively. Antigenic diversity was also observed within the currently dominating 2.3.2.1 and 2.3.4.4 clades, with maximum pairwise distance of 7.37, 8.80 and 5.72 AU in clade 2.3.2.1, 2.3.4.4 and 2.3.4.4b, respectively. In addition, antigenic differences were observed between antigens from clade 2.3.2.1 and 2.3.4.4 (table S5), which should be taken into account for (pre)pandemic vaccine design. The maximum pairwise distance between antigens from these clades was 11.87 AU, and the antigenic distance between the most recent clade 2.3.2.1 and 2.3.4.4 antigen was 6.84 AU. These observations underscore the challenge posed by the antigenic diversity of GsGd A(H5) viruses for pandemic preparedness, including vaccine design. To address challenges posed by the antigenic diversification of A(H5) avian influenza viruses, the WHO has selected over the years 44 CVVs (*16, 17*). Highlighting the A(H5) WHO CVV(- like) antigens (table S2) present in our dataset in the antigenic map revealed a good coverage of the antigenic space (Data S5A), but frequent updates are required to cope with antigenic evolution.

### Design of a central vaccine antigen

An alternative approach to the continuous production of CVVs would be the selection or design of antigen(s) inducing highly cross-reactive immune responses. Theoretically, high cross-reactivity could be achieved in a non-directional space just with one antigen provided that (i) it elicits an immune response centered in antigenic space and (ii) it is highly immunogenic, i.e. induces a high homologous HI titer. Interestingly, the maximum distance of antigens from the center of the map has not increased significantly since 2010, despite ongoing antigenic evolution (fig. S7A), suggesting that the A(H5) antigenic space might be constrained. This observation underscores the feasibility and potential sustainability of approaches based on the design of immunogenic and antigenically central vaccine antigens.

To design an immunogenic and antigenically central antigen, our initial exploration focused on wild-type antigens close (within a 2.5 AU radius) to the center of the antigenic map, defined as the arithmetic mean position of all the antigens in the antigenic map. Twelve antigens qualified (table S2, “Notes” column), which reacted to a higher number of sera from the antigenic map with a higher geometric mean titer (GMT) as compared to antigens located further away from the map center (fig. S7B, C). Eleven of the twelve centrally located antigens lacked a putative glycosylation site at position 154 (A(H5) numbering throughout the manuscript (*23*)), a feature present in 37% of the GsGd antigens of the antigenic map. The removal of putative HA glycosylation sites has previously been documented to enhance vaccine immunogenicity (*24, 25*). In addition, several of these centrally located antigens harbored amino acids such as 94N, 156A, 189R, 210I and 223N, previously associated with alterations in the HA receptor binding profile (*26, 27*). These observations, coupled with evidence from the literature highlighting the impact of receptor binding amino acid changes on increased HA cross-reactivity and immunogenicity (*24, 28*), prompted us to investigate the receptor binding profile of these twelve centrally located antigens. Notably, eight out of the twelve centrally located antigens exhibited a certain level of binding to α2,6 sialic acids (table S6).

Building upon these insights and upon the expectation that the next pandemic influenza virus will possess an α2,6 sialic acid binding specificity, we rationally engineered candidates vaccine antigens (CVAs) based on HAs of viruses from distinct genetic clades. Substitutions were introduced at positions altering the receptor binding profile (222QL and 224GS (*29*)) and removing a putative glycosylation site at position 154 (156TA), resulting in CVA-Vietnam, based on A/Vietnam/1194/2004 (clade 1), CVA-Indonesia, based on A/Indonesia/5/2005 (clade 2.1.3.2), and CVA-Anhui, based on A/Anhui/1/2005 (clade 2.3.4). These antigens exhibited a dual receptor binding profile to both α2,3 and α2,6 sialic acids, whereas their parent HAs exclusively bound to α2,3 sialic acids (table S6).

The rationally designed antigens were tested in HI assays against all sera from the dataset (table S7), and subsequently individually placed in the antigenic map. CVAs exhibited increased HI reactivity and, on average, were situated at a distance of 1.32 AU from the map center. Notably, their positions were on average 4.17 AU closer to the map center than their respective wild-type counterparts. To assess the immunogenicity of the CVAs, two ferrets were vaccinated and boosted with Addavax-adjuvanted whole-inactivated vaccines, 28 days apart. As a comparator, one of the natural central antigens mentioned earlier, A/Iraq/755/2006, was employed (Iraq).

Antibody profiles were generated for serum samples obtained four weeks after the boost vaccination (Iraq_VACC_, CVA-Vietnam_VACC,_ CVA -Indo_VACC_, CVA -Anhui_VACC_) by HI titration against 113 antigens of the antigenic map (Fig. 2, Data S6; Data S7 and table S8 for data from individual animals). With the exception of one Iraq_VACC_ serum, all sera had detectable titers against over half of the tested antigens. Although the position of the CVA-Anhui_VACC_ was slightly less central than CVA-Indonesia_VACC_ (1.66 versus 0.80 AU from the center, respectively), CVA-Anhui induced the broadest HI response, characterized by the highest GMT (43) against all A(H5) antigens, as well as by the highest GMT against viruses from currently circulating genetic clades 2.3.2.1 and 2.3.4.4 (29 versus 16 for CVA-Indonesia). CVA-Anhui was then selected for further investigation, due to its distinct characteristics.

**Fig. 2.**
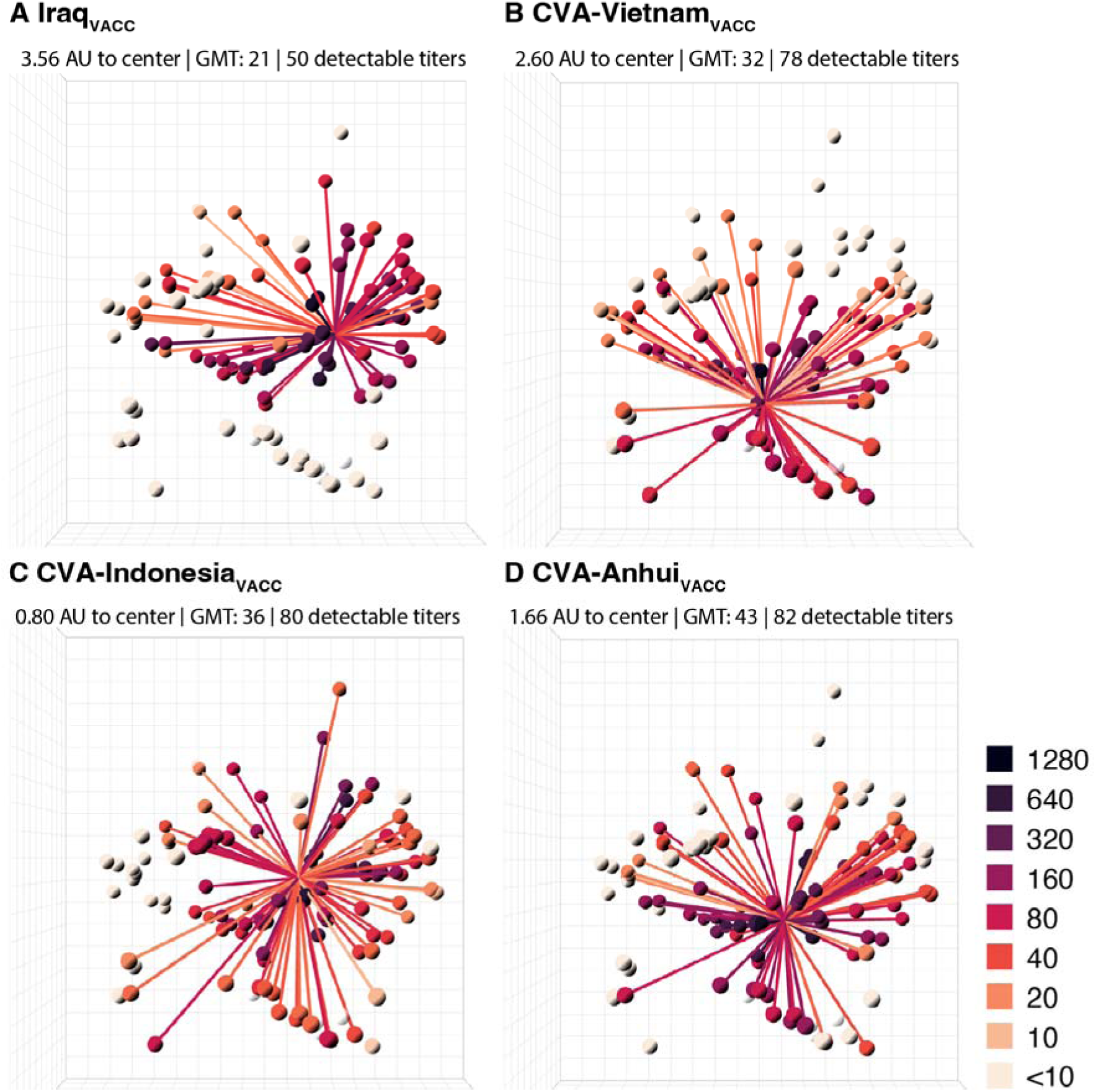
Antibody profiles illustrate the broad cross-reactivity in A(H5) antigenic space of ferret sera raised by vaccination with whole-inactivated vaccines containing engineered HAs. For each vaccine, (**A**) Iraq_VACC_, (**B**) CVA-Vietnam_VACC_, (**C**) CVA-Indonesia_VACC_ and (**D**) CVA-Anhui_VACC_, antibody profiles are displayed in the antigenic map from Fig. 1B. Each plot displays the mean titers for two vaccinated ferrets. Lines connect the position of the serum with the antigens against which a titer above the assay’s detection limit was observed (i.e. ≥ 10). Antigens and connecting lines are colored by the HI titer as indicated in the right-hand legend. Numbers on top of each panel indicate the distance of the mean serum position to the center of the antigenic map, the overall GMT of the mean serum, and the mean number of titers above the detection limit per serum. The four antigens that were not titrated against these sera are shown in a transparent white color. One CVA-Anhui_VACC_ animal reached humane endpoints during the study because of a malignant lymphoma, which was unrelated to the experimental procedures. Due to the premature sacrifice of this ferret, data of a single serum were used for analysis and visualization in (D). GMT: geometric mean titer.

Assessment of the genetic stability of the CVA-Anhui virus through 5 to 10 serial passages in embryonated chicken eggs, the substrate most frequently used to produce influenza virus vaccines, revealed the selection of amino acid substitution 134TA in HA in four out of five independent experiments. When this substitution was introduced in CVA-Anhui, it stabilized the virus across two independent experiments of five serial passages. Given that 134A is consistently found in all antigens in the antigenic map and is also present in two database entries of the A/Anhui1/2005 HA (table S2), it was incorporated in the CVA-Anhui antigen (A/Anhui/1/2005_134TA, 156TA, 222QL, 224GS_). HI assays in which the resulting CVA-Anhui antigen was titrated against all sera in the antigenic map, revealed its closer proximity to the antigenic map center compared to A/Anhui/1/2005_156TA, 222QL, 224GS_ (1.28 AU versus 2.09 AU). This antigenically central (AC) antigen, from here on termed AC-Anhui, underwent subsequent evaluation in preclinical studies in ferrets.

### AC-Anhui_VACC_ provides non-inferior protection against antigenically divergent A(H5) HPAIVs in ferrets compared to homologous vaccines

Preclinical studies in ferrets were performed to assess (i) the protective capacity of AC-Anhui_VACC_ as compared to the standard of care, i.e., vaccines homologous to challenge viruses, and (ii) the impact of the engineered substitutions on the height and breadth of the neutralizing antibody response, and protection against challenge.

The selected challenge viruses were A(H5N1) clade 2.2.1.2 A/duck/Giza/15292S/2015 (Giza) and A(H5N6) clade 2.3.4.4a A/Sichuan/26221/2014 (Sichuan). These viruses were chosen because their respective HAs are genetically and antigenically distinct from one another (9.20 AU apart), and from the A/Anhui/1/2005 virus (7.40 and 10.59 AU, respectively), which was the wild-type counterpart of AC-Anhui (Data S5B, table S5). Groups of six ferrets were vaccinated twice, 28 days apart, with Addavax-adjuvanted split-inactivated vaccines containing either the antigenically central HA (AC-Anhui_VACC_), the wild-type counterpart HA (Anhui_VACC_), the challenge viruses’ HAs (Giza_VACC_ and Sichuan_VACC_), or phosphate-buffered saline (PBS) (Mock_VACC_) (fig. S8). We chose to perform pre-clinical work using split-inactivated vaccines, given that the only licensed A(H5) pre-pandemic vaccine for human use is of this type. To isolate the effects of varying the HA antigen, the neuraminidase present in the vaccine was mismatched with that of the challenge virus (fig. S8).

Upon boosting, an increase in HI titer against the vaccine antigens of the respective study was observed for virtually all vaccinated animals (table S9). The post-boost sera were titrated against 113 antigens of the antigenic map, and this HI data was utilized to generate mean antibody profiles for each experimental group (Fig. 3, Data S8; Data S9-10 and table S8 for data from individual animals). The mean AC-Anhui_VACC_ post-boost serum was positioned in closer proximity to the center of the antigenic map than that raised upon vaccination with wild-type antigens (Fig. 3, Data S8). In addition, the GMT and mean number of antigens with a detectable HI titer were higher for the mean AC-Anhui_VACC_ post-boost serum (Fig. 3B, E) than for the mean Anhui_VACC_ post-boost serum (Fig. 3A, D). These observations indicated that the engineered substitutions led to a more central and enhanced immune response, both in terms of height and breadth. It is noteworthy that the height and breath of immune responses in both the Anhui_VACC_ and AC-Anhui_VACC_ groups from the Giza study exceeded those from the Sichuan study, potentially attributable to the use of a different batch of ferrets and vaccines. Anhui_VACC_ failed to induce detectable HI titers against the Giza and Sichuan challenge viruses in any of the animals (table S8, S9). In contrast, AC-Anhui_VACC_ induced detectable HI titers against the challenge viruses in all ferrets of the Giza challenge and four out of six ferrets of the Sichuan challenge, showing a comparable response to that observed for the homologous vaccines (table S8, S9).

**Fig. 3.**
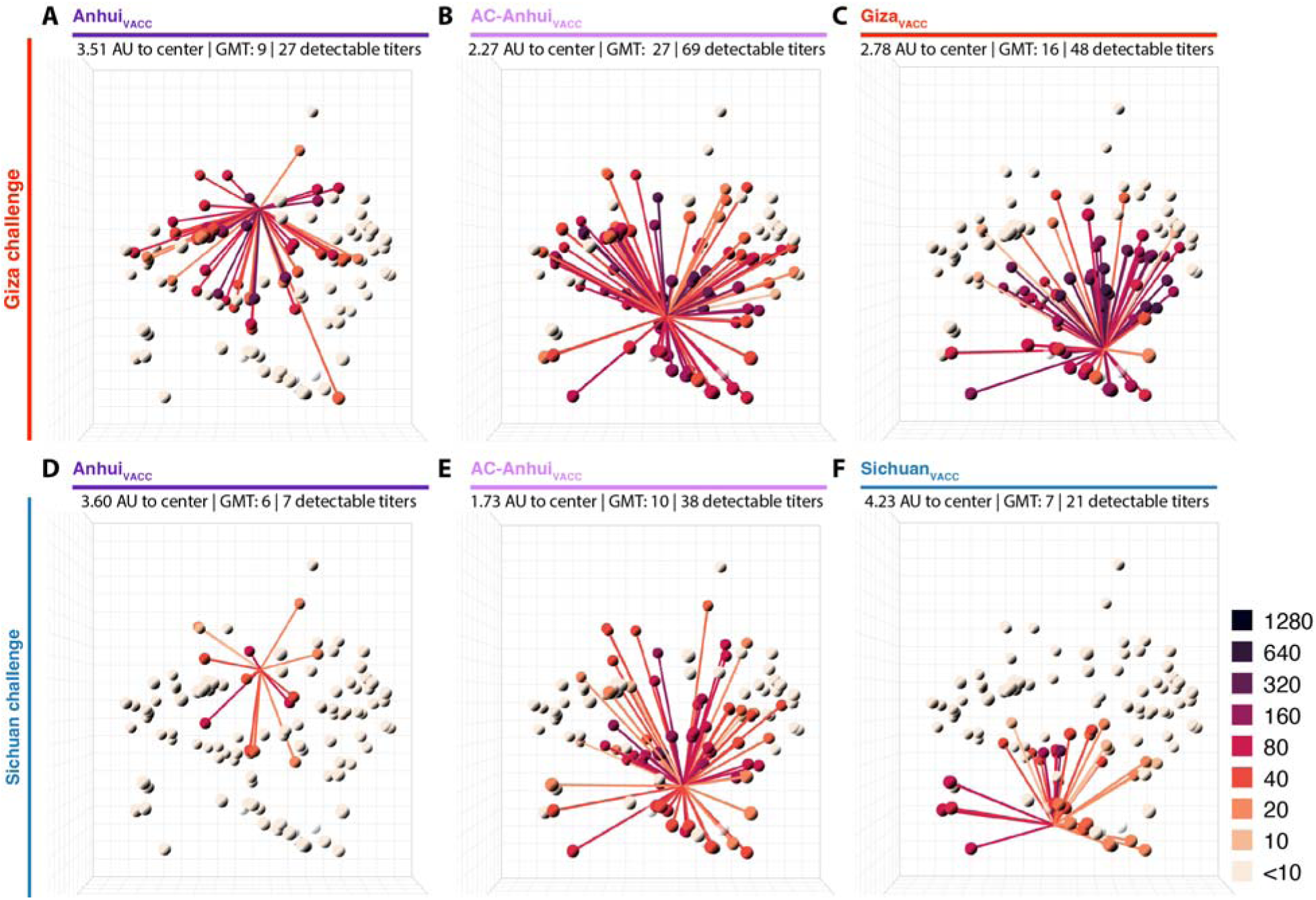
Broader, higher, and more central antibody responses in ferrets upon vaccination with split-inactivated vaccines containing an engineered HA as compared to its wild-type counterpart. For each vaccine group, antibody profiles representing the mean HI titers per group (n=6) are displayed in the antigenic map from Fig. 1B. Antibody profiles of ferrets from the vaccination-challenge with the Giza (A-C) and the Sichuan (D-F) virus, vaccinated with vaccines containing the following HA: (**A**, **D**) Anhui_VACC_, (**B**, **E**) AC-Anhui_VACC_, (**C**) Giza_VACC_ or (**F**) Sichuan_VACC_. Same representation as in Fig. 2. For an interactive version of this figure, see Data S8. AU: antigenic unit; GMT: geometric mean titer.

Four weeks after the administration of the second vaccine dose, ferrets were inoculated intranasally and intratracheally with the Giza virus or the Sichuan virus, at doses determined in preliminary studies to induce a reproducible and consistent upper and lower respiratory tract infection. Nose and throat swabs were collected daily, alongside monitoring of body weight, body temperature, clinical signs and activity (fig. S8). Four days post-inoculation (dpi), ferrets were euthanized and relevant tissues, selected based on virus detection in preliminary studies, were collected for virological and histopathological analyses (fig. S8).

Upon Giza virus inoculation, Mock_VACC_ ferrets showed a maximum mean body weight reduction of 11.4% (Fig. 4A), and reduced activity from three dpi onwards (Fig. S9A), accompanied by an increased breathing frequency at four dpi. In contrast, ferrets from the three vaccinated groups showed no significant alteration in body weight, activity nor clinical status (Fig. 4A, Fig. S9A). In the Mock_VACC_ group, the mean increase in body temperature between one and four dpi (2 °C) was significantly higher compared to the Anhui_VACC_ (1.3 °C), AC-Anhui_VACC_ (1.1 °C) and Giza_VACC_ (0.9 °C) groups (Fig. 4C, D, fig. S10). Unsurprisingly, infectious virus was recovered from nose and throat swabs of ferrets from all groups, corroborating that intramuscular immunization does not prevent upper respiratory tract infection (fig. S9C, E). While infectious virus was isolated from respiratory tissues of all Mock_VACC_ animals at four dpi (Fig. 5A), significantly lower virus levels were observed in nasal turbinates, tracheas, bronchi, and lungs from vaccinated ferrets (Fig. 5A). In fact, many vaccinated animals showed no infectious virus in trachea, bronchus, and lung samples (Fig. 5A), indicating robust protection of vaccination against lower respiratory tract infection with the Giza virus, in spite of undetectable HI titers in some animals. The mean relative lung weight, reflective of pulmonary inflammatory infiltrate and edema, was significantly higher in the Mock_VACC_ group compared to the vaccinated groups (fig. S9G). Histopathological lesions and lymphocyte presence in respiratory tissues were consistent with viral replication and correlated with antigen expression detected by immunohistochemistry (IHC), as detailed in supplementary text S4, table S10, and fig. S11. Overall, vaccination strongly reduced disease severity and viral replication in the lower respiratory tract upon Giza virus inoculation, irrespective of the used vaccine. However, AC-Anhui_VACC_ and Giza_VACC_ animals showed significantly reduced mean virus titers in the cerebrum compared to Mock_VACC_ group, while Anhui_VACC_ animals did not. These results showcased the non-inferiority of the antigenically central antigen to the homologous Giza vaccine and demonstrated its ability to prevent extra-respiratory spread of the Giza virus to the brain.

**Fig. 4.**
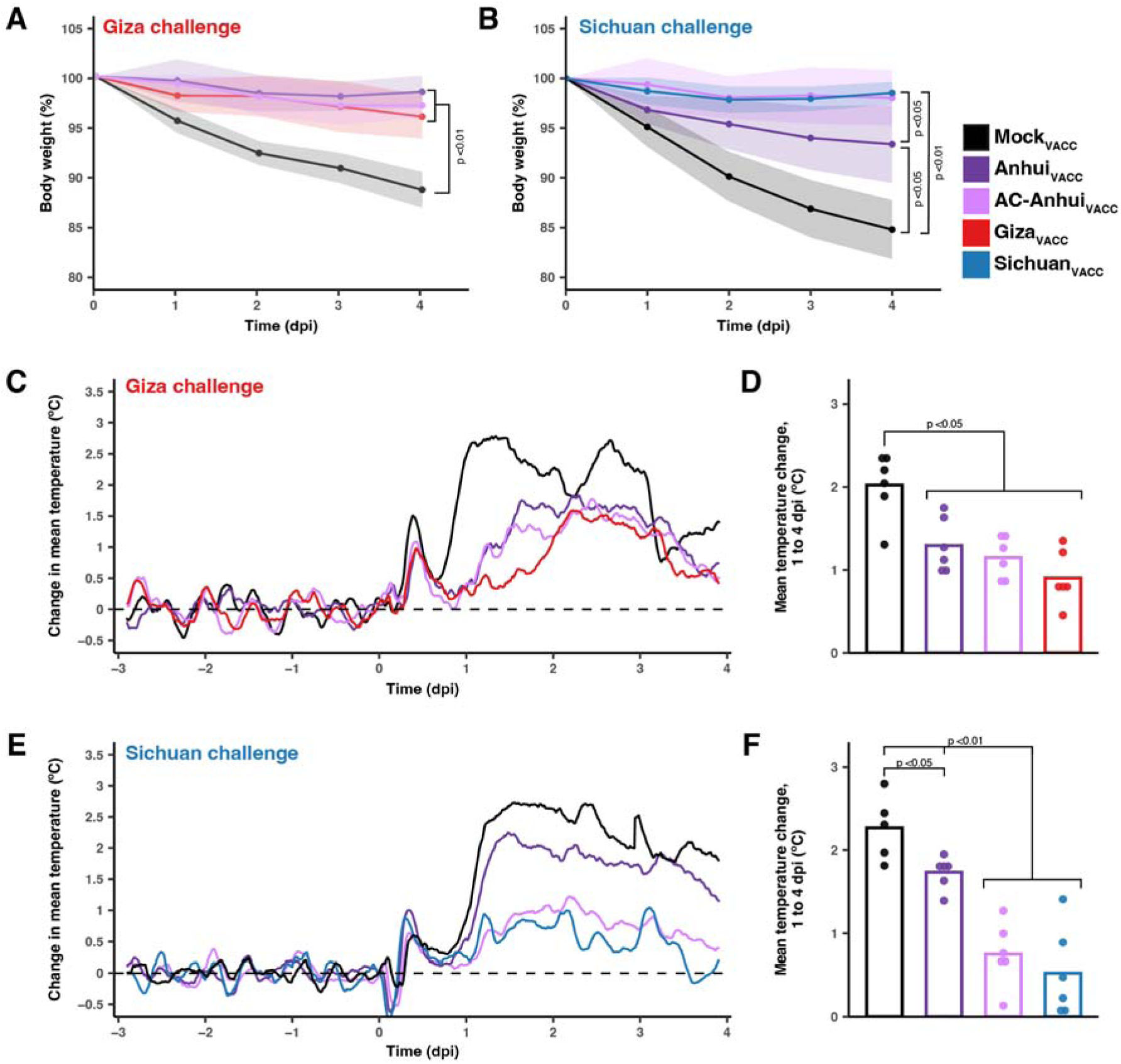
Ferret body weight and temperature changes reveal non-inferior protection of AC- Anhui_VACC_ vaccination compared to homologous vaccination. (**A**, **B**) Body weight expressed as a percentage of starting body weight of ferrets (n=6 per group) challenged with the Giza (A) or Sichuan (B) virus. Lines connect the daily arithmetic means. Shaded areas indicate the standard deviation of the mean per group. (**C, E**) Temperature change from baseline (mean temperature recorded during the three days prior inoculation, indicated as dashed line) of ferrets challenged with the Giza (C) or Sichuan (E) virus. Per group (n=6, n=5 for Mock_VACC_ in Sichuan challenge due to temperature probe malfunctioning), the mean of individual four-hour sliding means is displayed. (**D**, **F**) Mean temperature change from baseline between 1 and 4 dpi of ferrets challenged with the Giza (D) or Sichuan (F) virus. Dots show data of individual animals. In (E, F), data of the two deceased animals in the Mock_VACC_ group are included in visualization and analysis up until three dpi. Statistically significant differences, as determined with pairwise Mann-Whitney tests, are indicated with the corresponding p-value in (A, B, D, F). In (A, B), areas under the curve were used for statistics. Dpi: days post-inoculation.

**Fig. 5.**
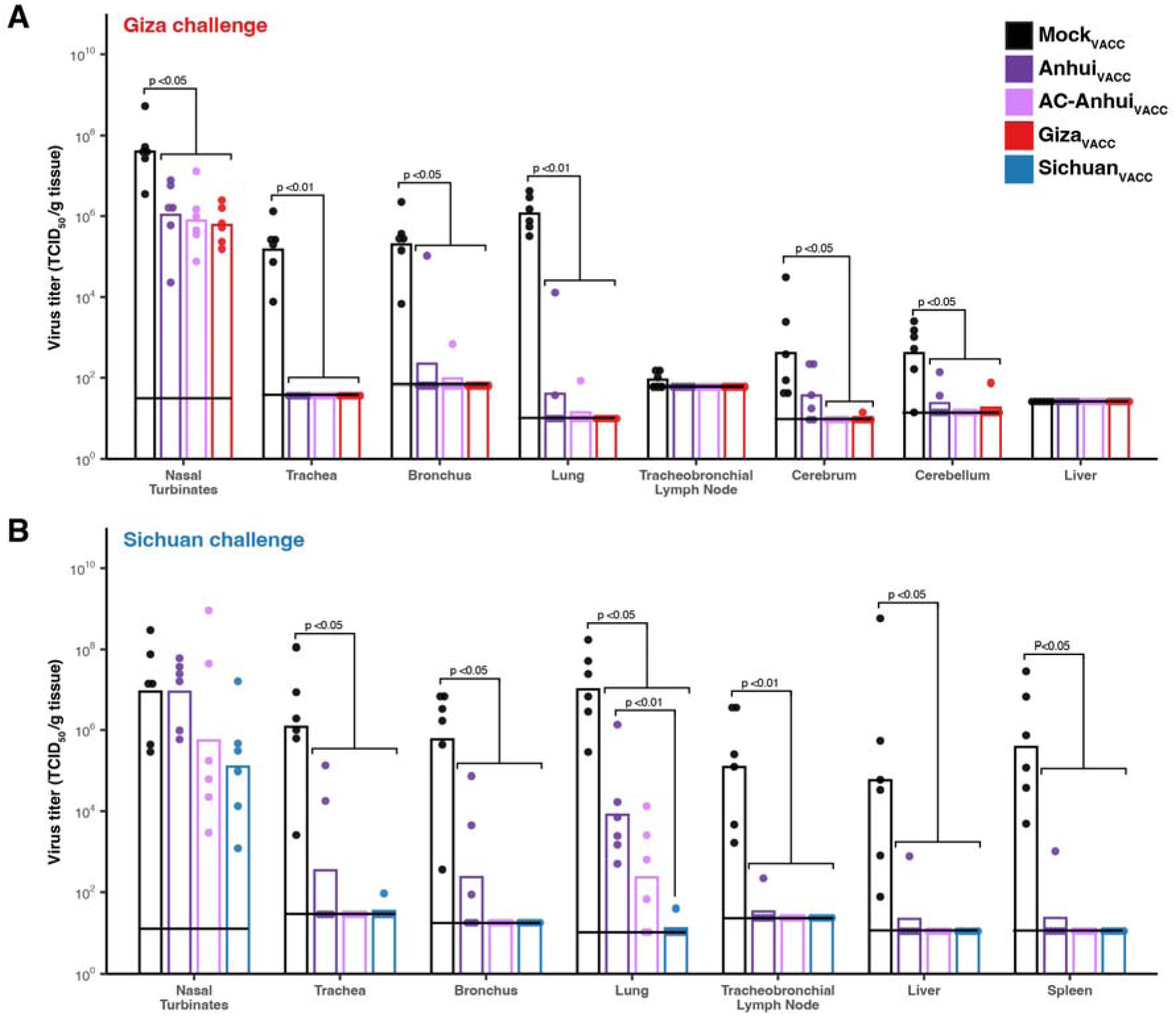
Vaccination with AC-Anhui reduced infectious virus titers in ferret tissues equally well as homologous vaccination. (A) Vaccination-challenge with the Giza virus. (B) Vaccination-challenge with the Sichuan virus. Data are color-coded based on vaccine group as indicated in the legend. Bars represent the geometric mean titer (TCID_50_/g tissue) per group (n=6). Dots represent titers in tissues of individual animals. Horizontal black lines indicate the detection limit for each tissue. Statistically significant differences, as determined with pairwise Mann-Whitney tests, are indicated with the corresponding p-value. TCID_50_: 50% tissue culture infectious dose.

The Sichuan virus challenge resulted in more severe disease compared to the Giza virus. Mock_VACC_ animals displayed reduced activity from three dpi onwards (fig. S9B). At four dpi, two animals were found dead, and the remaining four exhibited difficulty walking and a large reduction in activity (fig. S9B). In the vaccinated groups, no significant changes in activity score (fig. S9B) or clinical signs were observed upon Sichuan virus inoculation. However, Anhui_VACC_ and Mock_VACC_ ferrets experienced a mean body weight reduction of 6.6% and 15.2% respectively, significantly different from AC-Anhui_VACC_ and Sichuan_VACC_ ferrets (Fig. 4B). Moreover, Anhui_VACC_ ferrets exhibited a significantly higher temperature increase (1.7 °C) compared to AC-Anhui_VACC_ (0.7 °C) and Sichuan_VACC_ ferrets (0.5 °C) groups (Fig. 4E, F, fig. S10). As expected, infectious virus was isolated from nose and throat swabs from two dpi onwards in the majority of animals across all groups, as observed for the Giza challenge (fig. S9D, F). In the Mock_VACC_ group, infectious virus was isolated from respiratory tissues, liver, and spleen of all animals, demonstrating extra-respiratory spread of the virus, confirmed by histopathological and IHC analysis (Fig. 5B, fig. S11, supplementary text S4, table S10). Respiratory tract virus titers were generally lower in AC-Anhui_VACC_ and Sichuan_VACC_ ferrets as compared to Anhui_VACC_ animals, with significant differences observed in lung titers between Sichuan_VACC_ and Anhui_VACC_ animals, but not between Sichuan_VACC_ and AC-Anhui_VACC_ animals (Fig. 5B). Mean relative lung weight (fig. S9H), histopathological, and IHC analysis of lung tissue samples corroborated these results (supplementary text S4, table S10 and fig. S11). AC-Anhui_VACC_ and Sichuan_VACC_ also protected all animals against extra-respiratory virus spread, while infectious virus was isolated from the liver and spleen of one Anhui_VACC_ animal (Fig. 5B). Analogous to the Giza virus challenge, AC-Anhui_VACC_ conferred non-inferior protection compared to Sichuan_VACC_, despite more severe disease. In addition, AC-Anhui_VACC_ outperformed Anhui_VACC_ in reducing infection severity, evidenced by reduced body weight loss, body temperature increase, virus titers, histopathological changes, antigen expression in the lungs, and extra-respiratory virus spread.

## Discussion

Here, we employed antigenic cartography to visualize and quantify the antigenic evolution of A(H5) avian influenza viruses and design antigenically central vaccine antigens. Unlike the directional evolution observed for human seasonal A(H3) influenza viruses (*21, 30, 31*), the A(H5) antigenic evolution exhibited non-directionality, and more than two dimensions were required to retrace complex evolution patterns, as seen for other influenza viruses circulating in animal populations (*32–37*). Discordance between antigenic and genetic evolution was observed, reminiscent of findings on avian A(H7) influenza viruses (*32*) and other animal influenza viruses (*36*). This suggests that A(H5) HA global antigenic evolution might be driven by only a few amino acid changes, as already proposed (*33, 38, 39*), and similar to what has been observed for human seasonal influenza viruses (*30, 31, 40*) and other animal influenza A viruses (*32, 35*).

The evolutionary patterns of animal influenza A viruses are difficult to understand and predict because underlying drivers remain unclear. Unlike in humans, in which population immunity buildup leads to the selection of drifted influenza viruses evolving away from all previously circulating variants, herd immunity is not expected to accumulate as such in poultry, allowing the co-circulation of antigenically diverse viruses. This is probably primarily attributable to the low reinfection likelihood given segregation of animal populations in time and location, shorter life spans, virus lethality, and rapid renewal of susceptible populations. These aspects are particularly relevant for terrestrial poultry, the primary hosts of A(H5) GsGd viruses. Recent enzootic establishment of A(H5) GsGd in various wild bird species (*41*) may result in different patterns of antigenic evolution. On the other hand, vaccination of domestic poultry, carried out in certain countries to mitigate avian influenza virus outbreaks, has been proposed as potential cause of antigenic evolution of avian influenza viruses (*42, 43*). However, it is improbable that vaccination alone can fully explain the observed antigenic evolution of A(H5) GsGd viruses, especially during the early diversification period around 2004-2005, when vaccination campaigns were in early implementation stages. Alternatively, antigenic evolution of the HA of A(H5) GsGd viruses could be in part a bystander effect of the accumulation of substitutions in the HA resulting from other adaptive processes, such as adaptation of the virus to replicate systemically in terrestrial birds or to switch between multiple host species. Understanding whether the other main antigen of influenza viruses, NA, undergoes antigenic evolution in avian hosts would be valuable to further identify overall underlying drivers of antigenic evolution of avian influenza viruses.

The diverse antigenic landscape of A(H5) influenza viruses poses significant challenges to pandemic preparedness and the design of effective vaccines against antigenically diverse viruses. In the pursuit of universal influenza virus vaccines, subtype-wide vaccines offering protection against drifted variants have been identified as crucial initial steps towards the development of truly universal vaccines (*44*). Previous subtype-wide approaches involved the design of reconstructed ancestral HAs (*45*), genetic consensus HA antigens (*46–52*), mosaic HAs with conserved T and B cell epitopes (*53*) or phylogenetically central antigens (*54*), all with the aim to recapitulate the genetic diversity of A(H5) viruses. While these synthetic antigens occasionally induced broader antibody responses than wild-type comparators, they are representative of a population of genetic sequences rather than of the antigenic properties of a population of viruses. Given that antigenic phenotype might be governed by only a few amino acid changes, genetic information at the level of the full HA gene may not be predictive of antigenic phenotype. In our approach, we focused on designing antigens based on antigenic phenotype rather than genetic information. Our observations indicated a relatively stable A(H5) antigenic space, which, along with a non-directional pattern of antigenic evolution, supported the design of antigenically central HA antigens. Toward this goal, we simultaneously altered the receptor binding specificity and glycosylation status of A(H5) HAs, inspired by natural viruses located centrally in antigenic space. Anticipating the emergence of an influenza pandemic virus with an α2,6 sialic acid binding specificity, it is rational to adjust the receptor binding specificity of avian influenza vaccine candidates to mirror that of human influenza viruses. The breadth and height of polyclonal antibody responses upon vaccination was assessed against over a hundred different A(H5) antigens, leveraging the antigenic map as unique tool to visualize antibody responses. The CVAs demonstrated increased reactivity in HI assays and induced robust and central antibody responses within the antigenic space. Vaccination with AC-Anhui_VACC_ conferred non-inferior protection against two genetically and antigenically distinct challenge viruses as compared to homologous antigens, representing the standard of care. Particularly in the Sichuan virus challenge, the antigenically central antigen outperformed its wild-type counterpart in reducing disease severity.

There are avenues to further improve the immunogenicity and breadth of A(H5) antigenically central influenza virus vaccines. While vaccination with AC-Anhui_VACC_ provided a good coverage within clade 2.3.4.4 antigenic space, coverage in the 2.3.2.1 space could be enhanced. Expanding antigenic space coverage could be achieved through heterologous prime-boost strategies utilizing vaccine antigens situated in different regions of the map (*20*), and/or by enhancing the immunogenicity and breadth of emerging variants using the same design as described here. Increased knowledge on the molecular determinants of antigenic change of A(H5) HAs would also offer opportunities to further tune the positioning of vaccine antigens in space. Immunogenicity and breadth enhancement could be achieved by employing different vaccine platforms. Here, we chose to use inactivated vaccines, given that the only currently licensed A(H5) pre-pandemic vaccine is of this type. We observed that whole-inactivated vaccines were more immunogenic than split-inactivated vaccines, as reported previously in unprimed individuals (*55*). However, antigenically central vaccine antigens could be utilized in conjunction with other vaccine platforms that may induce higher B and T cell immune responses, such as mRNA vaccines (*56*) or vector-based vaccines. Alternative routes of administrations that may elicit mucosal immunity in addition to systemic immunity will also be important in designing vaccines reducing not only severe disease but also transmission. Lastly, our study has been focused on the design of immunogenic and broad HAs. However, there is a growing body of evidence supporting the role of immunity against NA in conferring protection against influenza viruses (*57–60*). Here, we deliberately mismatched the NAs in antigenically central vaccines with that of challenge viruses to specifically study the impact of our antigen design on HA on vaccine immunogenicity, cross-reactivity, and protective capacity. However, ensuring the matching of the NA subtype in (pre-)pandemic vaccines with that of emerging A(H5) viruses will be of the utmost importance. Recent human infections caused by GsGd viruses carried the N1 or the N6 NA subtype. While cross-reactive NA antibodies to avian N1 NAs have been detected in humans due to past infections and/or vaccinations with seasonal influenza H1N1 viruses (*61–63*), humans are most likely immunologically naïve to N6 NAs. Understanding the immunogenicity and potential antigenic evolution of avian NAs could lead to the design of antigenically central neuraminidases, which might be instrumental for developing more effective (pre-)pandemic vaccines against A(H5) GsGd viruses.

We acknowledge two limitations within our current study. Firstly, immunogenicity and cross-reactivity was solely evaluated based on height and breadth of the polyclonal antibody response as measured in HI assays. Investigating non-neutralizing antibody responses and T-cell responses would offer valuable complementary data. In addition, understanding monoclonal antibody reactivity profiles would provide deeper insights into mechanisms underlying the observed increased immunogenicity and breadth. Secondly, further investigations are warranted to elucidate the relative contributions of receptor binding properties and glycosylation status to the increased immunogenicity and breadth of AC-Anhui_VACC_. Previous studies have reported increased immunogenicity attributed to α2,6 sialic acid binding specificity, sometimes in synergy with removal of glycosylation sites (*24*, *64–67*). However, these observations were made primarily in the context of live attenuated vaccines, and the mechanisms driving enhanced immunogenicity of α2,6 sialic acid binding vaccine viruses within inactivated formulations are likely different. These mechanisms may include improved antigen presentation resulting from enhanced specific binding to antigen-presenting cells, rather than solely relying on replication efficiency, as seen in live attenuated vaccines.

The antigenic map presented here will offer possibilities to further monitor the emergence of new A(H5) antigenic variants in the global context of the historical diversification. Assessing breadth of pre-pandemic vaccines is crucial given the diverse antigenic landscape of A(H5) viruses. Data from human clinical studies in which A(H5) pre-pandemic vaccines were assessed have often been limited to reactivity against a handful of heterologous viruses (*20*), old antigenic variants in most cases. Selection of a panel of antigens recapitulating the A(H5) antigenic space and current diversity for assessment of human immune responses could be further informed using the present antigenic map. Given the intrinsic low immunogenicity of natural avian influenza viruses, it is anticipated that a rationally designed immunogenic and broad A(H5) vaccine would offer better antibody responses than any vaccine based on a wild-type antigen matched to a potential pandemic strain. Antigenically central vaccines could also be used as prime in pre-pandemic time or at the beginning of a pandemic until a matched vaccine is available. The present work thus provides a proof-of-concept for utilizing antigenic cartography to design and evaluate antigenically central A(H5) vaccine antigens, a strategy which could be expanded to other animal influenza A viruses with zoonotic or pandemic potential. The proof-of-concept presented here warrants follow-up with immunogenicity and breadth studies in humans to improve pandemic preparedness.

## Acknowledgments

We gratefully acknowledge all data contributors, i.e., the Authors and their Originating laboratories responsible for obtaining the specimens, and their Submitting laboratories for generating the genetic sequence and metadata and sharing via the GISAID Initiative, on which this research is based. Peter Bogner of the GISAID initiative is acknowledged in particular, for helpful discussions and advice on appropriate benefit sharing. We thank the following institutes for providing viruses and/or sequences for phenotypic characterization: St Jude Children’s Hospital (United States), the Southeast Poultry Research Laboratory (United States), Center for Disease Control and Prevention (United States), University of Wisconsin-Madison (United States), Cambodian Pasteur Institute (Cambodia), Hong-Kong University (China), Queen Mary Hospital (Hong-Kong, China), Chinese Center of Disease Control and Prevention (China), Vaksindo Satwa Nusantara (Indonesia), Umeå University (Sweden), National Public Health Laboratory (Luxembourg), WHO CC for Reference and Research on Influenza, Crick Institute London (United Kingdom), Friedrich-Loeffler Institut (Germany), Austrian Agency for Health and Food Safety, Mödling (Austria). We thank Hong-Kong University (China) and University of Wisconsin-Madison (United States) for providing cDNA/viruses used for the challenge experiments. We thank the Erasmus Laboratory Animal Science Center staff for animal care, and specifically Vincent Vaes and Dennis Akkermans for their assistance in animal experiments. We thank Peter van Run and Anja de Bruin for their help in processing the animal tissues for histology and immunohistochemistry, Djenolan van Mourik, Rachel Scheuer and Mark Pronk for their technical assistance, and David van de Vijver for input on statistical analyses. We thank Theo Luider and Lennard Dekker for performing the mass spectrometry experiments. We thank Sam Shepard (Center of Disease Control and Prevention, United States) for sharing a pre-released version of LABEL for A(H5) clade classification. We thank Gabriele Neumann and Yoshihiro Kawaoka (University of Wisconsin-Madison, USA) and their team for fruitful discussions and for providing the high-yield PR/8 backbone. Figure S8 was created with BioRender.com.

## Author contributions

Conceptualization: AK, DJS, RAM, MR

Methodology: AK, SHW, ST, SLJ, DFB, MF, DP, JvB, SH, MR

Investigation: AK, ST, SLJ, TB, DFB, MF, SvV, MIS, WR, DP, DdM, MER, PL, JvB, SH, MR

Visualization: AK, SHW, ST

Funding acquisition: DJS, RAMF, MR

Project administration: MR

Supervision: RAMF, MR

Writing – original draft: AK, MR

Writing – review & editing: AK, SHW, ST, SLJ, TB, DFB, MF, SvV, MIS, WR, DP, DdM, MER, PL, JvB, SH, DJS, RAMF, MR

## Data and materials availability

All data are available in the main text or the supplementary materials. Custom code used for data analysis is available at https://github.com/epiv-lab/H5-antigenic-evolution, https://github.com/epiv-lab/pepiniere and through Zenodo (*68, 69*). Data S1 to S10 are available via https://epiv-lab.github.io/H5-antigenically-central-vaccine/.

## References and Notes

1. S. W. Yoon, R. J. Webby, R. G. Webster, Evolution and ecology of influenza a viruses. Curr Top Microbiol Immunol 385, 359–375 (2014).

2. M. L. Shaw, P. Palese, “Orthomyxoviridae” in Fields Virology, D. M. Knipe, P. M. Howley, Eds. (Lippincott Williams & Wilkins, ed. 6, 2013), pp. 1152–1181.

3. U. Karakus, I. Mena, J. Kottur, S. S. El Zahed, R. Seoane, S. Yildiz, L. Chen, M. Plancarte, L. Lindsay, R. Halpin, T. B. Stockwell, D. E. Wentworth, G.-J. Boons, F. Krammer, S. Stertz, W. Boyce, R. P. de Vries, A. K. Aggarwal, A. García-Sastre, H19 influenza A virus exhibits species-specific MHC class II receptor usage. Cell Host Microbe 32, 1089–1102.e10 (2024).

4. X. Xu, K. Subbarao, N. J. Cox, Y. Guo, Genetic characterization of the pathogenic influenza A/Goose/Guangdong/1/96 (H5N1) virus: Similarity of its hemagglutinin gene to those of H5N1 viruses from the 1997 outbreaks in Hong Kong. Virology 261, 15–19 (1999).

5. J. C. de Jong, E. C. J. Claas, A. D. M. E. Osterhaus, R. G. Webster, W. L. Lim, A pandemic warning? Nature 389, 554–554 (1997).

6. R. Xie, K. M. Edwards, M. Wille, X. Wei, S. S. Wong, M. Zanin, R. El-Shesheny, M. Ducatez, L. L. M. Poon, G. Kayali, R. J. Webby, V. Dhanasekaran, The episodic resurgence of highly pathogenic avian influenza H5 virus. Nature 622, 810–817 (2023).

7. A. Bennison, A. M. P. Byrne, S. M. Reid, J. G. Lynton-Jenkins, B. Mollett, D. De Sliva, J. Peers-Dent, K. E. Finlayson, R. Hall, F. Blockley, M. Blyth, M. Falchieri, Z. Fowler, E. Fitzcharles, I. H. Brown, J. James, A. C. Banyard, Detection and spread of high pathogenicity avian influenza virus H5N1 in the Antarctic Region. bioRxiv, 2023.11.23.568045 (2023).

8. P. Kumar, A. Sharma, V. Apostolopoulos, A. M. Gaidhane, P. Satapathy, Australia’s first human case of H5N1 and the current H7 poultry outbreaks: implications for public health and biosecurity measures. Elsevier Ltd (2024). 10.1016/j.lanwpc.2024.101141.

9. FAO, Global Avian Influenza Viruses with Zoonotic Potential situation update (2024). https://www.fao.org/animal-health/situation-updates/global-aiv-with-zoonotic-potential/bird-species-affected-by-h5nx-hpai/en.

10. M. Ulloa, A. Fernández, N. Ariyama, A. Colom-Rivero, C. Rivera, P. Nuñez, P. Sanhueza, M. Johow, H. Araya, J. C. Torres, P. Gomez, G. Muñoz, B. Agüero, R. Alegría, R. Medina, V. Neira, E. Sierra, Mass mortality event in South American sea lions (Otaria flavescens) correlated to highly pathogenic avian influenza (HPAI) H5N1 outbreak in Chile. Veterinary Quarterly 43, 1–10 (2023).

11. V. Gamarra-Toledo, P. I. Plaza, R. Gutiérrez, G. Inga-Diaz, P. Saravia-Guevara, O. Pereyra-Meza, E. Coronado-Flores, A. Calderón-Cerrón, G. Quiroz-Jiménez, P. Martinez, D. Huamán-Mendoza, J. C. Nieto-Navarrete, S. Ventura, S. A. Lambertucci, Mass Mortality of Sea Lions Caused by Highly Pathogenic Avian Influenza A(H5N1) Virus. Emerg Infect Dis 29, 2553 (2023).

12. L. C. Caserta, E. A. Frye, S. L. Butt, M. Laverack, M. Nooruzzaman, L. M. Covaleda, A. C. Thompson, M. P. Koscielny, B. Cronk, A. Johnson, K. Kleinhenz, E. E. Edwards, G. Gomez, G. Hitchener, M. Martins, D. R. Kapczynski, D. L. Suarez, E. R. Alexander Morris, T. Hensley, J. S. Beeby, M. Lejeune, A. K. Swinford, F. Elvinger, K. M. Dimitrov, D. G. Diel, Spillover of highly pathogenic avian influenza H5N1 virus to dairy cattle. Nature, doi: 10.1038/s41586-024-07849-4 (2024).

13. Cumulative number of confirmed human cases for avian influenza A(H5N1) reported to WHO, 2003-2023, 24 April 2023, WHO (2023). https://www.who.int/publications/m/item/cumulative-number-of-confirmed-human-cases-for-avian-influenza-a(h5n1)-reported-to-who-2003-2023-24-april-2023.

14. Global influenza programme, Human-animal interface, Monthly risk assessment summary, WHO (2023). https://www.who.int/teams/global-influenza-programme/avian-influenza/monthly-risk-assessment-summary.

15. Avian Influenza Weekly Update Number 891, WHO (2023). https://www.who.int/docs/default-source/wpro---documents/emergency/surveillance/avian-influenza/ai_20230414.pdf?sfvrsn=5f006f99_113.

16. WHO, Summary of status of development and availability of A(H5N1) candidate vaccine viruses and potency testing reagents (2024). https://www.who.int/publications/m/item/a(h5n1)---northern-hemisphere-2024-2025.

17. WHO, Summary of status of development and availability of A(H5) non–A(H5N1) candidate vaccine viruses and potency testing reagents (2024). https://www.who.int/publications/m/item/a(h5)-non-a(h5n1)---northern-hemisphere-2024-2025.

18. R. B. Couch, W. K. Decker, B. Utama, R. L. Atmar, D. Niño, J. Q. Feng, M. M. Halpert, G. M. Air, Evaluations for In Vitro Correlates of Immunogenicity of Inactivated Influenza A H5, H7 and H9 Vaccines in Humans. PLoS One 7, 1–13 (2012).

19. S. S. Wong, J. DeBeauchamp, M. Zanin, Y. Sun, L. Tang, R. Webby, H5N1 influenza vaccine induces a less robust neutralizing antibody response than seasonal trivalent and H7N9 influenza vaccines. NPJ Vaccines 2, 1–8 (2017).

20. A. Kok, R. A. M. Fouchier, M. Richard, Cross-Reactivity Conferred by Homologous and Heterologous Prime-Boost A/H5 Influenza Vaccination Strategies in Humans: A Literature Review. Vaccines (Basel) 9 (2021).

21. D. J. Smith, A. S. Lapedes, J. C. De Jong, T. M. Bestebroer, G. F. Rimmelzwaan, A. D. M. E. Osterhaus, R. A. M. Fouchier, Mapping the antigenic and genetic evolution of influenza virus. Science (1979) 305, 371–376 (2004).

22. S. H. Wilks, Racmacs: R Antigenic Cartography Macros. version 1.1.35. https://acorg.github.io/Racmacs/.

23. D. F. Burke, D. J. Smith, A Recommended Numbering Scheme for Influenza A HA Subtypes. PLoS One 9, e112302 (2014).

24. W. Wang, B. Lu, H. Zhou, A. L. Suguitan, X. Cheng, K. Subbarao, G. Kemble, H. Jin, Glycosylation at 158N of the Hemagglutinin Protein and Receptor Binding Specificity Synergistically Affect the Antigenicity and Immunogenicity of a Live Attenuated H5N1 A/Vietnam/1203/2004 Vaccine Virus in Ferrets. J Virol 84, 6570–6577 (2010).

25. X. Zhang, S. Chen, Y. Jiang, K. Huang, J. Huang, D. Yang, J. Zhu, Y. Zhu, S. Shi, D. Peng, X. Liu, Hemagglutinin glycosylation modulates the pathogenicity and antigenicity of the H5N1 avian influenza virus. Vet Microbiol 175, 244–256 (2015).

26. A. Suttie, Y. M. Deng, A. R. Greenhill, P. Dussart, P. F. Horwood, E. A. Karlsson, Inventory of molecular markers affecting biological characteristics of avian influenza A viruses. Springer New York LLC (2019). 10.1007/s11262-019-01700-z.

27. FluSurver. http://flusurver.bii.a-star.edu.sg.

28. L. Xu, L. Bao, S. Y. Lau, W. L. Wu, J. Yuan, S. Gu, F. Li, Q. Lv, Y. Xu, P. Pushko, H. Chen, C. Qin, Hemagglutinin amino acids related to receptor specificity could affect the protection efficacy of H5N1 and H7N9 avian influenza virus vaccines in mice. Vaccine 34, 2627–2633 (2016).

29. S. Chutinimitkul, D. van Riel, V. J. Munster, J. M. A. van den Brand, G. F. Rimmelzwaan, T. Kuiken, A. D. M. E. Osterhaus, R. A. M. Fouchier, E. de Wit, In Vitro Assessment of Attachment Pattern and Replication Efficiency of H5N1 Influenza A Viruses with Altered Receptor Specificity. J Virol 84, 6825–6833 (2010).

30. M. E. Rosu, P. Lexmond, T. M. Bestebroer, B. M. Hauser, D. J. Smith, S. Herfst, R. A. M. Fouchier, Substitutions near the HA receptor binding site explain the origin and major antigenic change of the B/Victoria and B/Yamagata lineages. Proc Natl Acad Sci U S A 119, e2211616119 (2022).

31. M. Linster, E. J. A. Schrauwen, S. van der Vliet, D. F. Burke, P. Lexmond, T. M. Bestebroer, D. J. Smith, S. Herfst, B. F. Koel, R. A. M. Fouchier, The Molecular Basis for Antigenic Drift of Human A/H2N2 Influenza Viruses. J Virol 93 (2019).

32. A. Kok, R. Scheuer, T. M. Bestebroer, D. F. Burke, S. H. Wilks, M. I. Spronken, D. de Meulder, P. Lexmond, M. Pronk, D. J. Smith, S. Herfst, R. A. M. Fouchier, M. Richard, Characterization of A/H7 influenza virus global antigenic diversity and key determinants in the hemagglutinin globular head mediating A/H7N9 antigenic evolution. mBio 14 (2023).

33. B. F. Koel, S. van der Vliet, D. F. Burke, T. M. Bestebroer, E. E. Bharoto, I. W. W. Yasa, I. Herliana, B. M. Laksono, K. Xu, E. Skepner, C. A. Russell, G. F. Rimmelzwaan, D. R. Perez, A. D. M. E. Osterhaus, D. J. Smith, T. Y. Prajitno, R. A. M. Fouchier, Antigenic variation of clade 2.1 H5N1 virus is determined by a few amino acid substitutions immediately adjacent to the receptor binding site. mBio 5, 1–14 (2014).

34. Y. Wang, I. Davidson, R. Fouchier, E. Spackman, “Antigenic cartography of H9 avian influenza virus and its application to vaccine selection” in Avian Diseases (American Association of Avian Pathologists, 2016)vol. 60, pp. 218–225.

35. N. S. Lewis, J. M. Daly, C. A. Russell, D. L. Horton, E. Skepner, N. A. Bryant, D. F. Burke, A. S. Rash, J. L. N. Wood, T. M. Chambers, R. A. M. Fouchier, J. A. Mumford, D. M. Elton, D. J. Smith, Antigenic and Genetic Evolution of Equine Influenza A (H3N8) Virus from 1968 to 2007. J Virol 85, 12742–12749 (2011).

36. J. C. de Jong, D. J. Smith, A. S. Lapedes, I. Donatelli, L. Campitelli, G. Barigazzi, K. Van Reeth, T. C. Jones, G. F. Rimmelzwaan, A. D. M. E. Osterhaus, R. A. M. Fouchier, Antigenic and Genetic Evolution of Swine Influenza A (H3N2) Viruses in Europe. J Virol 81, 4315–4322 (2007).

37. N. S. Lewis, C. A. Russell, P. Langat, T. K. Anderson, K. Berger, F. Bielejec, D. F. Burke, G. Dudas, J. M. Fonville, R. A. M. Fouchier, P. Kellam, B. F. Koel, P. Lemey, T. Nguyen, B. Nuansrichy, J. S. Malik Peiris, T. Saito, G. Simon, E. Skepner, N. Takemae, R. J. Webby, K. Van Reeth, S. M. Brookes, L. Larsen, S. J. Watson, I. H. Brown, A. L. Vincent, S. Reid, M. A. Garcia, T. Harder, E. Foni, I. Markowska-Daniel, The global antigenic diversity of swine influenza A viruses. Elife 5 (2016).

38. G. Cattoli, A. Milani, N. Temperton, B. Zecchin, A. Buratin, E. Molesti, M. M. Aly, A. Arafa, I. Capua, Antigenic Drift in H5N1 Avian Influenza Virus in Poultry Is Driven by Mutations in Major Antigenic Sites of the Hemagglutinin Molecule Analogous to Those for Human Influenza Virus. J Virol 85, 8718–8724 (2011).

39. B. Peeters, S. Reemers, J. Dortmans, E. de Vries, M. de Jong, S. van de Zande, P. J. M. Rottier, • A. M. de Haan, Genetic versus antigenic differences among highly pathogenic H5N1 avian influenza A viruses: Consequences for vaccine strain selection. Virology 503, 83–93 (2017).

40. B. F. Koel, D. F. Burke, T. M. Bestebroer, S. Van Der Vliet, G. C. M. Zondag, G. Vervaet, E. Skepner, N. S. Lewis, M. I. J. Spronken, C. A. Russell, M. Y. Eropkin, A. C. Hurt, I. G. Barr, J. De Jong, G. F. Rimmelzwaan, A. D. M. E. Osterhaus, R. A. M. Fouchier, D. J. Smith, Substitutions near the receptor binding site determine major antigenic change during influenza virus evolution. Science (1979) 342, 976–979 (2013).

41. A. Pohlmann, J. King, A. Fusaro, B. Zecchin, A. C. Banyard, I. H. Brown, A. M. P. Byrne, N. Beerens, Y. Liang, R. Heutink, F. Harders, J. James, S. M. Reid, R. D. E. Hansen, N. S. Lewis, Hjulsager, L. E. Larsen, S. Zohari, K. Anderson, C. Bröjer, A. Nagy, V. Savic, S. van Borm, M. Steensels, F. X. Briand, E. Swieton, K. Smietanka, C. Grund, M. Beer, T. Harder, Has Epizootic Become Enzootic? Evidence for a Fundamental Change in the Infection Dynamics of Highly Pathogenic Avian Influenza in Europe, 2021. mBio 13 (2022).

42. G. Cattoli, A. Fusaro, I. Monne, F. Coven, T. Joannis, H. S. A. El-Hamid, A. A. Hussein, C. Cornelius, N. M. Amarin, M. Mancin, E. C. Holmes, I. Capua, Evidence for differing evolutionary dynamics of A/H5N1 viruses among countries applying or not applying avian influenza vaccination in poultry. Vaccine 29, 9368–9375 (2011).

43. C.-W. Lee, D. L. Suarez, Avian influenza virus: prospects for prevention and control by vaccination. Anim Health Res Rev 6, 1–15 (2005).

44. C. I. Paules, H. D. Marston, R. W. Eisinger, D. Baltimore, A. S. Fauci, The Pathway to a Universal Influenza Vaccine. Immunity 47, 599–603 (2017).

45. M. F. Ducatez, J. Bahl, Y. Griffin, E. Stigger-Rosser, J. Franks, S. Barman, D. Vijaykrishna, A. Webb, Y. Guan, R. G. Webster, G. J. D. Smith, R. J. Webby, Feasibility of reconstructed ancestral H5N1 influenza viruses for cross-clade protective vaccine development. Proc Natl Acad Sci U S A 108, 349–354 (2011).

46. H. Y. Liao, S. C. Wang, Y. A. Ko, K. I. Lin, C. Ma, T. J. R. Cheng, C. H. Wong, Chimeric hemagglutinin vaccine elicits broadly protective CD4 and CD8 T cell responses against multiple influenza strains and subtypes. Proc Natl Acad Sci U S A 117, 17757–17763 (2020).

47. M. W. Chen, T. J. R. Cheng, Y. Huang, J. T. Jan, S. H. Ma, A. L. Yu, C. H. Wong, D. D. Ho, A consensus - Hemagglutinin-based DNA vaccine that protects mice against divergent H5N1 influenza viruses. Proc Natl Acad Sci U S A 105, 13538–13543 (2008).

48. T. M. Ross, J. DiNapoli, M. Giel-Moloney, C. E. Bloom, K. Bertran, C. Balzli, T. Strugnell, M. Sá e Silva, T. Mebatsion, M. Bublot, D. E. Swayne, H. Kleanthous, A computationally designed H5 antigen shows immunological breadth of coverage and protects against drifting avian strains. Vaccine 37, 2369–2376 (2019).

49. D. J. Laddy, J. Yan, M. Kutzler, D. Kobasa, G. P. Kobinger, A. S. Khan, J. Greenhouse, N. Y. Sardesai, R. Draghia-Akli, D. B. Weiner, Heterosubtypic protection against pathogenic human and avian influenza viruses via In Vivo electroporation of synthetic consensus DNA antigens. PLoS One 3 (2008).

50. C. J. Crevar, D. M. Carter, K. Y. J. Lee, T. M. Ross, Cocktail of H5N1 COBRA HA vaccines elicit protective antibodies against H5N1 viruses from multiple clades. Hum Vaccin Immunother 11, 572–583 (2015).

51. B. M. Giles, T. M. Ross, A computationally optimized broadly reactive antigen (COBRA) based H5N1 VLP vaccine elicits broadly reactive antibodies in mice and ferrets. Vaccine 29, 3043–3054 (2011).

52. I. A. Nuñez, Y. Huang, T. M. Ross, Next-generation computationally designed influenza hemagglutinin vaccines protect against h5nx virus infections. Pathogens 10 (2021).

53. A. Kamlangdee, B. Kingstad-Bakke, T. K. Anderson, T. L. Goldberg, J. E. Osorio, Broad Protection against Avian Influenza Virus by Using a Modified Vaccinia Ankara Virus Expressing a Mosaic Hemagglutinin Gene. J Virol 88, 13300–13309 (2014).

54. R. J. Webby, E. A. Weaver, Centralized consensus hemagglutinin genes induce protective immunity against H1, H3 and H5 influenza viruses. PLoS One 10 (2015).

55. W. E. P. Beyer, A. M. Palache, A. D. M. E. Osterhaus, Comparison of serology and reactogenicity between influenza subunit vaccines and whole virus or split vaccines. A review and meta-analysis of the literature. Clin Drug Investig 15, 1–12 (1998).

56. C. Furey, N. Ye, L. Kercher, J. DeBeauchamp, J. C. Crumpton, T. Jeevan, C. Patton, J. Franks, M.-G. Alameh, S. H. Y. Fan, A. T. Phan, C. A. Hunter, R. J. Webby, D. Weissman, S. E. Hensley, Development of a nucleoside-modified mRNA vaccine against clade 2.3.4.4b H5 highly pathogenic avian influenza virus. bioRxiv, 1–13 (2023).

57. A. J. Mooney, J. D. Gabbard, Z. Li, D. A. Dlugolenski, S. K. Johnson, R. A. Tripp, B. He, S. M. Tompkins, Vaccination with Recombinant Parainfluenza Virus 5 Expressing Neuraminidase Protects against Homologous and Heterologous Influenza Virus Challenge. J Virol 91 (2017).

58. M. E. Rosu, A. Kok, T. M. Bestebroer, D. de Meulder, E. P. Verveer, M. R. Pronk, L. J. M. Dekker, T. M. Luider, M. Richard, J. M. A. van den Brand, R. A. M. Fouchier, S. Herfst, Contribution of Neuraminidase to the Efficacy of Seasonal Split Influenza Vaccines in the Ferret Model. J Virol 96 (2022).

59. I. M. Gilchuk, S. Bangaru, P. Gilchuk, R. P. Irving, N. Kose, R. G. Bombardi, N. J. Thornburg, C. B. Creech, K. M. Edwards, S. Li, H. L. Turner, W. Yu, X. Zhu, I. A. Wilson, A. B. Ward, J. E. Crowe, Influenza H7N9 Virus Neuraminidase-Specific Human Monoclonal Antibodies Inhibit Viral Egress and Protect from Lethal Influenza Infection in Mice. Cell Host Microbe 26, 715–728.e8 (2019).

60. M. J. Memoli, P. A. Shaw, A. Han, L. Czajkowski, S. Reed, R. Athota, T. Bristol, S. Fargis, K. Risos, J. H. Powers, R. T. Davey, J. K. Taubenberger, Evaluation of antihemagglutinin and antineuraminidase antibodies as correlates of protection in an influenza A/H1N1 virus healthy human challenge model. mBio 7 (2016).

61. M. R. Sandbulte, G. S. Jimenez, A. C. M. Boon, L. R. Smith, J. J. Treanor, R. J. Webby, Cross-reactive neuraminidase antibodies afford partial protection against H5N1 in mice and are present in unexposed humans. PLoS Med 4, 0265–0272 (2007).

62. P. Daulagala, S. M. S. Cheng, A. Chin, L. L. H. Luk, K. Leung, J. T. Wu, L. L. M. Poon, M. Peiris, H. L. Yen, Avian Influenza A(H5N1) Neuraminidase Inhibition Antibodies in Healthy Adults after Exposure to Influenza A(H1N1)pdm09. Emerg Infect Dis 30, 168–171 (2024).

63. E. Frobert, M. Bouscambert-Duchamp, V. Escuret, S. Mundweiler, M. Barthélémy, F. Morfin, M. Valette, C. Gerdil, B. Lina, O. Ferraris, Anti N1 cross-protecting antibodies against H5N1 detected in H1N1 infected people. Curr Microbiol 61, 25–28 (2010).

64. E. Hoffmann, A. S. Lipatov, R. J. Webby, E. A. Govorkova, R. G. Webster, Role of specific hemagglutinin amino acids in the immunogenicity and protection of H5N1 influenza virus vaccines. Proc Natl Acad Sci U S A 102, 12915–12920 (2005).

65. Q. Xu, W. Wang, X. Cheng, J. Zengel, H. Jin, Influenza H1N1 A/Solomon Island/3/06 Virus Receptor Binding Specificity Correlates with Virus Pathogenicity, Antigenicity, and Immunogenicity in Ferrets. J Virol 84, 4936–4945 (2010).

66. Z. Chen, H. Zhou, L. Kim, H. Jin, The Receptor Binding Specificity of the Live Attenuated Influenza H2 and H6 Vaccine Viruses Contributes to Vaccine Immunogenicity and Protection in Ferrets. J Virol 86, 2780–2786 (2012).

67. X. Sun, W. Cao, C. Pappas, F. Liu, J. M. Katz, T. M. Tumpey, Effect of receptor binding specificity on the immunogenicity and protective efficacy of influenza virus A H1 vaccines. Virology 464–465, 156–165 (2014).

68. A. Kok, S. Tureli, M. Richard, Epiv-lab/H5-antigenic-evolution: Archiving for initial submission. Zenodo v1.0 (2024). 10.5281/zenodo.13237525.

69. M. Funk, Epiv-lab/pepiniere: Archiving for initial submission. Zenodo v1.0 (2024). 10.5281/zenodo.12751132.

70. S. Elbe, G. Buckland-Merrett, Data, disease and diplomacy: GISAID’s innovative contribution to global health. Global Challenges 1, 33–46 (2017).

71. R. D. Olson, R. Assaf, T. Brettin, N. Conrad, C. Cucinell, J. J. Davis, D. M. Dempsey, A. Dickerman, E. M. Dietrich, R. W. Kenyon, M. Kuscuoglu, E. J. Lefkowitz, J. Lu, D. Machi, C. Macken, C. Mao, A. Niewiadomska, M. Nguyen, G. J. Olsen, J. C. Overbeek, B. Parrello, V. Parrello, J. S. Porter, G. D. Pusch, M. Shukla, I. Singh, L. Stewart, G. Tan, C. Thomas, M. VanOeffelen, V. Vonstein, Z. S. Wallace, A. S. Warren, A. R. Wattam, F. Xia, H. Yoo, Y. Zhang, C. M. Zmasek, R. H. Scheuermann, R. L. Stevens, Introducing the Bacterial and Viral Bioinformatics Resource Center (BV-BRC): a resource combining PATRIC, IRD and ViPR. Nucleic Acids Res 51, D678–D689 (2023).

72. K. Katoh, D. M. Standley, MAFFT Multiple Sequence Alignment Software Version 7: Improvements in Performance and Usability. Mol Biol Evol 30, 772–780 (2013).

73. B. Q. Minh, H. A. Schmidt, O. Chernomor, D. Schrempf, M. D. Woodhams, A. Von Haeseler, R. Lanfear, E. Teeling, IQ-TREE 2: New Models and Efficient Methods for Phylogenetic Inference in the Genomic Era. Mol Biol Evol 37, 1530–1534 (2020).

74. S. Kalyaanamoorthy, B. Q. Minh, T. K. F. Wong, A. von Haeseler, L. S. Jermiin, ModelFinder: fast model selection for accurate phylogenetic estimates. Nat Methods 14, 587–589 (2017).

75. B. Q. Minh, M. A. T. Nguyen, A. Von Haeseler, Ultrafast approximation for phylogenetic bootstrap. Mol Biol Evol 30, 1188–1195 (2013).

76. I. Letunic, P. Bork, Interactive tree of life (iTOL) v5: An online tool for phylogenetic tree display and annotation. Nucleic Acids Res 49, W293–W296 (2021).

77. G. Yu, D. K. Smith, H. Zhu, Y. Guan, T. T. Y. Lam, ggtree: an r package for visualization and annotation of phylogenetic trees with their covariates and other associated data. Methods Ecol Evol 8, 28–36 (2017).

78. S. S. Shepard, C. T. Davis, J. Bahl, P. Rivailler, I. A. York, R. O. Donis, LABEL: Fast and accurate lineage assignment with assessment of H5N1 and H9N2 influenza A hemagglutinins. PLoS One 9, 16–18 (2014).

79. E. Hoffmann, J. Stech, Y. Guan, R. G. Webster, D. R. Perez, Universal primer set for the full-length amplification of all influenza A viruses. Arch Virol 146, 2275–2289 (2001).

80. E. De Wit, M. I. J. Spronken, T. M. Bestebroer, G. F. Rimmelzwaan, A. D. M. E. Osterhaus, R. A. M. Fouchier, Efficient generation and growth of influenza virus A/PR/8/34 from eight cDNA fragments. Virus Res 103, 155–161 (2004).

81. E. de Wit, T. M. Bestebroer, M. I. J. Spronken, G. F. Rimmelzwaan, A. D. M. E. Osterhaus, R. A. M. Fouchier, Rapid sequencing of the non-coding regions of influenza A virus. J Virol Methods 139, 85–89 (2007).

82. S. Herfst, C. K. P. Mok, J. M. A. van den Brand, S. van der Vliet, M. E. Rosu, M. I. Spronken, Z. Yang, D. de Meulder, P. Lexmond, T. M. Bestebroer, J. S. M. Peiris, R. A. M. Fouchier, M. Richard, Human Clade 2.3.4.4 A/H5N6 Influenza Virus Lacks Mammalian Adaptation Markers and Does Not Transmit via the Airborne Route between Ferrets. mSphere 3, 1–17 (2018).

83. T. L. Williams, J. L. Pirkle, J. R. Barr, Simultaneous quantification of hemagglutinin and neuraminidase of influenza virus using isotope dilution mass spectrometry. Vaccine 30, 2475–2482 (2012).

84. P. D. Reuman, S. Keely, G. M. Schiff, Assessment of signs of influenza illness in the ferret model. J Virol Methods 24, 27–34 (1989).

85. M. Richard, J. M. A. van den Brand, T. M. Bestebroer, P. Lexmond, D. de Meulder, R. A. M. Fouchier, A. C. Lowen, S. Herfst, Influenza A viruses are transmitted via the air from the nasal respiratory epithelium of ferrets. Nat Commun 11, 1–11 (2020).

86. J. M. A. van den Brand, J. H. C. M. Kreijtz, R. Bodewes, K. J. Stittelaar, G. van Amerongen, T. Kuiken, J. Simon, R. A. M. Fouchier, G. Del Giudice, R. Rappuoli, G. F. Rimmelzwaan, A. D. M. E. Osterhaus, Efficacy of Vaccination with Different Combinations of MF59-Adjuvanted and Nonadjuvanted Seasonal and Pandemic Influenza Vaccines against Pandemic H1N1 (2009) Influenza Virus Infection in Ferrets. J Virol 85, 2851–2858 (2011).

87. J. M. A. Van Den Brand, K. J. Stittelaar, G. Van Amerongen, G. F. Rimmelzwaan, J. Simon, E. De Wit, V. Munster, T. Bestebroer, R. A. M. Fouchier, T. Kuiken, A. D. M. E. Osterhaus, Severity of pneumonia due to new H1N1 influenza virus in ferrets is intermediate between that due to seasonal H1N1 virus and highly pathogenic avian influenza H5N1 virus. Journal of Infectious Diseases 201, 993–999 (2010).

88. R. G. and S. D. H. Pagès, P. Aboyoun, Biostrings: Efficient manipulation of biological strings. R package version 2.62.0.

89. G. Sugiarto, K. Lau, J. Qu, Y. Li, S. Lim, S. Mu, J. B. Ames, A. J. Fisher, X. Chen, A sialyltransferase mutant with decreased donor hydrolysis and reduced sialidase activities for directly sialylating Lewisx. ACS Chem Biol 7, 1232–1240 (2012).

90. J. B. McArthur, H. Yu, J. Zeng, X. Chen, Converting Pasteurella multocida α2-3- sialyltransferase 1 (PmST1) to a regioselective α2-6-sialyltransferase by saturation mutagenesis and regioselective screening. Org Biomol Chem 15, 1700–1709 (2017).

91. S. H. Wilks, r3js: ‘WebGL’-Based 3D Plotting using the “three.js” Library. version 0.0.1 (2022). https://github.com/shwilks/r3js.

92. H. Wickham, “ggplot2: Elegant Graphics for Data Analysis” (Springer-Verlag New York, 2016; https://ggplot2.tidyverse.org).

93. C. Sievert, R. Iannone, J. Allaire, B. Borges, flexdashboard: R Markdown Format for Flexible Dashboards. R package version 0.6.0 (2022).

94. S. Tureli, PyRacmacs. (2024). https://github.com/iAvicenna/PyRacmacs.

95. M. Dawson-Haggerty, trimesh. version 3.2.0 (2020). https://trimsh.org/.

