## Supplementary materials for "A vaccine antigen central in influenza A(H5) virus antigenic space confers subtype-wide immunity"

**The PDF file includes:**

Materials and Methods

Supplementary Text S1 to S4

Figs. S1 to S11

**Other Supplementary Materials for this manuscript include the following:**

Movie 1

Tables S1 to S11

Data S1 to S10

Materials and Methods

Phylogenetic tree construction

All available A(H5) HA nucleotide sequences and accompanying metadata were downloaded from the Global Initiative on Sharing All Influenza Data (GISAID) (*70*) and the Bacterial and Viral Bioinformatics Resource Center (BV-BRC) (*71*) databases on 05/05/2023. The HA sequences of three antigens present in-house (A/Vietnam/3218/2004, A/duck/Hong-Kong/1091/2011 and A/eurasian-wigeon/Netherlands/EMC-3/2014), which were not yet available through the abovementioned databases at the time, were added to the dataset manually and deposited to GISAID in hindsight. The dataset was then preprocessed using Pépinière (a jupyter notebook available at [https://github.com/epiv-lab/pepiniere](https://eur01.safelinks.protection.outlook.com/?url=https%3A%2F%2Fgithub.com%2Fepiv-lab%2Fpepiniere&data=05%7C02%7Cm.richard%40erasmusmc.nl%7Cddde160cd5754a49d9c908dca5a48299%7C526638ba6af34b0fa532a1a511f4ac80%7C0%7C0%7C638567373204890746%7CUnknown%7CTWFpbGZsb3d8eyJWIjoiMC4wLjAwMDAiLCJQIjoiV2luMzIiLCJBTiI6Ik1haWwiLCJXVCI6Mn0%3D%7C0%7C%7C%7C&sdata=H%2FitIHT45Qjobx44Kh2GKO9WsQDevPAVfvEK6mvWR5Q%3D&reserved=0) and Zenodo (*69*)). This included deduplication of sequences present in both datasets based on identical accession numbers, identification and extraction of the open reading frame (ORF) corresponding to the longest ORF, and removal of sequences i) without metadata or ii) shorter than 90% of the mean ORF length. Identical sequences were then grouped, and only the earliest (by isolation date) representative was kept. After preprocessing, sequences were aligned using MAFFT v7.515 (*72*), and the alignment was trimmed to the start and stop codons of the majority of sequences. Trimmed sequences were again filtered to remove identical sequences (keeping the earliest). The resulting dataset contained 14896 sequences (table S1) and was realigned using MAFFT v7.515 and maximum-likelihood trees were generated using IQ-Tree2 (*73*) with the GTR+F+R10 model (chosen by ModelFinder (*74*)) and 10000 UFboot bootstrap approximations (*75*). Trees were midpoint rooted, annotated, and visualized using iTOL (*76*). The display item was generated using ggtree (*77*) in R. Genetic clades were predicted using LABEL (*78*) with the H5v2023 pre-release 1 (2023-05-05) module, courtesy of Sam Shepard, and this prediction was used to annotate the tree.

Cells

Cells were maintained as described previously (*32*). 293T cells (ATCC) were cultured in Dulbecco modified Eagle’s medium (DMEM) (Lonza) supplemented with 10% fetal calf serum (FCS) (Sigma-Aldrich), 1x non-essential amino acids (Lonza), 1 mM sodium pyruvate (Gibco), 2 mM L-glutamine (Lonza),100 U/mL of both penicillin and streptomycin (Lonza), and 0.5 mg/mL Geneticin (Invitrogen). Madin-Darby canine kidney (MDCK) cells (ATCC) were cultured in Eagle’s minimal essential medium (EMEM) (Lonza), supplemented with 10% FCS, 1x non-essential amino acids (Lonza), 1.5 mg/mL sodium bicarbonate (Lonza), 10 mM HEPES (Lonza), 2 mM L-glutamine (Lonza), and 100 U/mL of both penicillin and streptomycin (Lonza). Cells were cultured at 37 °C, 5% CO_2_, and passaged twice weekly.

Generation of plasmids

To generate plasmids for recombinant virus production, viral RNA was isolated from in-house available virus isolate stocks using the High Pure RNA Isolation Kit (Roche) according to the manufacturer’s instructions. The RNA was then used to generate viral copy DNA (cDNA) with the SuperScript IV Reverse Transcriptase (Thermo Fisher Scientific) according to the manufacturer’s instructions, from which individual HA or NA gene segments were amplified using segment-specific PCR primers (*79*) and the PfuUltra II Fusion HS DNA Polymerase (Agilent) according to the manufacturer’s instructions. Individual viral gene segments were cloned into a previously described modified pHW2000 plasmid (*80*) by restriction site-based cloning or seamless cloning using the GeneArt™ Seamless Cloning kit (Thermo Fisher Scientific). If the respective virus isolate was not present in-house, synthetic genes containing HA sequences with a monobasic cleavage site were synthesized by BaseClear or Integrated DNA Technologies. When applicable, specific mutations were introduced in the HA genes and /or the HA multibasic cleavage site (MBCS) was removed by site-directed mutagenesis using the PfuUltra II Fusion HS DNA Polymerase (Agilent) and specific primers. To produce reverse genetics plasmids for A(H5N6) A/Sichuan/26221/2014 (Sichuan, accession number: EPI_ISL_163493), all eight gene segments were amplified from cDNA by PCR using specific primers (*79*) and cloned into our bidirectional reverse genetics plasmid. To produce reverse genetics plasmids for A(H5N1) A/duck/Giza/15292S/2015 (Giza, accession number: EPI_ISL_257168), viral RNA was extracted, and all eight gene segments were amplified from cDNA by PCR using specific primers (*79*) and cloned into our bidirectional reverse genetics plasmid. Non-coding regions, which are not part of the abovementioned sequences, were sequenced upon RNA circularization as described previously (*81*)**.** The non-coding regions used are listed in table S11.

Benefit-sharing of synthetic constructs and viruses

Prior to the start of this work, we discussed and publicly announced our plans to generate synthetic hemagglutinin constructs and produce recombinant A(H5) viruses and A(H5)-specific ferret sera via the GISAID website (https://gisaid.org/collaborations/collaboration-on-h5-antigenic-cartography/). Specifically, we made the commitment that the synthetic HA constructs, reverse genetics viruses and ferret sera will be shared with the laboratories that contributed the genome sequence data to GISAID (*70*). We committed to publish the antigenic maps with open access to the public. We also indicated that reagents may be provided to other researchers, including National Influenza Centers and global reference laboratories, upon assurance that the originating laboratory, where the clinical specimen or virus isolate was first obtained, and the submitting laboratory, where sequence data have been generated and submitted through the GISAID mechanism, are fully recognized, to ensure fair attribution of contributions to the results benefitting from the data. We are indebted to GISAID and all scientists contributing to the GISAID initiative, without whom this work would not have been possible. We specifically wish to thank the governments and scientists of Austria, Bangladesh, Cambodia, China, Egypt, Germany, Ghana, Indonesia, India, Iraq, Japan, Nepal, Nigeria, Mongolia, Russia, Scotland, South Africa, Sweden, Turkey, United States and Vietnam for their contributions that made this research possible.

Biosafety

All experiments were reviewed by the Erasmus MC Institution Review Entity (IRE), in accordance with the U.S. Government September 2014 Dual-Use of Research of Concern (DURC) policy. The Erasmus MC IRE concluded that the studies described here were not falling under any of the seven DURC categories. Recombinant viruses which contained the HA (without MBCS) and/or NA of interest in the background of the attenuated vaccine strain A/Puerto Rico/8/1934 (PR/8) or the high yield version thereof (PR/8 HY, *76*) (table S2) were handled under biosafety level (BSL) 2 conditions in agreement with national regulations. Highly pathogenic avian influenza virus (HPAIV) wild-type isolates used in hemagglutination inhibition (HI) assays (Table S2) were handled under Biosafety level (BSL) 3 conditions. For the ferret challenge experiments under BSL3+ conditions, Sichuan and Giza recombinant viruses which contained all eight wild-type gene segments of a single virus were produced. These experiments were performed in the enhanced animal biosafety level 3 (ABSL3+) facility of the Erasmus University Medical Center as described previously (*82*).

Recombinant virus production

Recombinant influenza viruses were generated by reverse genetics using eight bidirectional plasmids as described previously (*32*, *80*). One day prior to transfection, about 3x10^6^ 293T cells were seeded in gelatin-coated 10 cm culture dishes. Calcium phosphate-mediated transfection was used to deliver a total of 40 μg of plasmid DNA per dish. About 16 hours after transfection, the cells were washed once with PBS and fresh media containing 2% FCS with 200-350 μg/mL N-tosyl-L-phenylalanine chloromethyl ketone (TPCK)-treated trypsin (Sigma-Aldrich) was added. Virus stocks were generated by inoculating either MDCK cells or 11-day old embryonated chicken eggs with dilutions of the supernatant harvested from the 293T cells three days post-transfection or virus isolates. Virus stock production in MDCK cells was performed using EMEM medium containing the same supplements as described above, but without FCS and with the addition of 20-35 μg/mL TPCK-treated trypsin, referred to as infection medium. MDCK supernatants or embryonated egg allantoic fluids were harvested two to three days post-inoculation and centrifuged at 2,100 *g* for 10 minutes to remove cellular debris. The presence of virus was confirmed by hemagglutination (HA) assays using 1% turkey red blood cells (TRBCs, from in-house turkeys) in PBS. Sequences from all plasmids and from the non-PR/8 and PR/8 HY genes, i.e. HA and NA of interest, of all virus stocks were confirmed with Sanger sequencing using the BigDye™ Terminator v3.1 Cycle Sequencing Kit (Applied Biosystems) and the 3500xL Genetic Analyzer (Applied Biosystems).

Virus titrations

Virus titrations were performed in MDCK cells as described previously (*82*). Briefly, flat-bottom 96 wells plates containing confluent MDCK cells were inoculated with 10-fold serial dilutions of the samples and incubated for one hour at 37 °C, 5% CO_2_. Cells were washed once with PBS and 200 μL of infection medium was added to each well. After three days of incubation at 37 °C, 5% CO_2_, the presence of virus in the supernatants was determined using HA assays to determine the 50% tissue culture infectious dose (TCID_50_). Virus titrations of the virus stocks were performed in ten replicates, and those of respiratory swabs and tissue homogenates from the vaccination-challenge experiments in four replicates.

Vaccine production

Vaccines were produced with recombinant viruses containing an engineered or wild-type HA, without MBCS, in the PR/8 HY background. For the initial screen of CVAs whole-inactivated vaccines were generated with the corresponding matched NA. For the vaccination-challenge experiment, split-inactivated vaccines were used. To isolate the effect of varying the HA antigen in these studies, the neuraminidase present in the vaccine was mismatched with that of the challenge virus. Specifically, the vaccines in the Giza challenge contained the N6 NA of Sichuan, and the vaccines in the Sichuan challenge contained the N1 NA of Giza.

Whole-inactivated and split-inactivated vaccines were generated as described previously (*58*). Eleven-day old embryonated chicken eggs were inoculated with the virus of interest. Allantoic fluid was harvested two days post-inoculation and centrifuged for ten minutes at 2,100 *g* to remove cellular debris. Subsequent centrifugation steps were performed at 124,000 *g* (SW 32 Ti, Beckman Coulter) at 4 °C, unless indicated otherwise. The allantoic fluid was concentrated on a 60% sucrose cushion by centrifuging for 2 hours. Subsequently, resuspended sucrose cushions from multiple tubes were pooled and loaded on 60-50-40-30-20% sucrose gradients, which were centrifuged overnight at the lowest deceleration setting. The virus band, located on top of the 30% sucrose layer, was harvested, diluted in PBS, and subsequently pelleted by centrifugation for 2 hours to remove the sucrose. The pellet was dissolved in either PBS (whole-inactivated vaccines) or PBS with 2% Mega10 (Sigma-Aldrich) (split-inactivated vaccines). Incubation with PBS with 2% Mega10 was performed for one hour at 37 °C. For both whole- and split-inactivated vaccines, the dissolved pellets were transferred to dialysis chambers (Slide-A-Lyzer™ Dialysis Cassettes, 10K MWCO, Thermo Fisher Scientific) which were subsequently submerged in PBS containing 0.01% formalin for three days. Subsequently, the dialysis chambers were immerged in PBS for a day, during which the PBS was refreshed twice. The resulting vaccines were aliquoted and stored at -80 °C. Vaccine inactivation was confirmed by two serial blind passages on MDCK cells and/or in embryonated chicken eggs.

Total protein content was determined using the Pierce BCA total protein analysis kit (Thermo Fisher Scientific). For the whole-inactivated vaccines, the absolute and relative HA content was estimated from SDS-PAGE protein gels using a bovine serum albumin (BSA) standard and stained with instant Blue (Expedeon). The absolute HA content of the split-inactivated vaccines was estimated from SDS-PAGE using a BSA standard and stained with instant Blue (Expedeon). The relative HA content of split-inactivated vaccines was determined with mass spectrometry , using a protocol based on Williams et al. (*83*) with modifications as described previously (*58*). Diluted vaccines (10 μL of 125 μg/mL of total protein) were mixed 1:1 with 0.2% RapiGest (Waters), and denatured for 5 minutes at 100 °C. After cooling to room temperature, 5 μL of sequence-grade modified trypsin solution (0.4 μg/μL; Promega) was added, and samples were incubated at 37 °C for 2 hours. Digests were allowed to cool, and 55 μL of 0.5% trifluoroacetic acid was added. The samples were subsequently analyzed by a nano liquid chromatography (LC) Ultimate 3000 system (Thermo Fisher Scientific) coupled to the Orbitrap Fusion Lumos mass spectrometer (Thermo Fisher Scientific).

Data from initial screens was used to select three peptides for stable isotope (SI) labelling (LVLATGLR, VNSIIDK and TLDFHDSNVK), based on intensity, length, sequence, and sequence conservation within the A(H5) HA subtype. Digested vaccines were spiked with SI labeled peptides with heavy lysine or arginine (Pepscan) at a final concentration of 50 fmol/μL, and measured on a nano-LC system (Ultimate 3000; Thermo Fisher Scientific) combined with an Orbitrap Fusion Eclipse Tribrid mass spectrometer (Thermo Fisher Scientific). For each peptide, the ratio between the SI labeled peptide and the endogenous peptide was calculated, which was subsequently used to determine the concentrations of the endogenous peptide in the vaccines. The HA concentration based on the three individual peptides was averaged for each vaccine, and subsequently used to determine the relative HA content of the vaccines.

Ferret experiments

Ferret experiments were performed in strict compliance with the Dutch legislation on the protection of animals used for scientific purposes (2014, European Union directive 2010/63/EU implemented). Experiments were performed at the Erasmus Medical Center in Rotterdam, the Netherlands under a project license accredited by the Dutch competent authority (license number AVD101002015340). Study protocols were approved by the Erasmus Medical Center Animal Welfare Body (permit numbers 15-340-01, -04, -06, -22, -23, and -24). Ferrets were seronegative for Aleutian disease, seasonal influenza A(H1N1), A(H3N2), and B viruses.

*Serum production*

Ferret sera were generated as described previously (*32*, *33*) in class III isolators under BLS3 conditions using recombinant viruses unless indicated otherwise (table S2, S3). Recombinant viruses were produced carrying the HA (without MBCS) and the closest matching NA present in-house, in the background of PR/8 or PR/8 HY. Briefly, male ferrets were inoculated intranasally by applying dropwise 250 μL of virus stock per nostril. Unless indicated otherwise (table S2), a boost was administered after 14 days, by subcutaneously injecting a total of 250 μL concentrated virus combined with 250 μL TiterMax® Gold adjuvant (Sigma-Aldrich) at two different spots in the back.

The concentrated virus used for the subcutaneous boost was prepared by inoculating five 11-day old embryonated chicken eggs per virus. The allantoic fluid was harvested two days post-inoculation and cleared from debris by centrifuging for 10 minutes at 2,100 *g*. About 36 mL of the cleared allantoic fluid was concentrated by centrifuging for 2 hours at 124,000 *g* (SW 32 Ti, Beckman Coulter), and the resulting pellet was resuspended in 700 μL PBS. Ferrets were terminally bled 14 days after the subcutaneous boost, or 14 days after the intranasal inoculation if no boost injection was performed. Before virus inoculation, subcutaneous boost injection, and the terminal bleed, ferrets were anesthetized with ketamine and medetomidine (10 and 0.05 mg/kg body weight, respectively), the latter which was antagonized with atipamezole (0.25 mg/kg body weight).

The blood samples were collected in VACUETTE® CAT Serum Separator Clot Activator tubes (Greiner Bio-One), incubated at least for 15 minutes to allow clotting, and centrifuged for 15 minutes at 2000 *g* to obtain the serum.

*Vaccination(-challenge) studies*

Vaccination experiments were performed similarly as described previously (*58*). Female ferrets were vaccinated twice intramuscularly on day 0 (prime) and day 28 (boost) with 250 μL of whole or split-inactivated vaccine estimated to contain about 7.5 μg HA, adjuvanted with 250 μL AddaVax™ (InvivoGen), which was equally divided between the two hind legs. For the mock-vaccinated groups, animals were vaccinated with 250 μL of PBS adjuvanted with 250 μL AddaVax™. Prior to each vaccination, a blood sample was obtained through the cranial vena cava and serum was obtained as described above (pre- and pre-boost sera). The pre-sera were tested in HI assays as described below against seasonal influenza A(H1N1), A(H3N2), and B viruses (using vaccine strains of the respective year), as well as PR/8 recombinant viruses carrying A(H5) HAs from the respective vaccines and challenge virus, if applicable. Pre-sera were negative in HI assays against the tested viruses. The pre-boost sera of the ferrets from the challenge experiments were titrated in HI assays against PR/8 recombinant viruses with three vaccine antigens employed in the respective study.

For vaccination-only experiments, ferrets were sacrificed 28 days after the boost vaccination through a cardiac puncture, and post-boost sera were obtained from whole blood as described above.

For vaccination-challenge experiments, DST micro-T temperature loggers (Star-Oddi) were surgically implanted in the abdominal cavity of the ferrets 14 days after the prime vaccination. Serum samples were collected from whole blood sampled via the cranial vena cava one week before inoculation (post-boost sera) (fig. S8, table S6, S7). Vaccinated ferrets were then transferred to class III isolators for acclimatization a week prior to the challenge virus inoculation. Ferrets were inoculated intranasally and intratracheally with wild-type recombinant viruses containing all eight segments of the respective challenge virus. The inoculation doses were 10^5.5^ and 10^3.4^ TCID_50_ per animal for the Giza and Sichuan virus, respectively, divided over 3 mL intratracheally and 250 μL in each nostril. These doses were determined prior to the challenge in a pilot experiment using three ferrets per tested dose. The doses were selected to induce a reproducible and consistent infection of the upper and lower respiratory tracts. Subsequently, daily nose and throat swabs were collected under light ketamine anesthesia, and body weight and activity level score were monitored daily as described previously (*82, 84*). Body temperature was recorded every ten minutes by the implanted temperature loggers. Four days post-inoculation, ferrets were euthanized through cardiac puncture, after which tissues (selected based on virus detection in the pilot studies) were collected for virological and pathological analysis as described previously (*82*). For virological analysis, the right nasal turbinates, trachea, right bronchus, right lung lobes, tracheobronchial lymph node, and liver (for both challenges), right cerebrum and right cerebellum (for the Giza challenge only), and the spleen (for the Sichuan challenge only) were collected. For pathological examination, left nasal turbinates, trachea, left bronchus, and left lung lobes were collected.

During blood collection, vaccination, virus inoculation, and cardiac puncture, ferrets were anesthetized with a mixture of ketamine and medetomidine, and antagonized with atipamezole, as described above.

Histopathology and immunohistochemistry

Upon necropsy, tissues were stored in 10% neutral-buffered formalin (lungs after careful inflation with formalin) for at least two weeks, after which the tissues were embedded in paraffin. Four-μm slides were made, and subsequent slides were either stained with hematoxylin and eosin (HE) or used for immunohistochemistry as described previously (*85*). Briefly, after deparaffinization, antigen retrieval, and blocking of endogenous proteases, slides were incubated for one hour at room temperature with either a primary antibody against influenza A virus nucleoprotein (Clone Hb65, American Type Culture Collection) or a mouse IgG2a isotype control (R&D), diluted in PBS with 0.1% BSA (1:400 and 1:200, respectively). After three washes with PBS with 0.05% Tween 20, slides were incubated for one hour at room temperature with a goat anti-mouse IgG2a secondary antibody coupled to horseradish peroxidase (HRP) (Biorad, Star133P), diluted 1:100 in PBS with 0.1% BSA. HRP was revealed using 3-Amino-9-Ethylcarbazole and a hematoxylin counterstain was performed. A lung section from an animal experimentally infected with 2009 pandemic A(H1N1) virus was used as positive control in each staining experiment.

The pathological changes and the presence of viral antigen in respiratory tissues were blindly scored in a semi-quantitative fashion by a veterinary pathologist. Semi-quantitative assessment of influenza virus-associated inflammation in the lungs (four slides with longitudinal section and cross-section of cranial and caudal lobes per animal) was performed blindly on every slide as reported earlier (*86*). The extent of alveolitis and alveolar damage was scored as follows: 0: 0%; 1: 1–25%; 2: 25–50%; 3: >50%. The severity of alveolitis, bronchiolitis, bronchitis, bronchial adenitis, tracheitis, and rhinitis were scored as follows: 0: no inflammatory cells; 1: few inflammatory cells; 2: moderate numbers of inflammatory cells; 3: many inflammatory cells. The presence of alveolar edema, alveolar hemorrhage and type II pneumocyte hyperplasia were scored as follows: 0: no; 1: yes. Finally, the extent of peribronchial, peribronchiolar, and perivascular infiltrates were scored as follows: 0: none; 1: one to two cells thick; 2: three to ten cells thick; 3: more than ten cells thick. Semi-quantitative assessment of influenza virus antigen expression in the lungs was performed as reported earlier (*87*). For the alveoli, twenty-five arbitrarily chosen fields of lung parenchyma of the four lung sections per animal were blindly examined by light microscopy, using a 20x objective, for the presence of influenza virus nucleoprotein. The cumulative scores for each animal were presented as percentage corresponding to the number of positive fields. The percentage of positive epithelium in the bronchi and bronchioles was estimated on all four lung slides and averaged per animal. The percentage of positively staining epithelium in the nose and trachea was estimated for one slide.

Serological assays

HI assays were performed with recombinant viruses in PR/8 or PR/8 HY background and virus isolates (Table S2) as described previously (*32*) using in-house TRBCs. Sera were treated overnight at 37 °C with five volumes of a *Vibrio cholerae* filtrate (generated in-house) containing receptor-destroying enzyme (*RDE*), to prevent aspecific inhibition. After inactivation for one hour at 56 °C, sera were adsorbed using an equal volume of 10% TRBCs for one hour at 4 °C, to prevent aspecific agglutination. Two-fold serial dilutions of sera in PBS were prepared in round-bottom 96-wells plates starting at 1:20 in a volume of 50 μL. To each well, 25 μL of virus, diluted in PBS to 4 hemagglutinating units (HAU), was added. After incubation for 30 minutes at 37 °C, 25 μL of 1% TRBCs was added to each well. Plates were subsequently incubated for one hour at 4 °C before reading the HI titer. The HI titer was determined as the reciprocal value of the highest serum dilution which completely inhibited TRBC agglutination. For the calculation of geometric mean titers (GMTs), threshold titers of <10 were converted to 5 unless stated otherwise.

VN assays were performed in MDCK cells as described previously (*32*, *40*). Firstly, sera were incubated for 30 minutes at 56 °C to inactivate complement. Two-fold serial dilutions of sera in PBS, starting at 1:10, were combined with 100 TCID_50_ of virus, and incubated for 2 hours at 37 °C. Subsequently, the virus-serum mixtures were added to flat-bottom 96-wells plates containing confluent MDCK cells previously washed once with PBS. After incubation for 2 hours at 37 °C and 5% CO_2_, cells were washed once with PBS, and 200 μL infection media per well was added. Plates were incubated at 37 °C, 5% CO_2_, and the presence or absence of virus in supernatants was determined after three days using HA assays with TRBCs. The VN titer was determined as the reciprocal value of the highest serum dilution for which no virus in supernatants detected was detected. VN assays were performed in duplicate, and the arithmetic means of logarithm base 2 (log_2_) titers were calculated.

Antigenic cartography and antibody profiles

Antigenic maps were constructed from HI data using a multidimensional scaling algorithm as described previously (*21*) using the R package ‘Racmacs’, version 1.2.3 (*22*). Firstly, HI titers are converted to a distance matrix (HI table distances) by (i) dividing each HI titer by 10 and applying a log_2_ transformation (hereafter defined as log_2_ transformed HI titers), and (ii) subtracting each log_2_ transformed titer to the highest one for each serum. Secondly, multidimensional scaling algorithms are used to find the best set of map coordinates to represent the distances from the distance matrix most closely. For each optimization, antigens and sera, hereafter named points, are randomly placed in n-dimensional space, and coordinates are optimized from these starting conditions using the L-Broyden-Fletcher-Goldfarb-Shanno (L-BFGS) algorithm, minimizing the sum of the squared differences between HI table distances and the map distances (Euclidian distance between points in the n-dimensional space). In an antigenic map, every direction represents antigenic distance, and one antigenic unit (AU) corresponds to a 2-fold change in HI titer.

Unless stated otherwise, antigenic maps were computed using the ‘make.acmap’ function, with 1000 optimization runs in three dimensions, and the minimum column basis set to zero. The antigenic map was validated using several tests which are described in the supplementary text. Total map stress was extracted using the ‘mapStress’ function and individual antigen stresses were extracted using the ‘agStress’ function. HI table distances and pairwise antigen-serum Euclidian distances in the antigenic map were extracted using the ‘tableDistances’ and ‘mapDistances’ functions, respectively. When threshold HI titers (i.e., <10) are converted to table distances in the process of making an antigenic map, the resulting values are not an exact distance but a ‘greater-than’ value, i.e. thresholded distance. To include these points in the visualization in scatter plots of HI table distances versus map distances (fig. S3A), and corresponding regression coefficient (R^2^) calculations (fig. S1C, S3A), these values were converted to the thresholded distance increased by 1 on the log_2_ scale, e.g. a threshold distance of >7 on the log_2_ scale is converted to an 8. To compute distances between points in the map, antigen and serum coordinates were extracted using the ‘agCoords’ and ‘srCoords’ functions, respectively. The ‘dist’ function (base R) was subsequently used to compute pairwise Euclidean distances between points in the map. The map center was determined by computing the mean x, y and z of the antigen coordinates. Pairwise genetic hamming distances, i.e. the number of amino acid differences between two antigens, were computed using the ‘stringDist’ function (method = ‘hamming’) from the Biostrings package (*88*).

To average and visualize the immune response of multiple ferrets belonging to the same experimental group, the GMTs of multiple vaccination sera against each individual antigen were computed to generate antibody profiles. Individual threshold titers were first converted to the closest possible numerical titer (e.g., <10 to a 5), and subsequently, GMTs were calculated. For the visualization of the reactivity of post-vaccination sera using the A(H5) antigenic map, antigenic maps were optimized with datasets containing the antigenic map HI data and HI data of a single post-vaccination serum or mean data of multiple post-vaccination sera as described above. The resulting maps including individual or mean post-vaccination sera data shared similar conformations, also corresponding to that of the antigenic map (the mean median Procrustes distance between each map with post-vaccination data and the antigenic map was 0.14 AU). Therefore, to generate displays in which the positions of post-vaccination sera were visualized without changing the position of the antigens or sera in the antigenic map, maps, containing either individual or merged HI data, were superimposed on the antigenic map using the ‘mergeMaps’ function with the ‘frozen-merge’ method. These maps were then used to visualize and analyze the reactivity of the post-vaccination sera to antigens in the antigenic map using custom R code (<https://github.com/epiv-lab/H5-antigenic-evolution> and Zenodo (*68*)). For analysis of mutant antigens, antigenic maps were computed with datasets containing HI data of a single mutant antigen in addition to the antigenic map dataset. The resulting optimized maps were used to calculate the distances described in the text. For visualization in supplementary fig. S4B, superimposed maps were generated as described above for the visualization of post-vaccination sera.

Resialylated turkey red blood cell (TRBC) assay

Resialylated TRBC assays were performed as described previously (*82*). The pellet of 1.25 mL of 1% TRBCs was resuspended in 62.5 μL PBS and incubated for 1 hour at 37 °C with 50 μL of 1 mU/μL *Vibrio cholerae* neuraminidase (VCNA) (Roche) and 10 μL 0.1 M CaCl_2_ to remove all sialic acids from the TRBCs. After two washes with PBS, the TRBCs were combined with 3.75 μL of 30 mM CMP-sialic acid (Merck), and either 5 μL of α2,3-sialyltransferase (Recombinant Human ST3GAL6, Fc Chimera, R&D systems) or 5 μL α2,6-sialyltransferase (Recombinant Human ST6GAL1 (aa 44-406) Protein, R&D systems), and PBS up to 75 μL. Alternatively, 1 μg of in-house generated *Pasteurella multocida* sialyltransferase 1 (Pmst1) M144D (α2,3-sialyltransferase) (*89*) or Pmst1 M144L P34H (α2,6-sialyltransferase) (*90*) was used. TRBCs were incubated at 37 °C for two hours with the commercial enzymes or for four hours with the in-house generated enzymes. After re-sialylation, TRBCs were washed twice with PBS, and resuspended in PBS containing 1% BSA to a final concentration of 0.5% TRBCs. Besides α2,3- and α2,6-sialic acid specific TRBCs, untreated, and VNCA-treated TRBCs were taken along as controls when assessing the binding preference of viruses in HA assays. Moreover, in each independent assay, a minimum of three control viruses was used: one with α2,3 sialic acid specificity, one with α2,6-sialic acid specificity, and one with dual binding specificity.

Data visualization and statistics

Data were visualized with Racmacs (*22*), r3js (*91*) and/or ggplot2 (*92*) in R. The supplementary interactive figure files (Data S2-S10) were generated with flexdashboard (*93*) in R. Statistical analyses were performed using the base R functions for the Kruskal-Wallis test, the pairwise Mann-Whitney/Wilcoxon test, and linear regressions. For the comparison of multiple experimental groups, the Kruskal-Wallis test was first performed. If positive, a pairwise Mann-Whitney test was then performed to assess the significance of differences between two experimental groups.

Supplementary Text

Supplementary text S1 - Compilation of HI dataset for the A(H5) antigenic map

The A(H5) map dataset was generated using HI data from multiple individual assays. The 127 antigens titrated against 33 post-infection sera resulted in a total of 4191 datapoints. Each combination of antigen and serum was titrated 1.7 times on average, and 48% of antigen-serum pairs were titrated two or more times. The standard deviation (SD) of log_2_ transformed HI titers between assays was below 1.5 for 93% of the data points which were titrated two or more times, with an average of 0.46 (fig. S1A). Of note, one AU difference corresponds to an SD of 0.7, meaning that differences of one or two AUs represent most of the observed HI assay variation. The HI data of individual assays were compiled into a merged dataset through the ‘mergeMaps’ function in Racmacs, using the ‘table’ method and the merge options settings of method = ‘conservative’ and sd_limit = 1.5. The sample SD of 1.5 allowed for the merging of titers within a four-fold difference, which is generally considered as the maximum acceptable variation between HI assays. Titers which were outside of the assay’s detection limit, denoted with a ‘smaller-than’ or ‘larger-than’ sign, were converted to the closest numerical titer (e.g., a <10 is converted to a 5) for calculation of the sample SD. Generally, for each antigen-serum combination, the resulting merged titer was the geometric mean of all measured values. In the case of threshold titers, if the SD was below the set limit of 1.5, the merged titer was the highest ‘smaller-than’ or the lowest ‘larger-than’ which satisfied all the measured values. For 145 antigen and serum combinations (3.46% of the 4191 antigen and serum combinations in the full dataset), the sample SD of the titration repeats was above 1.5, and consequently the titer was set to ‘NA’ and considered as unknown for the generation of the antigenic map. The resulting merged dataset was used as a basis for the generation of the antigenic map (table S4).

Supplementary text S2 - A(H5) antigenic map dimensionality

Firstly, we investigated the number of dimensions required to best represent the A(H5) HI data in an antigenic map. Generally, for a map of *n* dimensions, each point should have a minimum of *n* detectable titers (i.e., non-threshold and non-‘NA’ titers) to be placed in the antigenic map, and a minimum of *n*+1 detectable titers for accurate placement. In the cross-HI dataset, 13 antigens had less than five detectable titers (table S4). To allow for a fair comparison between dimensions one to five, we excluded these antigens and their five corresponding homologous sera, resulting in a dataset with 114 antigens and 28 sera (114x28, table S2).

A dimension test was performed using the ‘dimensionTestMap’ function in Racmacs. For dimensions one to five, 100 antigenic maps were generated from 100 random start positions, each of them performed with 1000 optimizations. In each of these 100 repeats, 10% of the data were randomly excluded, and subsequently predicted from the resulting map. For each repeat, the root mean square error (RMSE) between predicted and measured titers was calculated. A comparison of the mean RMSE in each dimension is indicative of the dimension required to represent the data well (fig. S1B).

The largest improvement in mean RMSE was observed between dimensions one and two (mean RMSE difference of 0.47), followed by dimensions two and three (mean RMSE difference of 0.29), suggesting that three dimensions represent the HI data substantially better (fig. S1B). A slight improvement was observed between three and four dimensions (mean RMSE difference of 0.11), and barely any improvement between four and five dimensions (mean RMSE difference of 0.01), indicating that more than four dimensions did not substantially improve representation of the data (fig. S1B). A comparable pattern was observed when comparing the correlation between pairwise antigen-serum HI table distances and pairwise antigen-serum Euclidian distances in the antigenic map across dimension one to five. The correlation improved only slightly between three and four dimensions (R^2^ of 0.66 to 0.71), as compared to between one, two and three dimensions (R^2^ of 0.24, 0.51 and 0.66, respectively) (fig. S1C). Similarly, the reduction in the total map stress, corresponding to the sum of the squared differences between table and map distances, was only minimal between three and four dimensions (3875 to 3174), as compared to that between one, two and three dimensions (12276, 5920 and 3875, respectively) (fig. S1D).

We further investigated whether the A(H5) cross-HI data should be represented in three or four dimensions. The differences in point positions between two maps were compared using a Procrustes analysis through the ‘procrustesData’ function in Racmacs. Given two antigenic maps with identical antigens, sera, and dimensions, the Procrustes function aims to find the optimal Euclidean transformation (translation, rotation, scaling, reflection) of the first map such that the distance between the coordinates of the two maps (called the Procrustes error) is minimized, i.e., the best superimposition is determined. The resulting difference in map coordinates for each point is the Procrustes distance, expressed in AU. This analysis can be visualized within the antigenic map (with the ‘procrustesMap’ function in Racmacs), where the point positions of the first map are visualized as usual, and arrows point to the position of the corresponding points in the comparison map. The length of an arrow corresponds to the Procrustes distance between the positions of a point in the two maps. Total Procrustes distances between two maps can then be expressed by calculating the root mean square (RMS) or the median Procrustes distance. When dimensions are not identical, one can supplement the lower dimensional map with extra coordinates of zero. The resulting map comparison indicates how the extra dimension is utilized to better fit the observed titers in the larger dimensional map, or, how the larger dimensional map is projected on to the lower dimensional map. Little difference was observed in the point positions between the three- and four-dimensional maps (RMS of 2.40 AU and median Procrustes distance of 1.55 AU). Moreover, the distances between antigens and sera in the map, and the individual antigen stresses correlated well between the three- and four-dimensional maps (fig. S1E, F, R^2^ of 0.93 and 0.84, respectively).

To further compare the three- and four-dimensional maps, a piecewise Procrustes analysis was performed. To this end, we used a modified version of the Procrustes analysis to compare maps of different dimensions, which is available in PyRacmacs (*94*) (code available via <https://github.com/epiv-lab/H5-antigenic-evolution> and Zenodo (*68*)). Given two maps and *n* the number of pieces, piecewise Procrustes aims to find the optimal partition of the coordinates into *n* pieces such that the sum of Procrustes errors is minimized. In essence, the different pieces of the map are allowed to move freely relative to each other when transitioning between dimensions. If one can obtain a lower Procrustes error between a lower and higher dimensional map when using few pieces, the lower dimensional map can be embedded in the higher dimensions by a relatively simple transformation such that it would look like the higher dimensional map, i.e. the two maps are geometrically similar. Here, we have used this method to compare the antigenic map in three and four dimensions. One can see that the three-dimensional map fits the four-dimensional map quite well when split into only two pieces (fig. S2, Data S3A, B). These analyses suggest that the general topology of the map in four-dimensions is similar to that of the three-dimensional map.

Given that the three-dimensional map recapitulated the geometry of the four-dimensional map well, and that only minimal improvements in prediction error, correlation between HI table and antigenic map distances, and overall stress were observed upon the use of the fourth dimension, we concluded that its use to visualize A(H5) cross-HI data was not warranted. However, it is not excluded that four dimensions will be required with further expansion of the A(H5) map including novel antigens and sera.

Supplementary text S3 - Analysis of three-dimensional A(H5) antigenic map

The dataset initially contained 10 antigens that reacted with fewer than four detectable titers to ferret sera, insufficient to confidently place points in a map of three or more dimensions. This was despite the presence of homologous sera for four of them (A/chicken/West-Java/119/2010 (clade 2.1.3.2a), A/duck/Jiangxi/0114-NCJD064-P/2015 (clade 2.3.4.4f), A/Guandong/18SF020/2018, (clade 2.3.4.4h), A/chicken/Chiping/0321/2014 (clade 7.2)), which were raised in an attempt to better characterize low reactive antigens. Homologous titers against these ferret sera were high, ranging between 392 and 1580, indicating that these four antigens have divergent antigenic properties, rather than being generally low reactive in HI assays. However, the possibility of the other six viruses being low reactive rather than antigenically divergent cannot be excluded without the presence of a homologous serum. To allow placement in the three-dimensional antigenic map, these antigens and, if available, corresponding homologous sera, were removed from the dataset (table S2, S3), but raw HI reactivity patterns are shown in Table S4. The final dataset used to generate the A(H5) three-dimensional map contained 117 antigens and 29 sera (117x29, table S2). In this dataset, 127 out of the 3393 (3.74%) titers were set to ‘NA’ due to the SD of log_2_ transformed HI titers of repeats being above 1.5. The resulting map geometry was close to identical to the three-dimensional map used in the dimensionality investigation above (Procrustes RMS of 0.67 AU, median Procrustes difference of 0.32 AU). The correlation between the distances obtained from the HI table and from the map was comparable to that previously observed in the dimensionality testing described above (R^2^ = 0.64) (fig. S3A). The point positions were further validated using the ‘moveTrappedPoints’ and ‘checkHemisphering’ functions (Racmacs). Here, each point is moved individually in the map to assess whether better (trapped points) or equal (hemisphering points) local optima, i.e. positions with a lower or equal point stress as compared to its original position respectively, are found upon relaxing of the map (optimization of point positions starting from their current coordinates). These analyses revealed that no points located in local optima, nor hemisphering points, were detected in this map.

We then investigated whether better maps, i.e. with lower total stress, could be generated using dimensional annealing, through the ‘make.acmap’ function (Racmacs) with the option of dim_annealing set to ‘TRUE’. Using this method, each optimization of the map is first optimized in five dimensions. The dimension is then reduced by one, the map is relaxed, and optimized again, until the set dimension is reached. The three-dimensional map generated using dimensional annealing had a higher total stress (4177) as compared to the map generated using the default settings (4151), indicating that dimensional annealing did not allow the generation of a better antigenic map.

The stability of the map was first assessed by investigating the effect of removing each single individual antigen and serum. The median Procrustes distance of all points was determined for each resulting map, and the distribution was plotted as histograms (fig. S3B, C). Generally, only minimal changes in the map positions were observed upon removing individual antigens or sera, with a mean median difference of 0.09 AU and 0.34 AU when removing individual antigens and sera, respectively. Removal of three individual antigens and five individual sera led to maps with median Procrustes distances above 0.40 AU and 0.60 AU, respectively (fig. S3B, C). These maps were compared to the full map in further detail (Data S4). In six out of eight maps, the difference was mainly the result of significant changes in the positions of clade 2.3.4.4 antigens and sera. In addition, large changes in positions were observed for several antigens located on the periphery of the antigenic map. Most notably, the position of the A/turkey/Wisconsin /1968 antigen (non-GsGd) changed substantially in four out of the eight maps. Taken together, this analysis indicated that the three-dimensional map geometry was generally robust and insensitive to the absence of individual antigens and sera.

To assess the certainty in point positions in the antigenic map, we used two tests. Firstly, triangulation blobs, indicating the area in which a particular point can be located without increasing the total map stress by more than one unit, were generated with the ‘triangulationBlobs’ function (Racmacs) with the option of ‘grid_spacing’ = 0.25. This analysis revealed that, while the overall geometry of the map was consistent, the positions of points at the periphery of the map were less well coordinated than those located in the center, which is expected given that peripheral points show overall lower HI reactivity and are not constrained by surrounding sera (Data S3C). Secondly, Bayesian bootstrapping was performed through the ‘bootstrapMap’ function in Racmacs for 1000 repeats with 100 optimizations per repeat, to understand the confidence of the positions of each point in the map. For each repeat, weights were randomly assigned to each titer in the HI data used to construct the map. For each point, the blob which encompassed the positions of 680 of the 1000 bootstrap runs (corresponding to a SD of 1) was computed using the ‘bootstrapBlobs’ function in Racmacs. The volumes of the resulting bootstrap blobs were calculated with a functionality available in PyRacmacs (*94*), by fitting a triangular mesh to each bootstrap point using the package trimesh (*95*) (code available via <https://github.com/epiv-lab/H5-antigenic-evolution> and Zenodo (*68*)). The radius of a sphere of equal volume, expressed in antigenic units (AU), was used as an indicator of the uncertainty of point position (fig. S3E, G). Generally, points which are well triangulated have spherical blobs, and the thus the reported radius gives a good estimate of their uncertainty. A good correspondence between blob maximum width and sphere radius was observed for most of the points, indicating that most blobs were indeed spherical (fig. S3F, H). Bootstrap blob volumes were generally larger for antigens located at the periphery of the antigenic map (Data S3F), in accordance with the result of the triangulation blob analysis. Generally, the largest bootstrap blobs were observed for the non-GsGd and clade 1 antigens, suggesting a higher degree of uncertainty in their positioning in the map.

Further analysis on the map stability was performed by comparing its individual optimizations. Upon generation of the antigenic map, 1000 optimizations were performed, resulting in 1000 maps which were subsequently sorted by ascending stress, i.e., optimization (opt.) 1 corresponds to the one with lowest stress. The positions of points and the total map stress of the best map (opt. 1) were compared to those from subsequent optimizations (fig. S4A, B, respectively). Conformations of the 600 best optimizations were generally similar to opt. 1, and differences in point positions indicated by the median Procrustes distance (fig. S4A) were the result of relatively big changes in position of only a few points. Interestingly, from optimization 602 onwards, a total of 125 maps with similar Procrustes distance distributions and total map stress were observed (fig. S4). Further investigation of these maps revealed that their conformations were virtually identical to one another, but significantly different from opt. 1. This alternative map conformation (Data S3D) represented the current data slightly less well than opt. 1 (R^2^ of the linear regression of HI table versus map distances of 0.6314 for opt. 602 as compared to 0.6362 for opt. 1). The main differences were found in positions of antigens at the periphery of the antigenic map, including the non-GsGd, clade 1 and clade 2.3.4.4 antigens map (Data S3E). Generally, collective changes in the positions of genetically similar antigens were observed, suggesting similar topology, especially for central points. This analysis showed that the map conformation was relatively stable, since most low stress maps resulted in similar conformations as the lowest stress map, albeit with the occasional movement of individual points. However, the observation of an alternative conformation with comparable total stress and HI table versus map distances suggested metastability of the antigenic map. Considering the relatively minor difference in total map stress (4151 for optimization 1 and 4279 for optimization 602), it should be noted that the addition of new data could potentially favor either of these conformations in the future.

Supplementary text S4 - Histopathology and immunohistochemistry in pre-clinical studies

In the pre-clinical ferret studies, tissues obtained 4 dpi were used for histopathological and immunohistochemistry (IHC) analyses. Generally, lesions in the respiratory tract were detected in all animals, but the extent differed between experimental groups and challenges. This first paragraph will describe the observed respiratory tract lesions qualitatively, and the following paragraphs the quantitative differences between animals from the different experimental groups.

In the lungs, the lesions were mostly associated with the bronchioles and bronchi and characterized by a mild to moderate thickening of the alveolar septa with infiltration of few neutrophils, lymphocytes, plasma cells as well as variable interstitial edema and epithelial necrosis. The alveolar lumina contained variable amounts of edema and increased numbers of alveolar macrophages and occasional neutrophils. The bronchioles and bronchi showed exocytosis of neutrophils and lymphocytes with epithelial hyperplasia and hypertrophy. There was perivascular and peribronchiolar/bronchial cuffing and edema as well as multifocal moderate type II hyperplasia. There was multifocal bronchoadenitis and bronchus associated lymphoid tissue (BALT) hyperplasia. In the trachea, there was mild exocytosis, and focal loss of ciliated cells with occasionally multifocal tracheal adenitis with necrosis and neutrophils. In the nose, the epithelium was flattened with loss of ciliated cells, and severe inflammation with exocytosis of neutrophils, and in the alveolar lumina many neutrophils, macrophages, cellular debris. In the lamina propria, there were variable numbers of neutrophils, lymphocytes, plasma cells and less macrophages.

The histological parameters that were scored are summarized in table S10. In animals from the Giza challenge study, significant differences in scoring were only observed between the Mock_VACC_ and vaccinated groups, and not between the vaccinated groups. A general pattern was observed where the median scoring for severity and extent of alveolitis, severity of bronchitis/bronchiolitis, peribronchial cuffing, alveolar edema, and hemorrhage was highest in animals from the Mock_VACC_ group, lower in those from the Anhui_VACC_ and AC-Anhui_VACC_ groups, and even lower in those from the Giza_VACC_ group (table S10, Fig. S11). Influenza virus nucleoprotein (NP) expression was detected in the alveoli, bronchioles, bronchi, and trachea of the the Mock_VACC_ animals, but absent in animals from the vaccinated groups. In the noses, NP antigen expression was detected in animals from all groups with few cells positive, with the highest score observed in the Mock_VACC_ (p<0.05 as compared to vaccinated groups).

In animals from the Sichuan challenge study, differences in histopathological scorings between groups were generally not statistically significant. Generally speaking, the severity and extent of alveolitis, severity of bronchitis/bronchiolitis, tracheitis, peribronchial cuffing, alveolar edema and hemorrhage was highest in animals from the Mock_VACC_ group, slightly less high in those from Anhui_VACC,_ group, and lower in those from the AC-Anhui_VACC_ and Sichuan_VACC_ groups, with Sichuan_VACC_ animals having slightly lower scores (table S10, fig. S11) . Interestingly, the sum of severity and extent of alveolitis was significanlty lower in animals from the AC-Anhui_VACC_ and Sichuan_VACC_ groups as compared to those from the Mock_VACC_ and the Anhui_VACC_ groups, which were not significantly different from one another (table S10, fig. S11). Scoring of NP antigen expression in the alveoli, bronchioles, bronchi, and trachea were highest in animals from the Mock_VACC_ group, lower in those from the Anhui_VACC_ group , minimal to absent in those from the AC-Anhui_VACC_ and Sichuan_VACC_ groups. In the noses, few cells were positive for NP antigen expression in all groups.

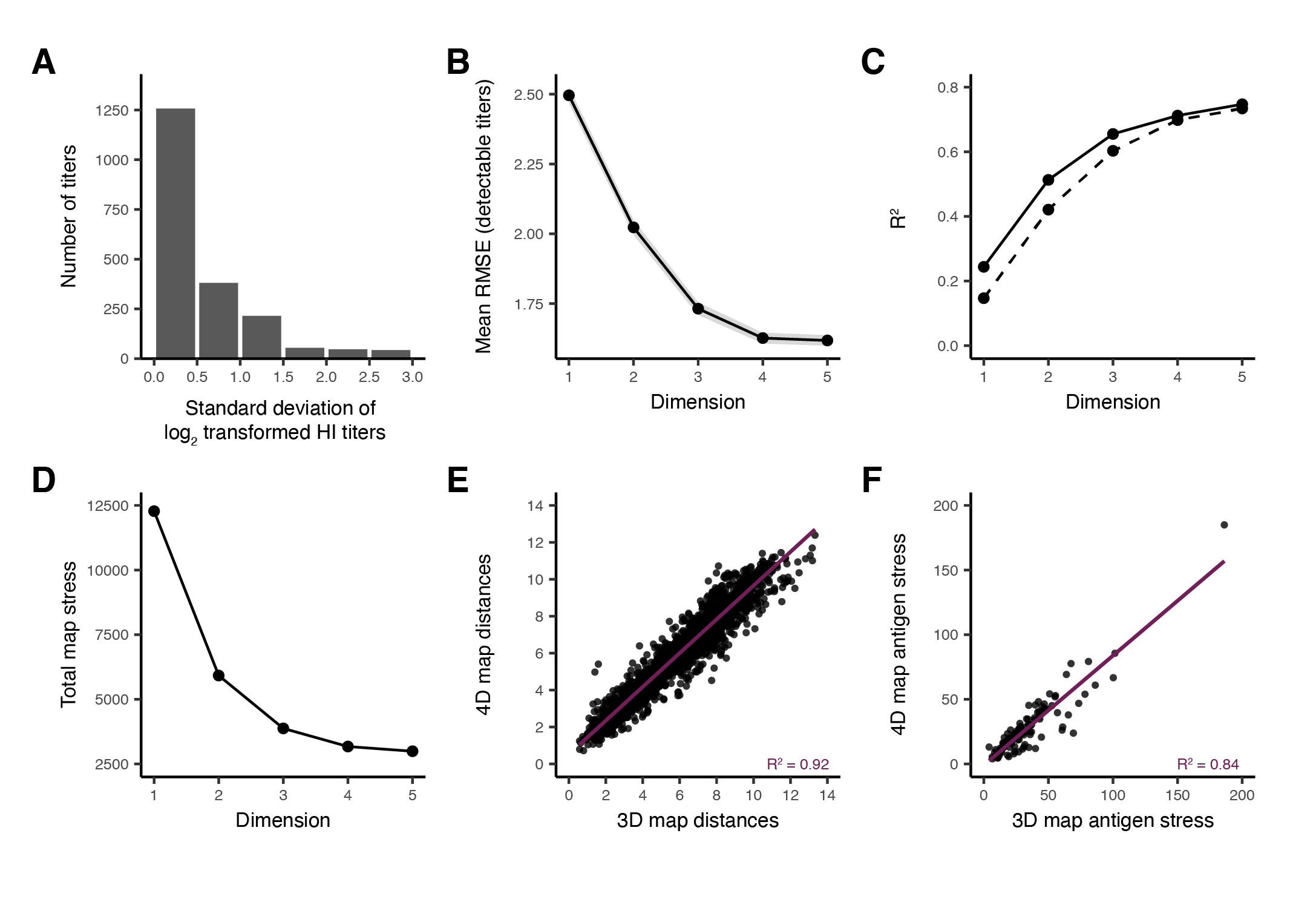

**Fig. S1.**

**Analyses of A(H5) HI titer dataset variability and map dimensionality.** (**A**) Histogram of standard deviation of log_2_ transformed HI titers between assays used to construct the full map dataset (127x33). (**B**) Dimensionality test, indicating the mean root mean square error (RMSE) between predicted and measured detectable HI titers when 10% of the titers was removed, using 1 to 5 dimensions. The grey shading indicates the 95% confidence interval of the RMSE. (**C**) Regression coefficient (R^2^) of linear regression between HI titers (table distances), and corresponding Euclidian distances in the antigenic maps generated in one to five dimensions. The R^2^ of the linear regressions based on all HI titers are shown as a solid line, and those including only detectable HI titers are shown as a dashed line. (**D**) Total map stress using 1 to 5 dimensions. (**E**) Scatter plots of pairwise antigen-serum map distances in the three- versus the four-dimensional map. (**F**) Scatter plots of individual antigen stress in the three- versus the four-dimensional map. In (E-F), linear regression lines are plotted, and the regression coefficient (R^2^) is indicated. Analyses in (B-F) are performed using the 114x28 dataset (table S2). HI: hemagglutination inhibition; 3D: three-dimensional; 4D: four-dimensional.

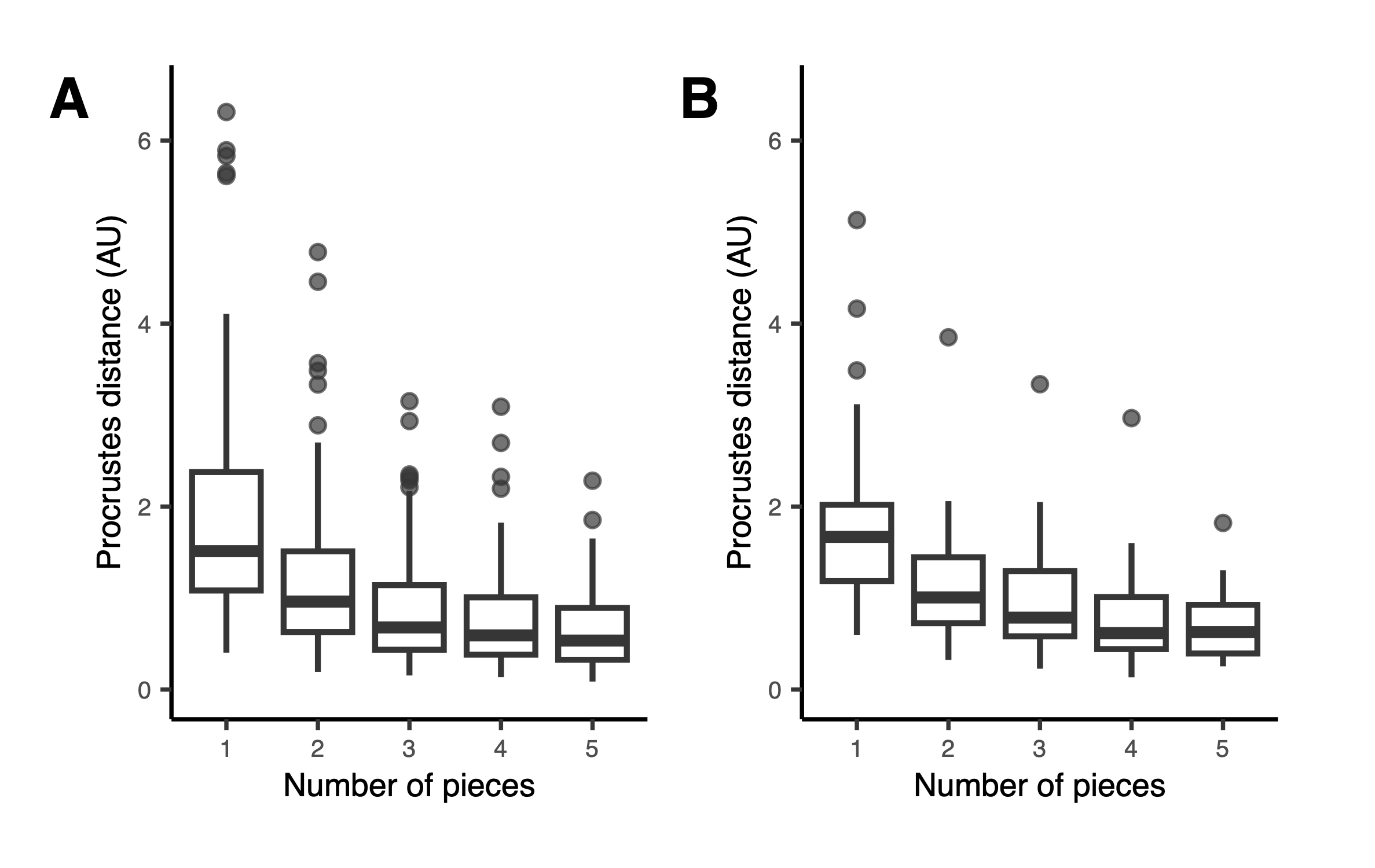

**Fig. S2.
Piecewise Procrustes analysis comparing the A(H5) antigenic maps in three and four dimensions.** Procrustes distances between the three- and four-dimensional maps using one to five pieces for the piecewise Procrustes analysis are displayed. Box plots summarize the Procrustes distance for individual antigens (**A**) and sera (**B**). The line indicates the median, the lower and upper hinges the first and third quartiles, respectively, and the upper and lower whiskers extend respectively to the largest and smallest values that fall within 1.5 times the inter-quartile range (distance between the first and third quartiles). Data points beyond the end of the whiskers are plotted individually. Analyses were performed using the 114x28 dataset (Table S2). AU: Antigenic units.

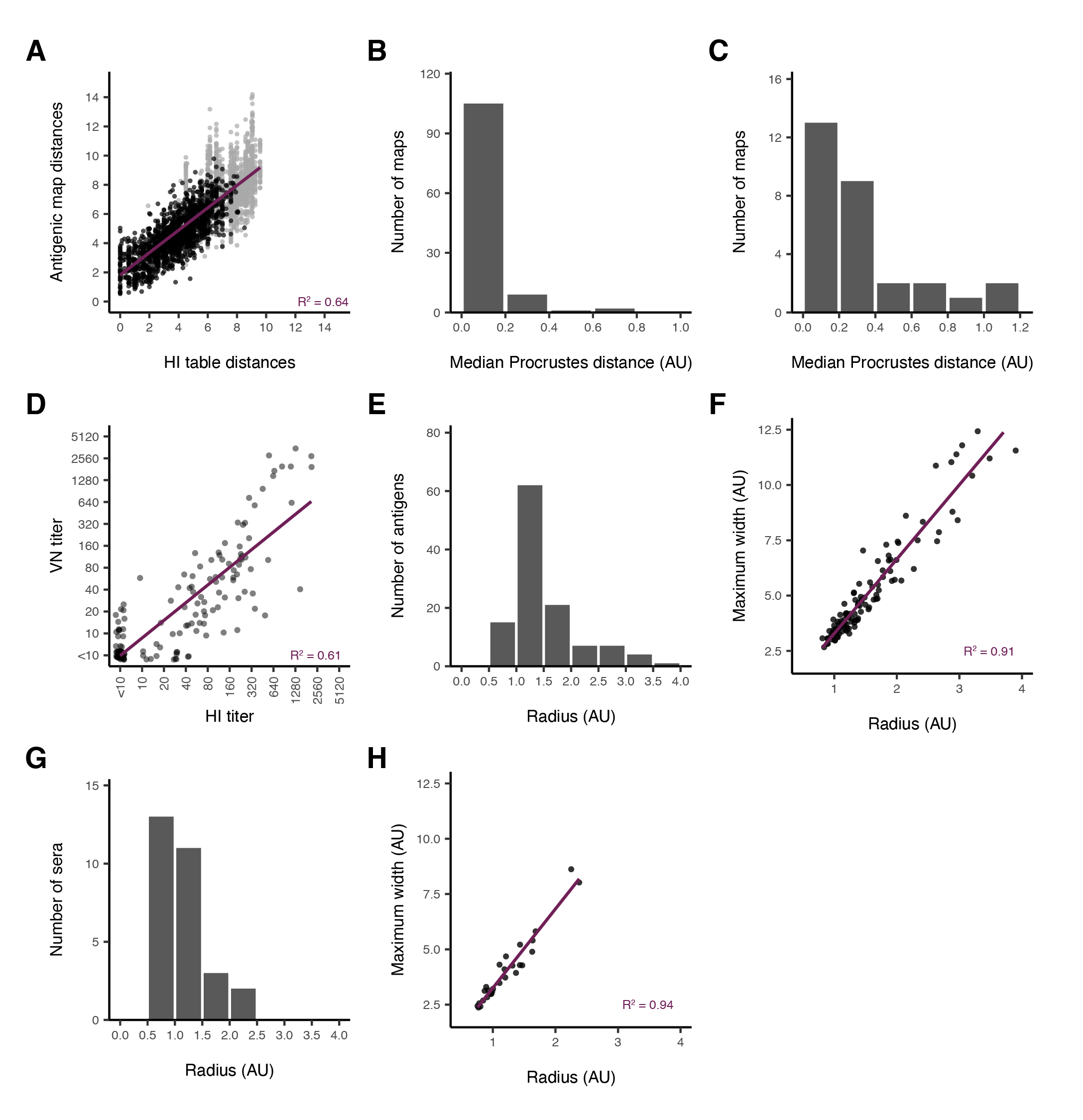

**Fig. S3.**

**Accuracy and stability of the three-dimensional A(H5) antigenic map.** Analyses were performed using the final 117x29 dataset (table S2). (**A**) Scatter plot of HI titer table distances versus the corresponding Euclidian distances in the antigenic map distances inferred from detectable titers (black) and from non-detectable titers (grey) are shown. (**B, C**) Histograms showing the distribution of the median Procrustes distance over individual maps as compared to the full antigenic map upon removing individual antigens (B) or sera (C) from the antigenic map. (**D**) Scatter plot of HI titers versus VN titers. (**E, G**) Histograms of the radius of Bayesian bootstrap blobs for antigens (E) and sera (G). (**F, H**) Comparison the maximum width of Bayesian bootstrap blobs versus the radius of equal-volume spheres for antigens (F) and sera (H). In (A, D, F, H), linear regression lines are plotted, and the regression coefficient (R^2^) is indicated. HI: hemagglutination inhibition; VN: Virus neutralization; AU: antigenic unit.

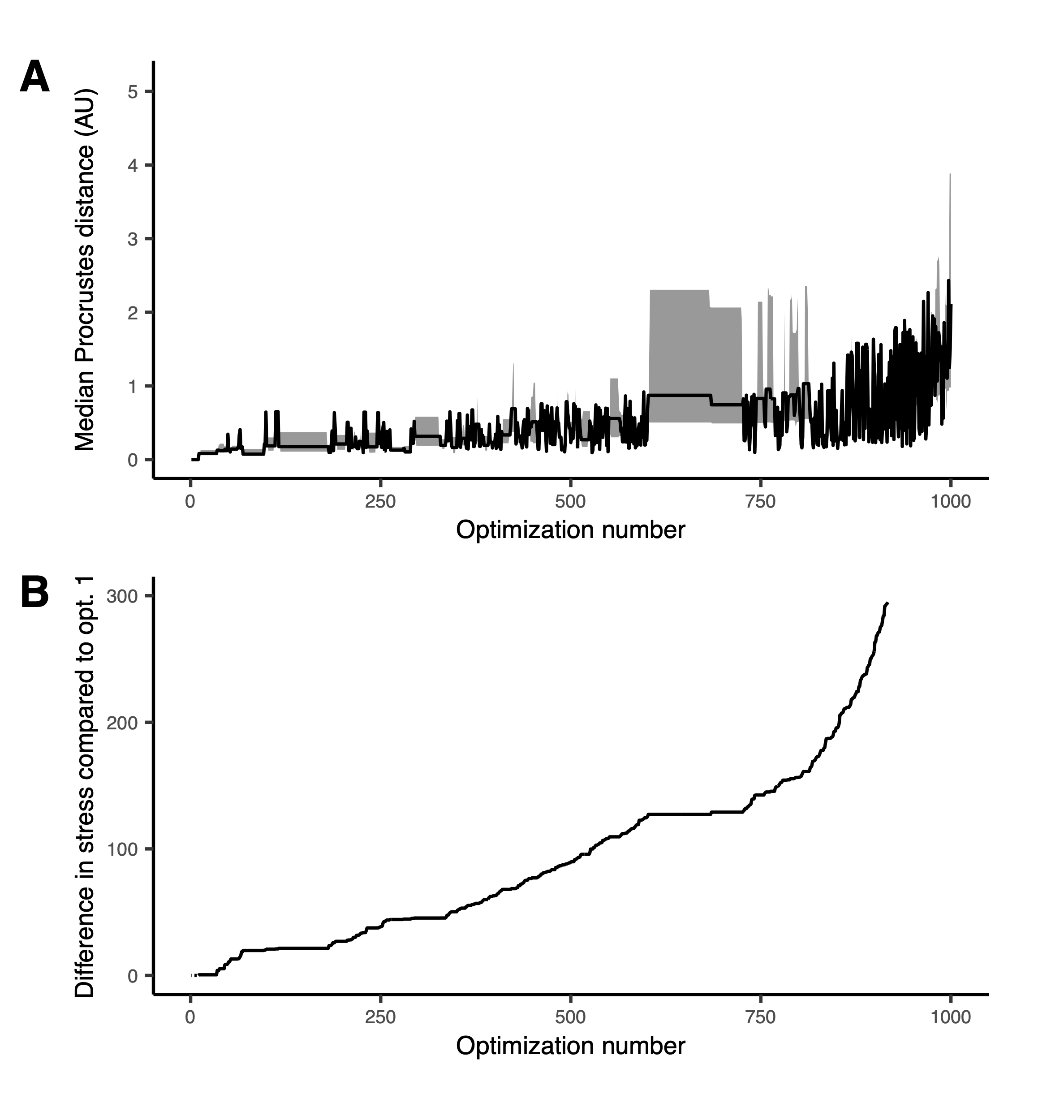

**Fig. S4.**

**Analyses of the A(H5) antigenic map stability across 1000 ranked optimizations.** Analyses were performed using the final 117x29 dataset (table S2). (**A**) Median Procrustes distance between the position of points in each optimization as compared to those in the lowest stress optimization (opt. 1) for ranked optimizations in increasing stress orders. Shown is the median Procrustes distance (black line), and the distribution of Procrustes distances excluding the highest and lowest 20% (grey shading). (**B**) Difference in total map stress as compared to the lowest stress optimization (opt. 1) for all 1000 optimizations. Values above 300 (beyond optimization 917) were not plotted to allow a detailed visualization of the values below 300. The maximum value observed was 1471 for optimization 1000. AU: Antigenic units; Opt.: optimization.

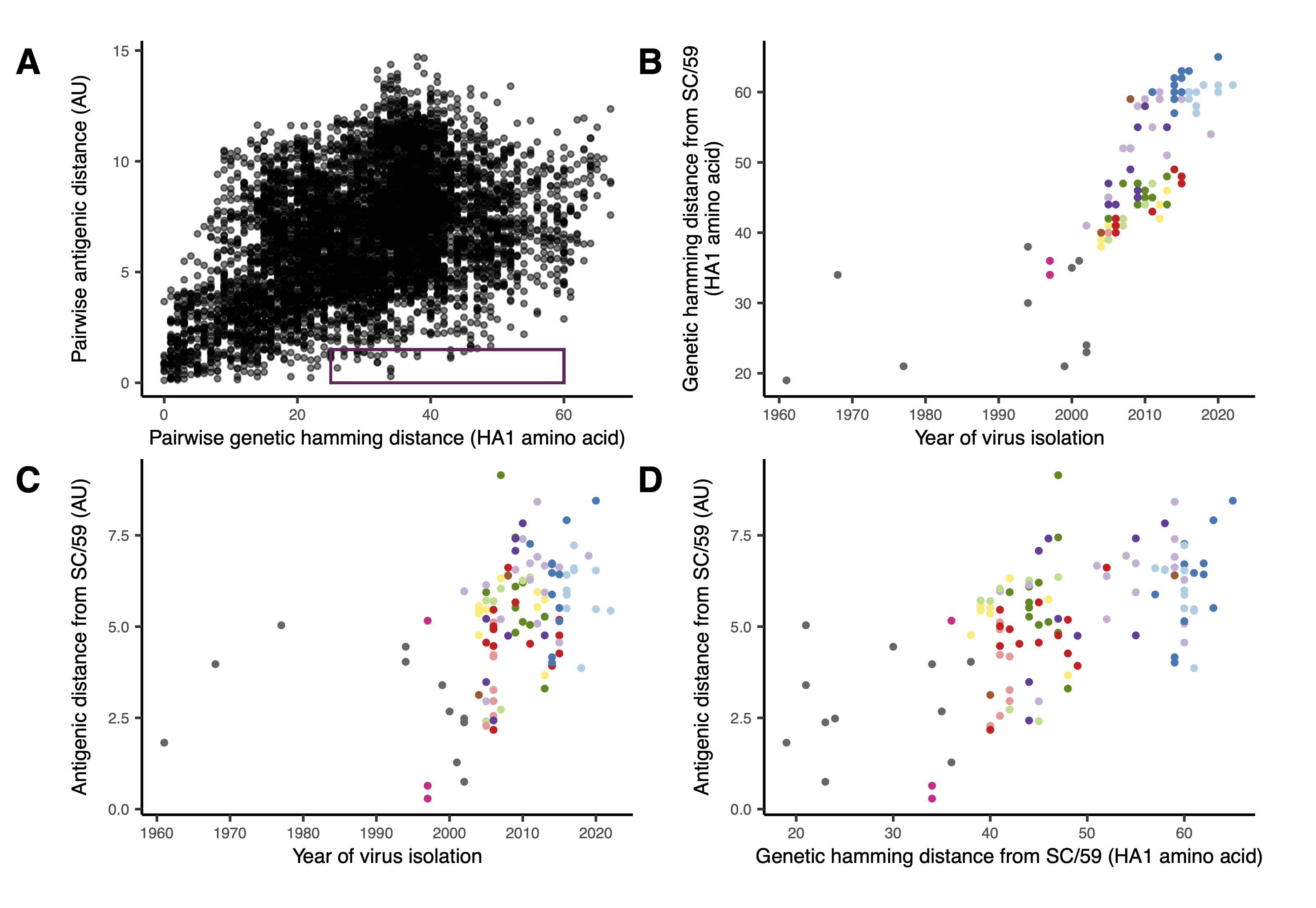
Fig. S5.­­­­­

**Genetic and antigenic diversity of antigens in the A(H5) antigenic map.** (**A**) Pairwise antigenic distances derived from the antigenic map as a function of pairwise genetic hamming distances (amino acid differences between two antigens) shown for all antigen-antigen pairs. The rectangle highlights the selected outlier antigen-antigen pairs with a genetic distance above 25 amino acids and an antigenic distance below 1.5 AU. Points are depicted in grey including a degree of transparency allowing the visualization of overlapping points. (**B**) Genetic hamming distance from A/chicken/Scotland/1959, first A(H5) virus ever isolated, as a function of the year of virus isolation, shown for each antigen in the map. (**C**) Antigenic distance from A/chicken/Scotland/1959 as a function of the year of virus isolation, shown for each antigen in the map. (**D**) Antigenic distance from A/chicken/Scotland/1959 as a function of genetic hamming distance from A/chicken/Scotland/1959, shown for each antigen in the map. In (B-D), the color of each point corresponds to the antigen’s genetic clade as indicated in Fig. 1B. AU: Antigenic units; SC/59: A/chicken/Scotland/1959.

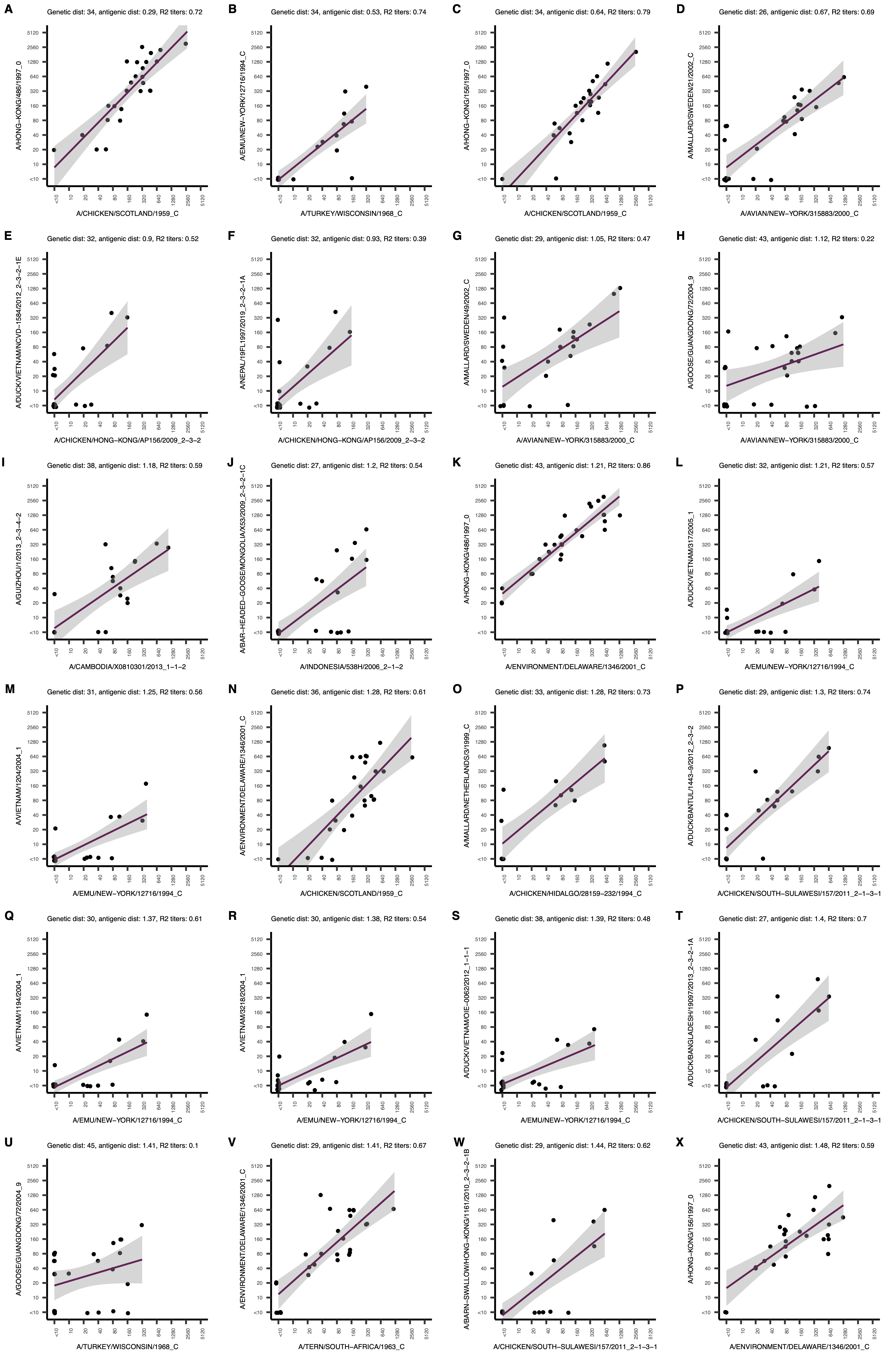

Fig. S6.

**Comparison of HI titers between antigen pairs with a small antigenic distance and a large genetic distance.** Antigen-antigen pairs with a genetic hamming distance above 25 HA1 amino acids and an antigenic distance below 1.5 AU were selected, as indicated in fig. S5A. Each panel (A-X) shows a scatter plot of the HI titers of the two antigens indicated along the axis. A linear regression line is plotted with the corresponding 95% confidence interval in grey. On top of each panel, the genetic and antigenic distance (dist.) between the two antigens as well as the R^2^ is indicated.

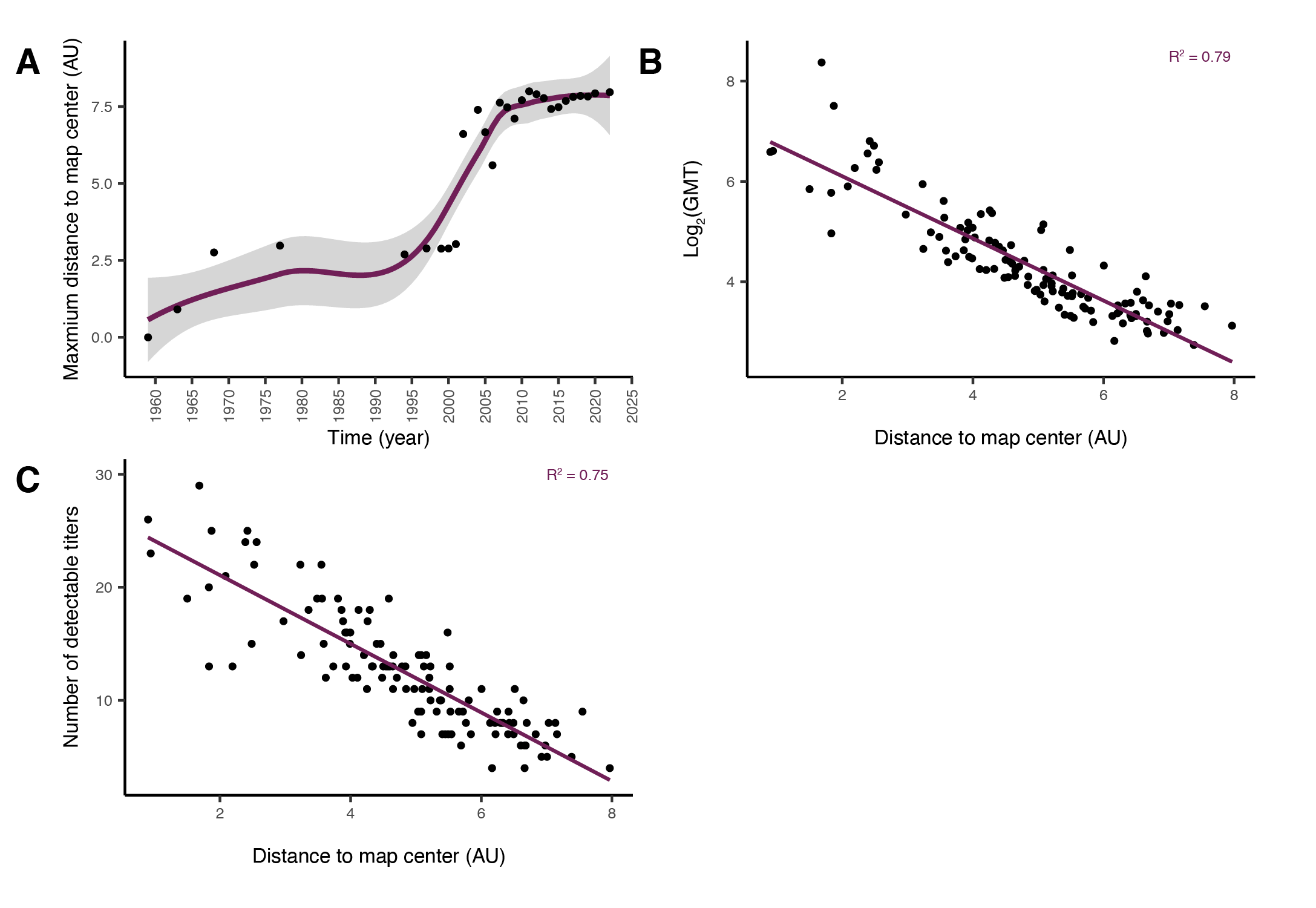
Fig. S7.

**Antigen distance to the center of the map.** The map center corresponds to the mean of all antigen position coordinates in each direction (x, y and z). (**A**) Maximum antigen distance from the map center over time. For each year in the dataset, the positions of all antigens from viruses isolated up until that year were used to compute the center of mass. The maximum distance of the respective antigens to the center was plotted. A smoothed local regression curve is plotted with the ‘geom_smooth’ function (ggplot2) using method = ‘loess’. The 95% confidence interval is displayed in grey. (**B**) Log_2_ transformed geometric mean titer as a function of the distance to the map center in antigenic units shown for each antigen in the map. (**C**) Number of detectable HI titers in the dataset as a function of the distance to the map center in antigenic units shown for all antigens in the map. In (C, D), linear regression lines are plotted, and the regression coefficient (R^2^) is indicated. AU: antigenic units; GMT: geometric mean titer.

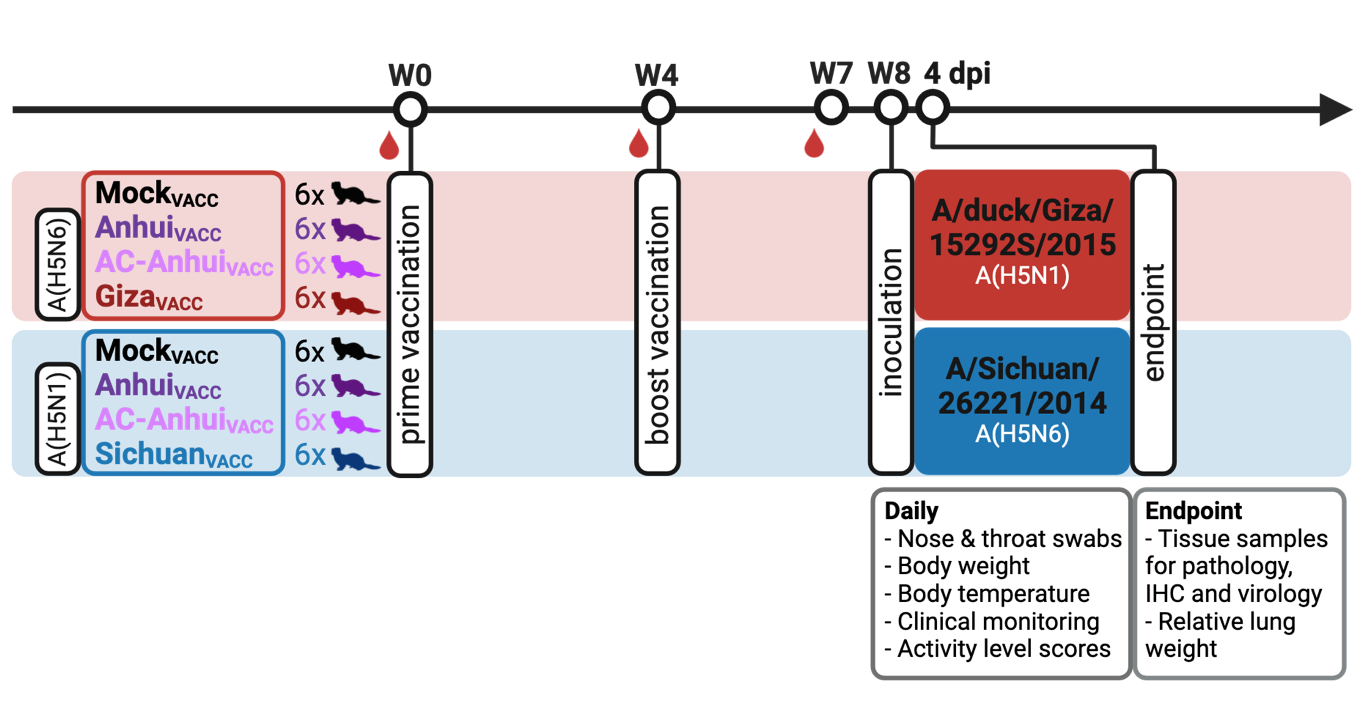

**Fig. S8.
Vaccination-challenge study design.** Schematic overview of the two vaccination-challenge experiments. The timeline in weeks (W) is indicated on top. Six ferrets per group were vaccinated twice with adjuvanted split-inactivated vaccines four weeks apart. Red drops denote serum sample collection before the first and second vaccination, as well as three weeks after the second vaccination. Four weeks after the boost vaccination, animals were inoculated either with A/duck/Giza/15292S/2015 virus (A(H5N1), clade 2.2.1.2) or with A/Sichuan/26221/2014 virus (A(H5N6), clade 2.3.4.4a). Daily, nose and throat swabs were collected, and body weight, body temperature, clinical signs, and activity level scores were monitored. Ferrets were sacrificed four days post-inocation, and relevant tissues were collected for virological and histopathological analysis. W: week; dpi: days post-inoculation; IHC: immunohistochemistry.

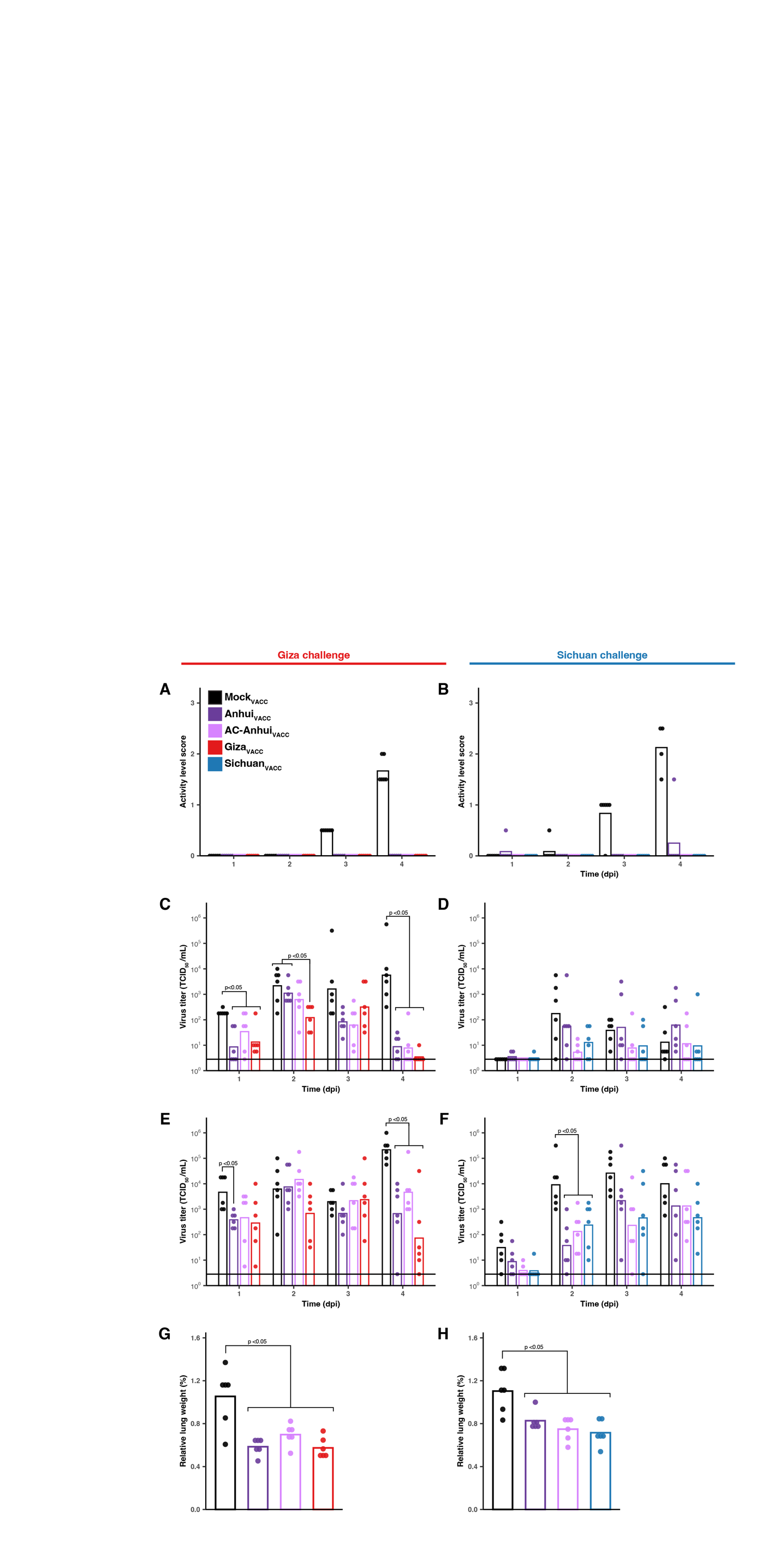
 **Fig. S9.
Inactivity scores, virus titers in respiratory swabs and­ relative lung weight of ferrets in vaccination-challenge studies.** Vaccination-challenge with the Giza virus (left panels), and the Sichuan virus (right panels). (**A** and **B**) Daily activity level scores determined as follows: 0 - alert and playful, 1- alert and playful only when stimulated, 2- alert but not playful when stimulated, 3- neither alert nor playful when stimulated. Bars represent the group mean (n=6), and dots represent the activity level score of individual animals. (**C**-**F**) Virus titers (TCID_50_/mL) in nose (C and D) and throat swabs (E and F) on a logarithmic scale. Bars represent the group geometric mean titer (n=6) and dots represent the titers in swabs of individual animals. The horizontal lines indicate the detection limit. (**G-H**) Relative lung weight, defined as the percentage of lung weight relative to total body weight at necropsy. Bars represent the mean per group (n=6), and dots represent the relative lung weight of individual animals. Statistically significant differences, determined with pairwise Mann-Whitney tests, are indicated with the corresponding p-value in panels (C-H). All graphs are color-coded according to the experimental groups, as indicated in the legend. Dpi: days post-inoculation; TCID_50_: 50% tissue culture infectious dose.

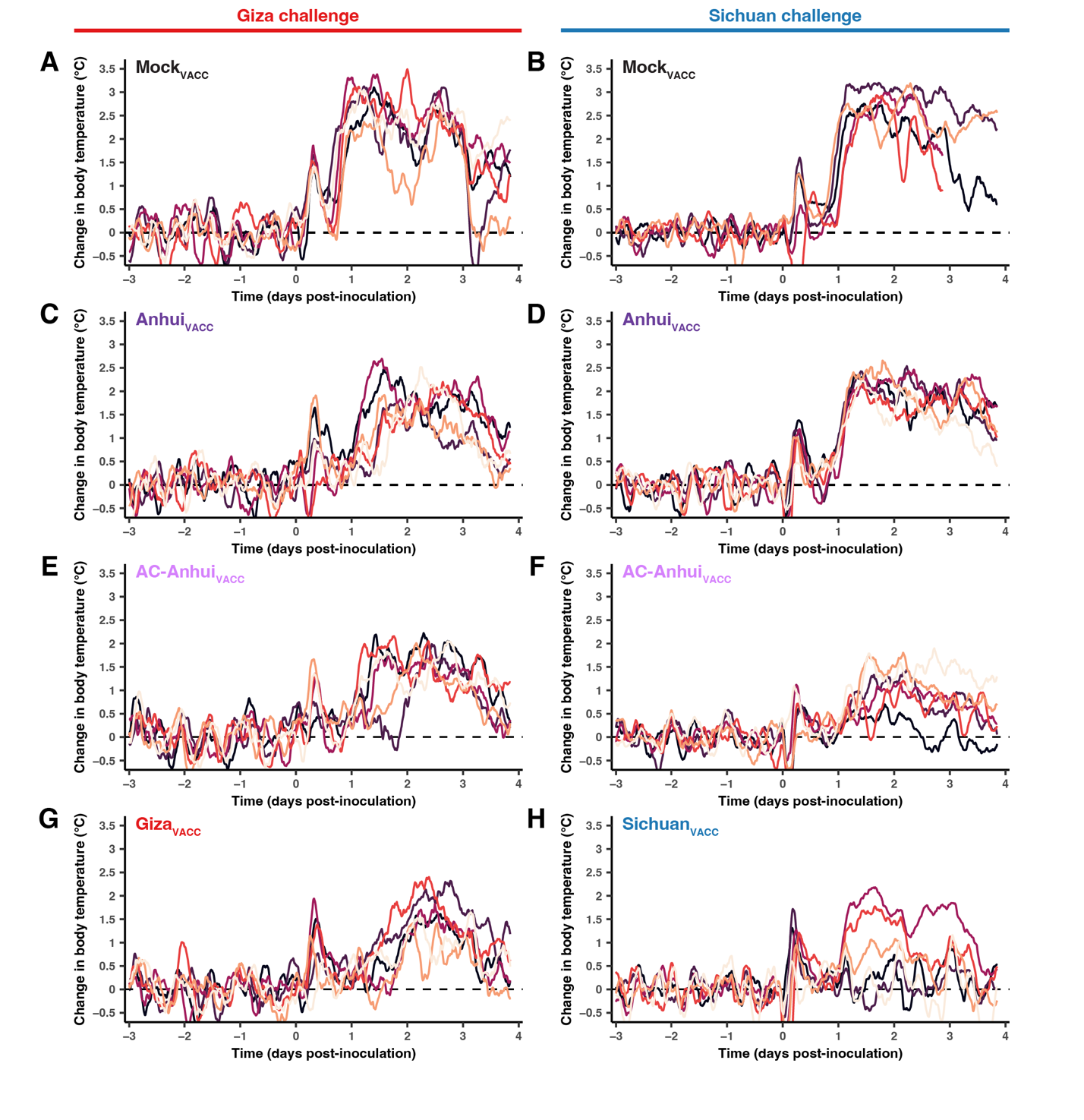
 **Fig. S10.
Body temperature change of individual ferrets in the vaccination-challenge studies.** Vaccination-challenge with the Giza virus (left panels): (**A**) Mock_VACC_, (**C**) Anhui_VACC_, (**E**) AC-Anhui_VACC_, and (**F**) Giza_VACC_ groups. Vaccination-challenge with the Sichuan virus (right panels): (**B**) Mock_VACC_, (**D**) Anhui_VACC_, (**F**) AC-Anhui_VACC_, and (**G**) Sichuan_VACC_. Body temperature change from baseline (mean body temperature recorded during the three days prior to inoculation, indicated as horizontal black line). Body temperature was recorded every ten minutes using probes surgically implanted in the peritoneal cavity. Four-hour sliding means are displayed. Each line represents the data of an individual animal (n=6). In the Sichuan challenge Mock_VACC_ group (B), data of five individual animals are shown due to a malfunctioning temperature probe, and data of the two deceased animals are included up until three days post-inoculation.

**­­­­**
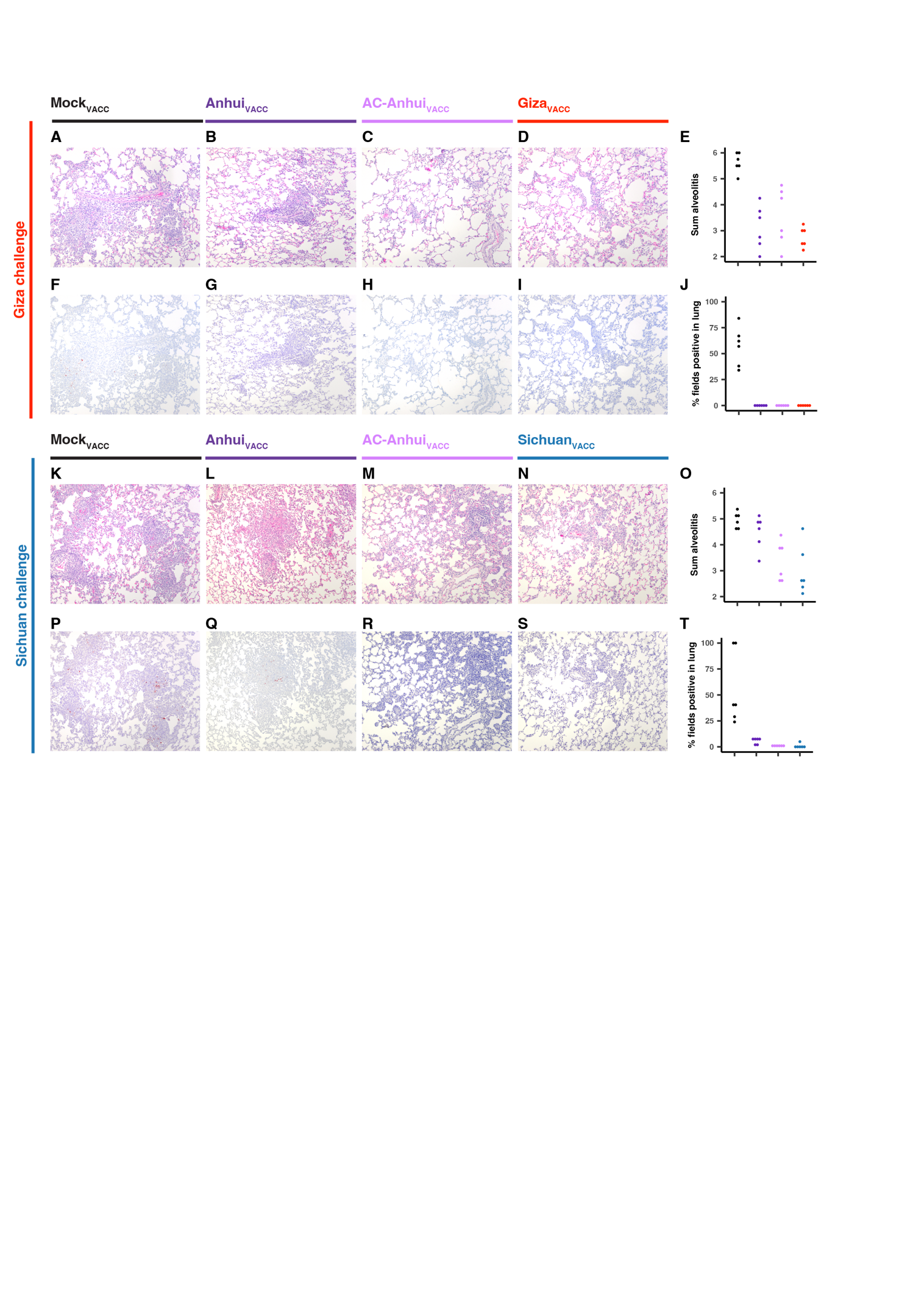

**Fig. S11.
Histopathology and immunohistochemistry of lung tissues from ferrets in the vaccination-challenge studies.** Representative images of formalin-fixed paraffin-embedded lung sections from animals with representative scores for histopathology and influenza virus nucleoprotein (NP) antigen expression for each experimental group are shown. (**A**-**D** and **K**-**N**) Lung sections stained with hematoxylin and eosin. (**E** and **O**) Sum of the scoring of severity and extent of alveolitis for each animal (n=6). Severity of alveolitis was scored as follows: 0: no inflammatory cells; 1: few inflammatory cells; 2: moderate number of inflammatory cells; 3: many inflammatory cells. Extend of alveolitis was scored as follows: 0: 0%; 1: 1–25%; 2: 25–50%; 3: >50%. (**F**-**I** and **P**-**S**) Immunohistochemical staining of lung sections with anti-influenza NP antibody. (**J** and **T**) Scoring of percentage of NP-positive fields for each animal (n=6). Cumulative scores for each animal are the percentage of fields positive for NP based on 25 arbitrarily chosen, 20x objective, fields of lung parenchyma. Vaccination-challenge studies and vaccine groups are indicated in the figure. Data are color-coded based on vaccine group as indicated in the figure.

Movie 1. A(H5) antigenic evolution over time. The antigens and sera of the three-dimensional antigenic map constructed from the final 117x29 dataset (as shown in Fig. 1B. and Data S2) appear based on the year of isolation of the respective virus, as indicated in the top-right corner. Antigens are displayed as closed spheres and sera are displayed as open cubes. Antigens and sera are color-coded based on the genetic HA clade, as indicated in the bottom-left legend. Each direction (x, y, z) represents antigenic distance, and one square of the grid corresponds to one antigenic unit, which is defined as a two-fold difference in HI titer. The antigenic map is oscillating for visualization purposes.

Table S1. Accession numbers and names of HA sequences for generation of the phylogenetic tree. For sequences obtained from the GISAID database (i.e. accession numbers starting with ‘EPI’), information on the contributors of the sequences is indicated.

Table S2. Antigens selected for antigenic characterization and sera production. Under ‘Antigen’, the names of the antigens as used in this study (virus name followed by the respective genetic clade, with ‘C’ for non-GsGd antigens) are listed. In the column ‘Homologous serum’, the names of the sera used in this study are indicated on the same row as the homologous antigen. Under ‘Accession number’, the accession numbers in the BV-BRC or GISAID (starting with ‘EPI’) databases are listed. The column ‘Present in dataset 114x28’ indicates whether the antigen was present in the dimension testing dataset, as detailed in supplementary text S2 and S3. The column ‘Present in dataset 117x29’ indicates whether an antigen was present in the final dataset. The column ‘Virus type’ indicates whether a full virus or a recombinant virus with the respective HA (without MBCS if applicable) in the background of PR/8 or PR/8 HY virus was used. The column ‘CVV-(like)’ indicates whether an antigen is a WHO candidate vaccine virus (CVV), or if it is the closest matching antigen in our dataset to a CVV. For all A/H5 WHO CVVs (*16, 17*) that were not present in our dataset, the genetically closest antigen in our dataset (full HA amino acid level excluding the signal peptide) was determined. If the difference between the CVV and the respective antigen was less than ten amino acids, these are labeled 'CVV-like' and the amino acid differences are indicated in the 'Notes' column. In ‘Notes’, further details are listed.

Table S3. Ferret sera used for antigenic characterization. Under ‘Virus’, the names of the virus used to generate the respective serum are indicated. Under ‘Genetic clade’, the respective HA genetic clade of the antigen is indicated. ‘Virus type’ indicates whether a full virus or a recombinant virus with the respective HA (without MBCS) in the background of PR/8 or PR/8 HY virus was used. Under ‘Boost’, a ‘+’ indicates the use of a subcutaneaous boost, and a ‘-‘ indicates that no boost was performed. Under ‘HA’ and ‘NA’, the names of the respective gene segments used are listed, and under ‘HA accession number’ and ‘NA accession number’, the corresponding database accession numbers are listed. ‘PR/8 backbone’ indicates whether PR/8 or PR/8 HY was used.

Table S4. Merged HI titer dataset used for generation of the A(H5) antigenic map. Rows correspond to antigens and columns to sera. Titers for which repeat titrations varied above the set limit (titers set to ‘NA’, see supplementary text S1) were excluded from the dataset and indicated with an Asterix (*).

Table S5. Pairwise distances between antigens in the A(H5) antigenic map. Distances were extracted from the antigenic map displayed in Fig. 1B and Data S2 and are expressed in antigenic units (AU).

Table S6. Resialylated turkey red blood cell assay with viruses carrying wild-type and engineered HAs. Data of individual assays are separated by thick horizontal lines. Undetectable titers (<0,5 HAU/25 µL) are denoted by “-“. TRBCs: turkey red blood cells; VCNA: *Vibrio cholerae* neuraminidase; α2,3: α2,3-sialyltransferase; α2,6: α2,6-sialyltransferase.

Table S7. HI reactivity of viruses carrying wild-type and engineered HAs. For viruses carrying wild-type HAs, merged data from the full H5 map dataset are shown. For the viruses carrying engineered HAs, data of a single representative HI assay are shown.

Table S8. HI dataset of vaccination sera. The first 7 columns contain the data of vaccination sera generated with whole-inactivated vaccines. The subsequent columns contain the data of sera from the vaccination-challenge studies. Titers which were not measured are indicated with an Asterix (*). Data of a single HI assay are shown.

Table S9. HI titer dataset of vaccination sera from the vaccination-challenge studies. HI titers of sera obtained pre- and post- boost vaccination against antigens used in the vaccination-challenge studies.

**Table. S10. Summary of histopathological and immunohistochemistry scoring in the vaccination-challenge studies.** The median is indicated followed by the range in brackets. The severity of alveolitis, bronchiolitis, bronchitis, bronchial adenitis, tracheitis, and rhinitis was scored as follows: 0: no inflammatory cells; 1: few inflammatory cells; 2: moderate numbers of inflammatory cells; 3: many inflammatory cells. The extend of alveolitis was scored as follows: 0: 0%; 1: 1–25%; 2: 25–50%; 3: >50%. The presence of alveolar edema, alveolar hemorrhage, and type II pneumocyte hyperplasia was scored as follows: 0: no; 1: yes. The extent of peribronchial, peribronchiolar, and perivascular infiltrates was scored as follows: 0: none; 1: one to two cells thick; 2: three to ten cells thick; 3: more than ten cells thick. For the lung, cumulative scores for each animal are the percentage of fields positive for influenza virus antigen based on 25 arbitrarily chosen, 20x objective, fields of lung parenchyma. For the bronchioles and bronchi in the lung, the main bronchus, trachea and nose, the percentage of positively staining epithelium was estimated for each slide and averaged per animal. Significant differences between groups (p<0.05 as determined with pairwise Mann-Whitney tests) are indicated as follows: ‘*’ indicates a significant difference between Mock_VACC_ and vaccinated groups, ‘†’ indicates a significant difference between Mock_VACC_ and Giza_VACC_, ‘‡’ indicates a significant difference between both Mock_VACC_ and Anhui_VACC_ as compared to both AC-Anhui_VACC_ and Sichuan_VACC_, and ‘§’ indicates a significant difference between all vaccinated groups but not between AC-Anhui_VACC_ and Sichuan_VACC_.A ‘¶’ indicates that for one animals, the data was not present.

**Table S11. Non-coding regions of all gene segments of viruses used for the vaccination-challenge studies.** Listed are the virus name, the respective gene segment, the 3’ non-coding regions, including the start codon, and the 5’ non-coding regions, including the stop codon.

**Data S1.**

**A(H5) HA maximum likelihood phylogenetic tree.**

Zoomable pdf version of the tree displayed in Fig. 1A. Visualization as described for Fig. 1A, and in addition, the isolate names corresponding the HA sequences are shown. Available via <https://epiv-lab.github.io/H5-antigenically-central-vaccine/Data_S1.pdf>.

****Data S2.****

****Three-dimensional A(H5) influenza antigenic map.****
An interactive version of the three-dimensional antigenic map constructed from the final 117x29 dataset, shown in Fig. 1B. Antigens are displayed as closed spheres, and sera are displayed as open cubes. Antigens and sera are color-coded based on the genetic HA clade, as indicated on the right-hand side of the figure. Antigens and sera names can be visualized by hovering over the points. Each direction (x, y, z) represents antigenic distance, and one square of the grid corresponds to one antigenic unit, which is defined as a two-fold difference in HI titer. The antigenic map can be rotated by clicking and dragging in the panel. On the top right are different functions to explore the map, and a brief description of each function appears when hovering over. The total map stress, mean stress per titer and mean stress per detectable titer are indicated at the bottom left. Available via <https://epiv-lab.github.io/H5-antigenically-central-vaccine/Data_S2.html>.

**Data S3.**
**Validation of the A(H5) antigenic map.**
Interactive versions of the three-dimensional antigenic map, represented as described for Data S2. (**A**, **B**) Piecewise Procrustes analysis (see fig. S2 and detailed in the supplementary text S3) comparing the antigenic maps in three and four dimensions. The results of analysis with one (A) and two (B) pieces are displayed. The antigen color hue indicated which piece it belongs to, and the shading indicates the Procrustes distance according to the gradient displayed on the right, in antigenic units (AU). (**C**) Triangulation blobs indicating the area in which each datapoint can be located in the antigenic map without increasing the total map stress by more than one unit. (**D**) Alternative antigenic map conformation found upon comparing all 1000 optimizations (see fig. S4 and supplementary text S3). The map from optimization 602 is shown. (**E**) The lowest stress antigenic map (optimization 1), with Procrustes arrows pointing towards the positions of each antigen and serum in the optimization 602 map conformation. (**F**) Bayesian bootstrap blob size analysis. The antigen color corresponds to the radius (AU) of a sphere of equal volume than each blob as displayed on the right. For interpretation, 1-2 AU differences correspond to the HI assay variation. Available via <https://epiv-lab.github.io/H5-antigenically-central-vaccine/Data_S3.html>.

**Data S4.**
**The effect of removing single individual antigens and sera on the map geometry.**

**Each antigen and serum were individually removed from the antigenic map, and the full antigenic map was compared to the resulting maps, as detailed in the supplementary text S3. The maps with the highest median Procrustes distance are displayed.** (A-H) Interactive versions of the antigenic map, represented as described for Data S2. In each panel, the full antigenic map is displayed (117x29), and Procrustes arrows point at the positions of each antigen and serum in the map in which a single individual antigen (**A**-**C**) or serum (**D**-**H**) was removed, as indicated above each panel. The removed point is faded out, and no Procrustes is drawn. Ag.: Antigen; Sr.: Serum. Available via <https://epiv-lab.github.io/H5-antigenically-central-vaccine/Data_S4.html>.

**Data S5.**
**A(H5) antigenic maps highlighting WHO candidate virus vaccines and antigens used in the ferret vaccination studies.**
Interactive versions of the antigenic map (117x29), represented as described for Data S2. Sera are not shown, and antigens of interest are highlighted as opaque spheres. (**A**) Highlighting the WHO candidate virus vaccines (larger spheres) and the WHO CVV-like (smaller spheres) antigens (see table S2). (**B**) Highlighting antigens used in the vaccination-challenge studies highlighted as larger spheres. The antigenic maps can be rotated by clicking and dragging in the panel, and scrolling allows zooming in and out. Available via <https://epiv-lab.github.io/H5-antigenically-central-vaccine/Data_S5.html>.

**Data S6.**
**Merged antibody profiles upon vaccination with whole-inactivated vaccines containing engineered HA antigens.** An interactive version of the antibody profiles displayed in Fig. 2. For each HA vaccine antigen, the position, breadth, and height of a mean merged serum per group (n=6) are represented in the antigenic map from Fig. 1B. HA present in vaccine: (**A**) Iraq_VACC_, (**B**) CVA-Vietnam_VACC_, (**C**) CVA-Indonesia_VACC_, (**D**) CVA-Anhui_VACC_. Representation is as described for Fig. 2. In addition, the map orientation can be changed by clicking and dragging within the visualization, and scrolling allows zooming in and out. Antigen names can be visualized by hovering over the points. GMT: geometric mean titer. Available via <https://epiv-lab.github.io/H5-antigenically-central-vaccine/Data_S6.html>.

**Data S7.**
**Individual antibody profiles upon vaccination with whole-inactivated vaccines containing engineered HA antigens.**
Individual animal data used to generate merged antibody profiles displayed in Fig. 2 and Data S6. For each HA vaccine antigen, the position, breadth, and height of individual sera are represented in the antigenic map from Fig. 1B. HA present in vaccine: (**A**, **B**) Iraq_VACC_, (**C**, **D**) CVA-Vietnam_VACC_, and (**E**, **F**) CVA-Indonesia_VACC_. Using the same representation as Data S6. GMT: geometric mean titer. Available via <https://epiv-lab.github.io/H5-antigenically-central-vaccine/Data_S7.html>.

**Data S8.**
**Merged antibody profiles of upon vaccination with split-inactivated vaccines containing wild-type HA antigens or the antigenically central HA antigen.**
An interactive version of the antibody profiles displayed in Fig. 3. For each group, the position, breadth and height of a mean merged serum per group (n=6) are represented in the antigenic map from Fig. 1B. (**A**-**C**) Immune responses upon vaccination with A(H5N6) split-inactivated vaccines in the Giza challenge study or (**D**-**F**) A(H5N1) split-inactivated vaccines in the Sichuan challenge study. HA antigen present in vaccine: (**A, D**) Anhui_VACC_, (**B, E)** AC-Anhui_VACC_, (**C**) Giza_VACC_, and (**F**) Sichuan_VACC_. Using the same representation as Data S6. GMT: geometric mean titer. Available via <https://epiv-lab.github.io/H5-antigenically-central-vaccine/Data_S8.html>.

**Data S9.**
**Individual antibody profiles of animals from the Giza vaccination-challenge study.**
Individual immune responses upon vaccination with A(H5N6) split-inactivated vaccines in the Giza challenge study. Individual animal data used to generate merged antibody profiles displayed in Fig. 3 and Data S8. For each HA vaccine antigen, the position, breadth, and height of individual sera are represented in the antigenic map from Fig. 1B. HA antigen present in vaccine: (**A**-**F**) Anhui_VACC_, (**G**-**L**) AC-Anhui_VACC_, and (**M**-**R**) Giza_VACC_. Using the same representation as Data S6. GMT: geometric mean titer. Available via <https://epiv-lab.github.io/H5-antigenically-central-vaccine/Data_S9.html>.

**Data S10.**
**Individual antibody profiles of animals from the Sichuan vaccination-challenge study.**
Individual immune responses upon vaccination with A(H5N1) split-inactivated vaccines in the Sichuan challenge study. Individual animal data used to generate the merged antibody profiles displayed in Fig. 3 and Data S8. For each HA vaccine antigen, the position, breadth, and height of individual sera are represented in the antigenic map from Fig. 1B. HA antigen present in vaccine: (**A**-**F**) Anhui_VACC_, (**G**-**L**) AC-Anhui_VACC_, and (**M**-**R**) Sichuan_VACC_. Using the same representation as Data S6. GMT: geometric mean titer. Available via <https://epiv-lab.github.io/H5-antigenically-central-vaccine/Data_S10.html>.
