## Supplementary figures and images for "A vaccine antigen central in influenza A(H5) virus antigenic space confers subtype-wide immunity"

### Data S1

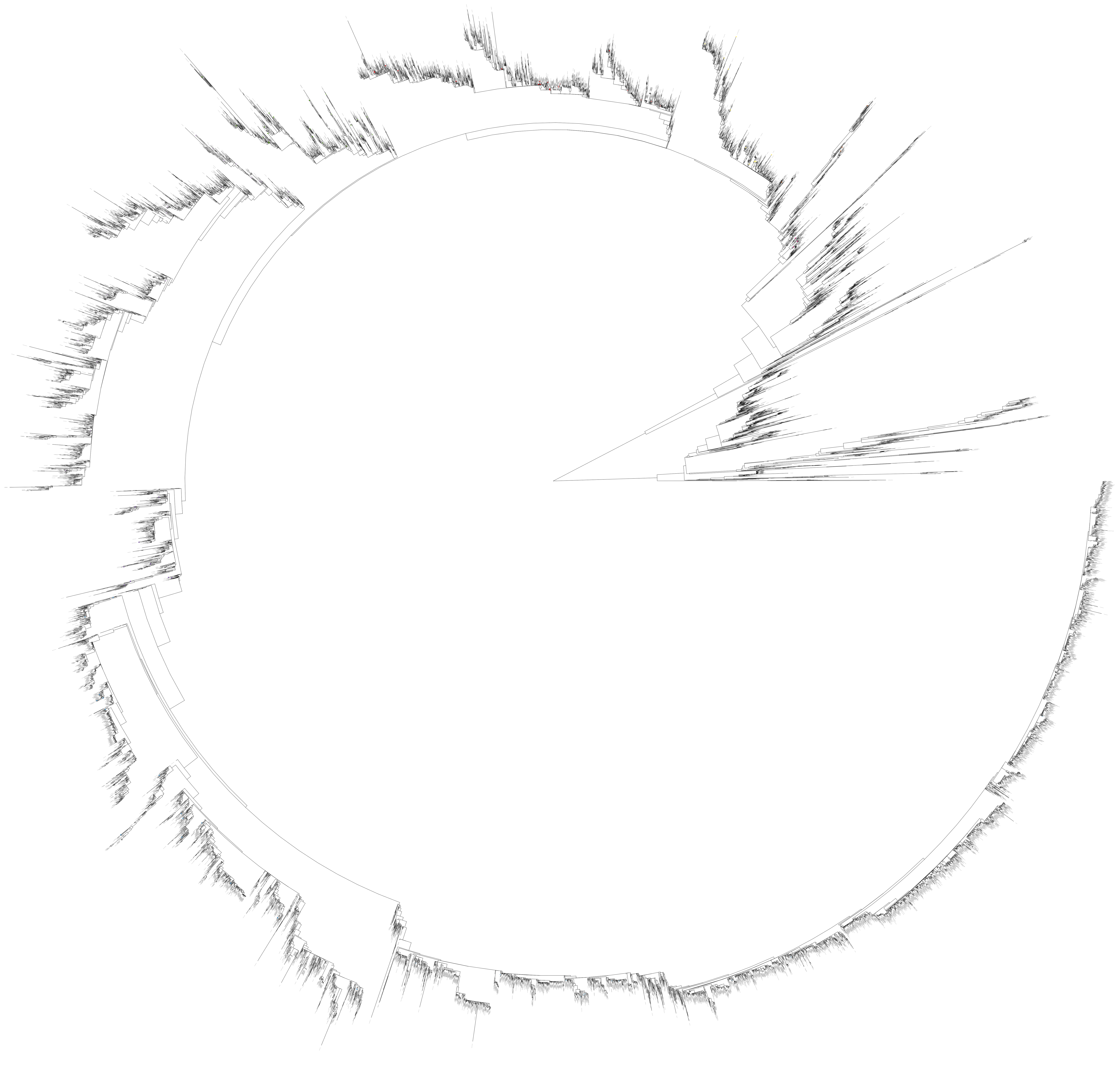
