## Supplementary material for "A vaccine antigen central in influenza A(H5) virus antigenic space confers subtype-wide immunity": Data S4: Data_S4.html


Data S4

### Row

#### A. Ag: A/HONG-KONG/486/1997\_0

#### B. Ag: A/TURKEY/65596/2006\_2-2

#### C. Ag: A/GUANGZHOU/39715/2014\_2-3-4-4E

#### D. Sr: A/HONGKONG/483B/1997

### Row

#### E. Sr: A/GUANGZHOU/39715A/2014

#### F. Sr: A/CHICKEN/JIANGSU/K0101B/2010

#### G. Sr: A/DUCK/GIZA/15292SA/2015

#### H. Sr: A/CHICKEN/VIETNAM/NCVD-15A59A/2015

### Row

**Data S4. The effect of removing single individual antigens
and sera on the map geometry.** Each antigen and serum were
individually removed from the antigenic map, and the full antigenic map
was compared to the resulting maps, as detailed in the supplementary
text. The maps with the highest median Procrustes distance are
displayed. (**A**-**H**) Interactive versions
of the antigenic map, represented as described for Data S2. In each
panel, the full antigenic map is displayed (117x29), and Procrustes
arrows point at the positions of each antigen and serum in the map in
which a single individual antigen (A-C) or serum (D-H) was removed, as
indicated above each panel. The removed point is faded out, and no
Procrustes is drawn. Ag.: Antigen; Sr.: Serum.
