## Supplementary material for "A vaccine antigen central in influenza A(H5) virus antigenic space confers subtype-wide immunity": Data S7: Data_S7.html


Data S7

### Row

#### A. Iraq_VACC_, I

#### B. Iraq_VACC_, II

#### C. VC-Vietnam_VACC_, I

### Row

#### D. VC-Vietnam_VACC_, II

#### E. VC-Indonesia_VACC_, I

#### F. VC-Indonesia_VACC_, II

### Row

**Data S7. Individual antibody profiles upon vaccination with
whole-inactivated vaccines containing engineered HA antigens.** 
Individual animal data used to generate merged antibody profiles
displayed in Fig. 2 and Data S6. For each HA vaccine antigen, the
position, breadth, and height of individual sera are represented in the
antigenic map from Fig. 1B. HA present in vaccine: (**A**,
**B**) Iraq_VACC_, (**C**,
**D**) CVA-Vietnam_VACC_, and (**E**,
**F**) CVA-Indonesia_VACC_. Using the same
representation as Data S6. GMT: geometric mean titer.
