## Supplementary material for "A vaccine antigen central in influenza A(H5) virus antigenic space confers subtype-wide immunity": Data S8: Data_S8.html


Data S8

### Row

#### A. Giza challenge, Anhui_VACC_

3.51 AU to center | GMT: 9 | 27 detectable titers

#### B. Giza challenge, AC-Anhui_VACC_

2.27 AU to center | GMT: 27 | 69 detectable titers

#### C. Giza challenge, Giza_VACC_

2.78 AU to center | GMT: 16 | 48 detectable titers

### Row

#### D. Sichuan challenge, Anhui_VACC_

3.60 AU to center | GMT: 6 | 7 detectable titers

#### E. Sichuan challenge, AC-Anhui_VACC_

1.73 AU to center | GMT: 10 | 38 detectable titers

#### F. Sichuan challenge, Sichuan_VACC_

4.23 AU to center | GMT: 7 | 21 detectable titers

### Row

**Data S8. Merged antibody profiles of upon vaccination with
split-inactivated vaccines containing wild-type HA antigens or the
antigenically central HA antigen.**  An interactive version of
the antibody profiles displayed in Fig. 3. For each group, the position,
breadth and height of a mean merged serum per group (n=6) are
represented in the antigenic map from Fig. 1B.
(**A**-**C**) Immune responses upon
vaccination with A(H5N6) split-inactivated vaccines in the Giza
challenge study or (**D**-**F**) A(H5N1)
split-inactivated vaccines in the Sichuan challenge study. HA antigen
present in vaccine: (A, D) Anhui_VACC_, (B, E)
AC-Anhui_VACC_, (C) Giza_VACC_, and (F)
Sichuan_VACC_. Using the same representation as Data S6. GMT:
geometric mean titer.
