## Supplementary material for "A vaccine antigen central in influenza A(H5) virus antigenic space confers subtype-wide immunity": Data S10: Data_S10.html


Data S10

### Row

#### A. Sichuan challenge, Anhui_VACC_, I

#### B. Sichuan challenge, Anhui_VACC_, II

#### C. Sichuan challenge, Anhui_VACC_, III

#### D. Sichuan challenge, Anhui_VACC_, IV

#### E. Sichuan challenge, Anhui_VACC_, V

#### F. Sichuan challenge, Anhui_VACC_, VI

### Row

#### G. Sichuan challenge, AC-Anhui_VACC_, I

#### H. Sichuan challenge, AC-Anhui_VACC_, II

#### I. Sichuan challenge, AC-Anhui_VACC_, III

#### J. Sichuan challenge, AC-Anhui_VACC_, IV

#### K. Sichuan challenge, AC-Anhui_VACC_, V

#### L. Sichuan challenge, AC-Anhui_VACC_, VI

### Row

#### M. Sichuan challenge, Sichuan_VACC_, I

#### N. Sichuan challenge, Sichuan_VACC_, II

#### O. Sichuan challenge, Sichuan_VACC_, III

#### P. Sichuan challenge, Sichuan_VACC_, IV

#### Q. Sichuan challenge, Sichuan_VACC_, V

#### R. Sichuan challenge, Sichuan_VACC_, VI

### Row

**Data S10. Individual antibody profiles of animals from the
Sichuan vaccination-challenge study.** Individual immune
responses upon vaccination with A(H5N1) split-inactivated vaccines in
the Sichuan challenge study. Individual animal data used to generate the
merged antibody profiles displayed in Fig. 3 and Data S8. For each HA
vaccine antigen, the position, breadth, and height of individual sera
are represented in the antigenic map from Fig. 1B. HA antigen present in
vaccine: (**A**-**F**) Anhui_VACC_,
(**G**-**L**) AC-Anhui_VACC_, and
(**M**-**R**) Sichuan_VACC_. Using
the same representation as Data S6. GMT: geometric mean titer.
